# Changing the initiation unit of nonribosomal peptide synthetases to access underexplored biosynthetic potential

**DOI:** 10.1101/2025.05.29.656725

**Authors:** Xianping Bai, Lin Zhong, Yang Liu, Hanna Chen, Xingxing Shi, Xingyan Wang, Qingsheng Yang, Xiaotong Diao, Dalei Wu, Youming Zhang, Xiaoying Bian

**Affiliations:** Helmholtz International Lab for Anti-Infectives, Shandong University-Helmholtz Institute of Biotechnology, State Key Laboratory of Microbial Technology, Shandong University, Qingdao, Shandong, 266237, China; CAS Key Laboratory of Quantitative Engineering Biology, Shenzhen Institute of Synthetic Biology, Shenzhen Institute of Advanced Technology, Chinese Academy of Sciences, Shenzhen 518055, China; College of Pharmacy, Linyi University, Shuangling Road, Linyi, 276000, China

**Keywords:** peptide, nonribosomal peptide synthetase, initiation module, starter condensation domain, genome mining

## Abstract

Nonribosomal peptide synthases (NRPSs) are large multimodular enzymes, capable of synthesizing nonribosomal peptides (NRPs) with diverse structures and bioactivities. Genome sequencing revealed a large number of uncharacterized NRPS biosynthetic gene clusters (BGCs) and their products are underexplored. The majority of NRPSs remain silent potentially attributed to factors such as the low activity of the initiation unit or insufficient precursor supply. Exchanging the starter condensation (Cs) domains within initiation unit can change the length of acyl chains of NRPs, hinting at a promising strategy through swapping of a well-studied Cs domain to activate the initiation unit and harness primary metabolites as precursors, which may offer a new option to access silent BGCs. Here, we first pinpointed two highly efficient fusion sites for initiation unit exchanges. Subsequently, they were validated by replacing the initiation region of endopyrrole pathway with a Cs-containing initiation unit for generation of a lipo-endopyrrole derivate. We promptly leveraged this strategy of changing initiation unit to target six previously silent NRPS BGCs, three BGCs were successfully activated and five novel lipopeptides were identified, demonstrating its application to recover silent BGCs. Furthermore, we extended this strategy in more BGCs from different bacteria. Utilizing a heterologous Cs-containing unit to replace the initiation region of chitinimide biosynthetic pathway led to successful incorporation of *N*-terminal fatty acid chains into chitinimide to create artificial lipo-chitinimides. This study provides a feasible strategy to rationally recover silent BGCs and add fatty acid chains to NRPs, enriching the genome mining and combinatorial biosynthesis approach for bacterial natural products.

**Significance:** Nonribosomal peptide synthetases (NRPSs) represent a valuable yet underexplored reservoir for bioactive natural products, but most of them are silent. This study introduces the concept that changing initiation unit to activate and optimize the functional expression of NRPSs, and successfully access three of six previously silent NRPS pathways, providing a feasible complement to the current genome mining approaches. Moreover, the *N*-terminal lipid chains are crucial for the activity of lipopeptides. We employed a heterologous Cs (starter condensation domain)-containing initiation unit to replace the original initiation region for creation of artificial nonribosomal lipopeptides, ultimately yielding three novel lipo-derivatives. This work presents a groundbreaking approach for activating and optimizing NRPSs and provides profound insight into the exploration of bacterial NRPS pathways.

## Introduction

Bacterial natural products (NPs) play a vital role in many fields such as medicine and agriculture due to their diverse chemical structure and biological activity (1–3). With the booming development of genome sequencing methods, a large number of microbial genome sequences have been published (4–6). Nonetheless, under standard laboratory conditions, most NP BGCs are silent or poorly expressed (7, 8). Currently, many NPs have been successfully identified using genome mining strategies (9–11), including OSMAC (12), promoter insertion (11, 13), regulatory gene overexpression (14), co-culture (15), transposon mutagenesis (16), ribosomal engineering (17, 18) and heterologous expression (19–21). However, it has been recently estimated that only 3% of bacterial genomes encoding potential NPs have been characterized (22). The vast realm of cryptic biosynthetic gene clusters (BGCs) represents a valuable source of novel bioactive compounds. However, the likelihood of activating gene clusters through genome mining remains low (10).

Failure in the field of genome mining to activate silent BGCs after a range of genome mining strategies may be due to insufficient availability of suitable substrates or diminished functionality of biosynthetic enzymes (16, 23–25). The initiation module serves as a critical control point that contributes to the overall specificity or functionality of the biosynthetic pathway (26, 27). If the starting module is inefficient, the entire synthesis process will be not work well. If the elongation modules or termination modules does not work properly, the product will be released from the assembly line in advance and the intermediate product may be detected. Consequently, streamlining the initiation module of silent biosynthetic pathways may provide a feasible alternative to current activation strategy to access their biosynthetic potential.

Nonribosomal peptide synthases (NRPSs) represents the largest class of secondary metabolic pathways found in bacteria and fungi, including initiation, elongation, and termination modules (5, 26–28). A typical NRPS generally contains three essential domains: adenylation (A), thiolation (T) and condensation (C) (28, 29). NRPs containing acyl chain at the N-terminus commonly initiate with starter condensation (Cs) domain, fatty acyl ligase (AL) / fatty acyl-AMP ligase (FAAL) (29, 30), or polyketide synthases (PKS) modules. The Cs domain introduces a fatty acid chain into the peptide by condensing substrates synthesized through the precursor biosynthesis pathway, which are typically activated by the acyl carrier protein (ACP) (31–34) or coenzyme A (CoA) (35–37). AL or FAAL selectively activate free fatty acids and loaded onto ACP or T domains transferred to NRPS assembly lines (38, 39). Based on different biosynthetic pathways, the initiating modules of NRPSs pathways are mainly composed of A-T, Cs-A-T and AL/FAAL-A-T. Here, we introduce the concept of the “initiation unit” to describe the initiating modules and the precursor biosynthetic pathway.

Engineering the “initiation unit” of NRPS offers the potential to improve yields, to change the relative proportion of products, and to generate new-to-nature peptides. Cheng’s team increased the proportion of polymyxin B1 by using the exchange of Cs domain or A domain to adjust the relative proportion of polymyxin analogues (40). Our previous study successfully obtained the co-crystallize structure of the RzmA-Cs (R148A) with octanoyl-CoA, and changed the fatty acid chains of the three lipopeptides by Cs domain swapping, establishing the novel method for acyl chain modification (41). The Kang group used a foreign FAAL homolog HmqF from *Burkholderia* sp. to replace DptE that loads medium-chain fatty acids to obtain purity daptomycin, and the yield was significantly increased by the gene fusion of FAAL and ACP (42).

The engineering methods for the “initiation unit” have been successfully developed and applied to a certain degree, yet genome mining still faces a high risk of failure. We speculate that the main reasons for its failure are likely due to: 1) inadequate precursor supply, and 2) inactivation of the initiation unit, including both the initial module and the precursor biosynthesis pathway. To address these challenges, we introduce the “initiation unit engineering” method to activate and optimize NRPSs gene clusters. This approach leverages initiation modules with primary metabolites as precursors to replace potentially deactivated initiation modules, overcoming the limitations of genome mining and enabling further product optimization to unlock the full biosynthetic potential of NRPSs.

In this study, we first compare the different fusion sites for changing the Cs domain-containing initiation regions *in vivo*. Then, based on the highly efficient fusion sites, we use the native Cs-containing unit to replace the initiation modules of endopyrrole biosynthetic pathway (*epy*) and other previously silent NRPS BGCs lead to generation of a new endopyrrole derivative containing an octenyl chain and the activation of three gene clusters, respectively. Furthermore, a heterologous Cs-containing unit was used to replace the initiation region of one NRPS pathways form phylogenetically distant species, yielding the new-to-nature products with an extra fatty acid chain. Through “initiation unit engineering”, we have activated and modified the underexplored NRPSs and unlocked their biosynthetic potential, providing a feasible strategy of genome mining and combinatorial biosynthesis for discovering and engineering untapped NRPs.

## Results

### Poor activity of initiation modules of silent NRPSs

Our previous studies identified two classes of lipopeptides, rhizomides and holrhizins, from *Mycetohabitans rhizoxinica* HKI 454 using *in situ* insertion of functional promoters in front of the core NRPS genes (43, 44), but this method is not workable on other NRPS BGCs in this strain. To investigate the reason why these NRPS BGCs are not activated, we selected six previously activate-failed NRPSs in this strain, named *2A, 2C, 3C, 5C, 7C* and *10C* based on their genomic locations, for functional validation of their initiation modules (Fig.4, *SI Appendix*, Table S4). These BGCs were directly cloned, inserted constitutive promoters and transferred to the chassis *Caldimonas brevitalea* DSM 7029 (45) (*Burkholderiales*, formerly *Schlegelella brevitalea* DSM 7029) for heterologous expression. The results indicated that there were no other special peaks compared to the control, further confirming that these BGCs are in a silent or low-expression state under laboratory conditions (*SI Appendix*, Fig. S2a and S2b).

If the initiation unit (here Cs domain) that responsible for loading lipid chains is inactive or lacks suitable precursors, the NRPSs may be silent or the product cannot be synthesized. To test this hypothesis, the Cs domain of RzmA was replaced with the Cs domains from several silent NRPSs *2C, 3C, 5C, epy*, and *10C* in *M. rhizoxinica* HKI 454, respectively. The Cs domain and the full-length linker between Cs and A domain from these BGCs were used for construction of recombinant plasmids. *C. brevitalea* DSM 7029 Δ*glb* was used as a heterologous chassis for product formation. The results showed that no derivatives were detected in all the fermentation crude extracts, providing evidences to support for our hypothesis that the poor activity of initiation units (*SI Appendix*, Fig. S3a and S3b). Conversely, using known functional Cs domains to replace silent ones could address the issue of inactive initiation modules while offer sufficient precursors, providing a new perspective for exploring biosynthetic potential of NRPSs.

### Screening of efficient fusion sites for initiation unit exchange

Although methods such as exchange units (XU, XUT) (46–48), module or domain substitution (C-A-T, Cs, A) (33, 40, 49) have been successfully applied to engineer some NRPs, the compatible fusion site (FS) is still the key to the success of NRPS engineering. We first aimed to identify an optimal fusion site to engineering the “initiation unit” for activation. The different bioinformatics analysis databases have different algorithms, the annotations of domain boundary will be different. We here used the “NRPS Motif Finder” platform which can resolve motifs and inter-motif structures of domains in NRPS (50), to create an initiation region bioinformatics analysis map using RzmA as a template. Ten conserved fusion sites (FS1∼FS10) to form ten initiation units (M1∼M10) containing Cs domains through the sequence alignment of multiple NRPS initiation regions were selected for comparison (Fig. 1, *SI Appendix*, Fig. S1). These sites include re-engineering points that have been successfully applied in NRPS so far (41, 47, 49), and each group avoids at least one co-evolving sector (50). The initiation module regions M1 to M10 correspond to CsN (N-lobe, 1–188 aa), Cs (with short linker, 1–350 aa), CsXU (1–442 aa), Cs (full-length linker, 1–479 aa), CsA (with short linker, 1–918 aa), CsA (full-length linker, 1–958 aa), CsAT (full-length linker, 1–1043 aa), CsATC_2_N (1–1226 aa), CsATC_2_XU (1–1492 aa), and CsATC_2_ (full-length linker, 1–1529 aa). By exchanging the initiation module regions M1 to M4 at the fusion sites FS1 to FS4, only the acyl chain of the product is altered. When exchanging modules M5 to M10 at the fusion sites FS5 to FS10, both the acyl chain and the first amino acid will be changed (Fig.1 and *SI Appendix*, Fig. S1).

**Figure 1.**
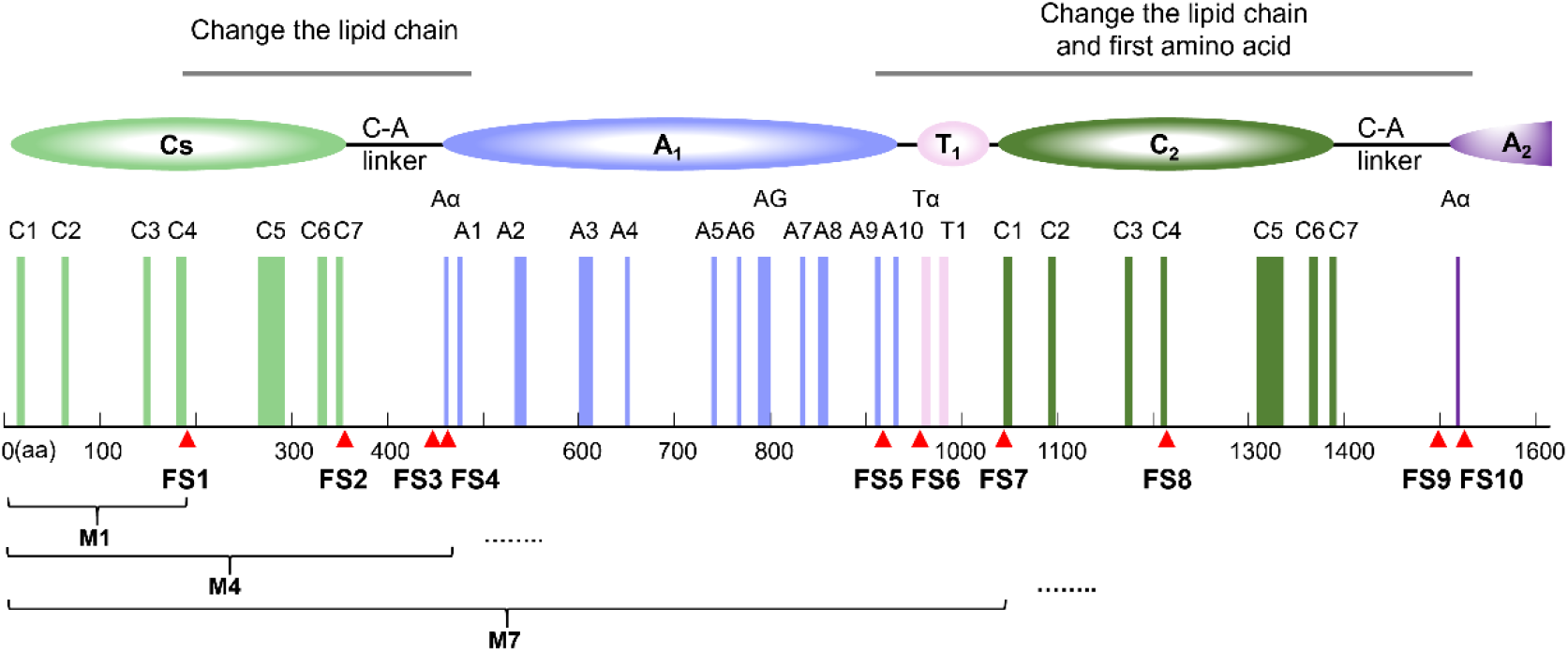
Selection of ten fusion sites of initiation unit containing the Cs domain. Information on the initiation regions of RzmA annotated by the “NRPS Motif Finder”. The light green and dark green represent the conserved motifs of the Cs domain and the second C domain, respectively. The light purple stripes indicate the conserved motifs of the first A domain, refers to the positions of the ten sites (FS1-FS10). M1, M4 and M7 represent three initiation modules composed of the Cs domains at sites FS1, FS4 and FS7, respectively.

We selected two previously activated NRPSs RzmA and HolA from *M. rhizoxinica* as the engineering materials. Both of them have initiation units that use precursors derived from primary metabolism and they are from the same strain as the above-mentioned silent BGCs, making them particularly suitable for applying the “initiation unit engineering” approach. The two nonribosomal lipopeptides synthesized, rhizomide A and holrhizin A, have fatty acid chains recognized by their Cs domains: acetyl-CoA (C2-CoA) and octanoyl-CoA (C8-CoA), respectively (Fig. 2A). The initial module regions M1 to M10 of HolA (HM1 to HM10) were used to replace the corresponding M1 to M10 modules in RzmA, creating recombinant plasmids with different exchanged modules (Fig.2 and *SI Appendix*, Fig. S5a). After that, the recombinant plasmid was transferred into *E. coli* GB05-MtaA (51) for heterologous expression, and the product was detected by high resolution electrospray ionization mass spectroscopy (HRESIMS). We found that the expected products could be detected in the crude extracts of all ten recombinant mutants, which were compound **1** at *m/z* 816.4834 [M+H] ^+^ (calc 816.4866) in the RzmA-HM1 to RzmA-HM4 and compound **2** at *m/z* 802.4689 [M+H] ^+^ (calc 802.4709) in the RzmA-HM5 to RzmA-HM10, with chemical formulas of C_41_H_65_N_7_O_10_ and C_40_H_63_N_7_O_10_, respectively (Fig. 2B and 2D, *SI Appendix*, Fig. S11).

**Figure 2.**
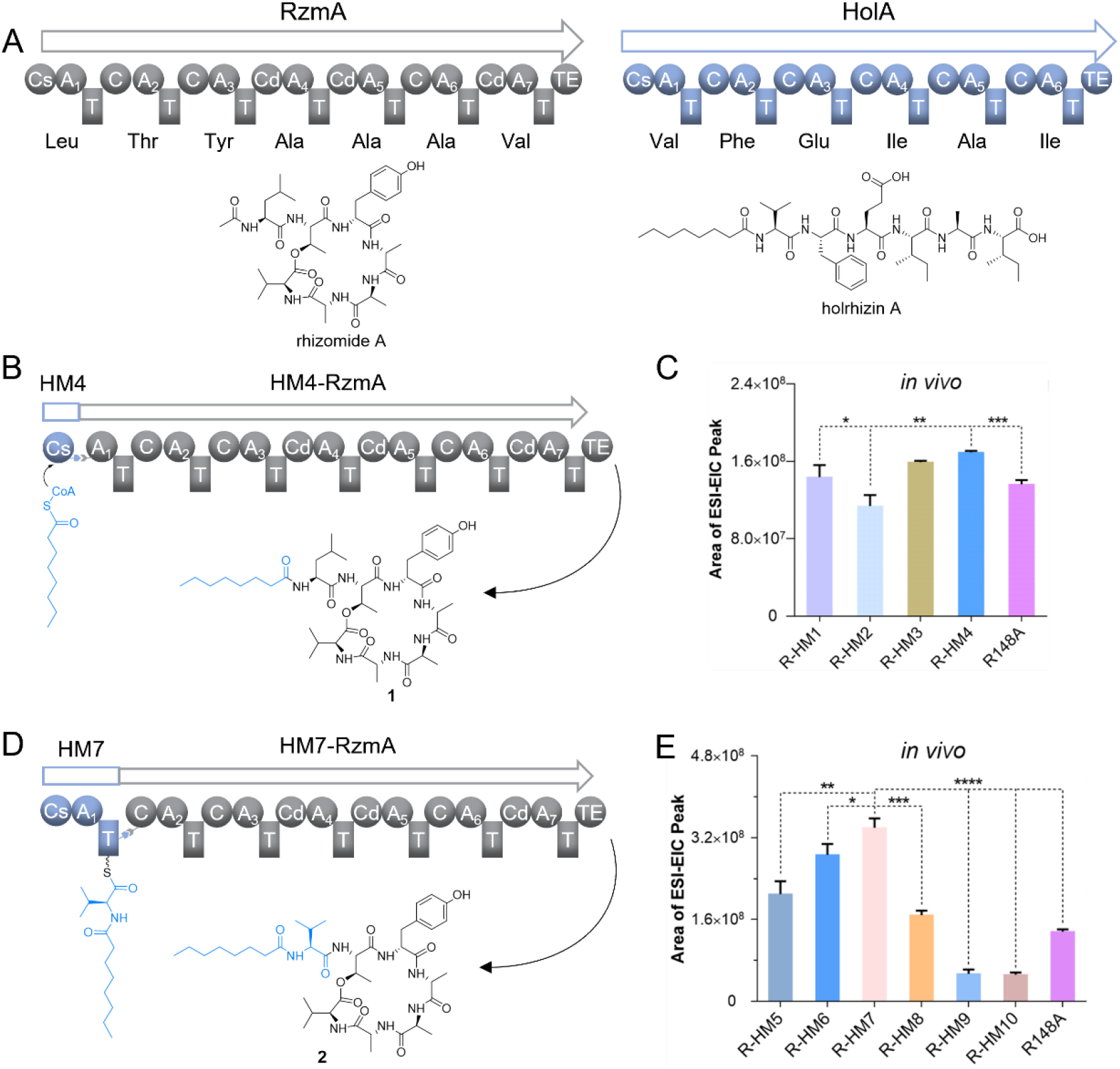
Screening of highly efficient fusion sites for initiation unit exchange. **A**. The *rzmA* and *holA* gene clusters of *M. rhizoxinica* HKI 454 and their main products rhizomide A and holrhizin A. Diagram of the biosynthesis of recombinant products after replacing the RzmA initiation region with different units from HolA, using HM4 (only the lipid chain is changed, **B**) and HM7 (the lipid chain and the first amino acid are changed, **D**) as examples. Comparison of the yields of rhizomide A derivatives **1** (**C**) and **2** (**E**) after swapping with different initiation units from HolA. R148A represents RzmA-Cs*, a mutant of RzmA with a point mutation R148A in the Cs domain, which can recognize the octanoyl-CoA. *P* values were calculated by two-tailed unpaired *t* test. \**p* < 0.05, \*\**p* < 0.01, \*\*\**p* < 0.001 and \*\*\*\**p* < 0.0001.

The relative quantification results show that the product yield was higher when substituting with the initiation regions M4, M5, M6, M7 and M8 of HolA. Specifically, if only the Cs domain is replaced to change the acyl chain, the replacement of the full-length Cs domain (HM4) is more effective. HM4—where the Cs domain containing the full-length linker from HolA was substituted—yielded slightly better results than the previously reported site HM3. When both the acyl chain and the first amino acid were altered, HM7, the CsAT initiation module exchange with full-length linker had a higher yield, and it was also the highest yield among all groups. Moreover, as the initiation module from HM2 to HM7, the replacement region extends from the Cs domain to the full-length CsAT module, resulting in an increasing yield. When the length of exchange region is further extended to the C domain of the second C-A-T module, the yield begins to decline. Compared to HM8, the yields for HM9 and HM10 as substitutes were approximately three times lower, while the effect of HM8 was similar to HM4 (Fig. 2C and 2E). Therefore, when engineering the starting module, the preferred fusion sites should be FS4 and FS7, depending on the specific requirements.

### Verification of initiation unit exchange in the native strain

To evaluate the availability of the screened sites, we initially applied it to alter the starting region of the endopyrrole A biosynthetic gene cluster (*epy*) in same strain *M. rhizoxinica* HKI 454. The product of *epy* was only activated by co-culture of the HKI 454 with its fungal host, but it is silent in varied culture conditions of itself (15), suggesting that insufficient supply of specific pyrrole substrates may be responsible for its silent (Fig. 3A).

**Figure 3.**
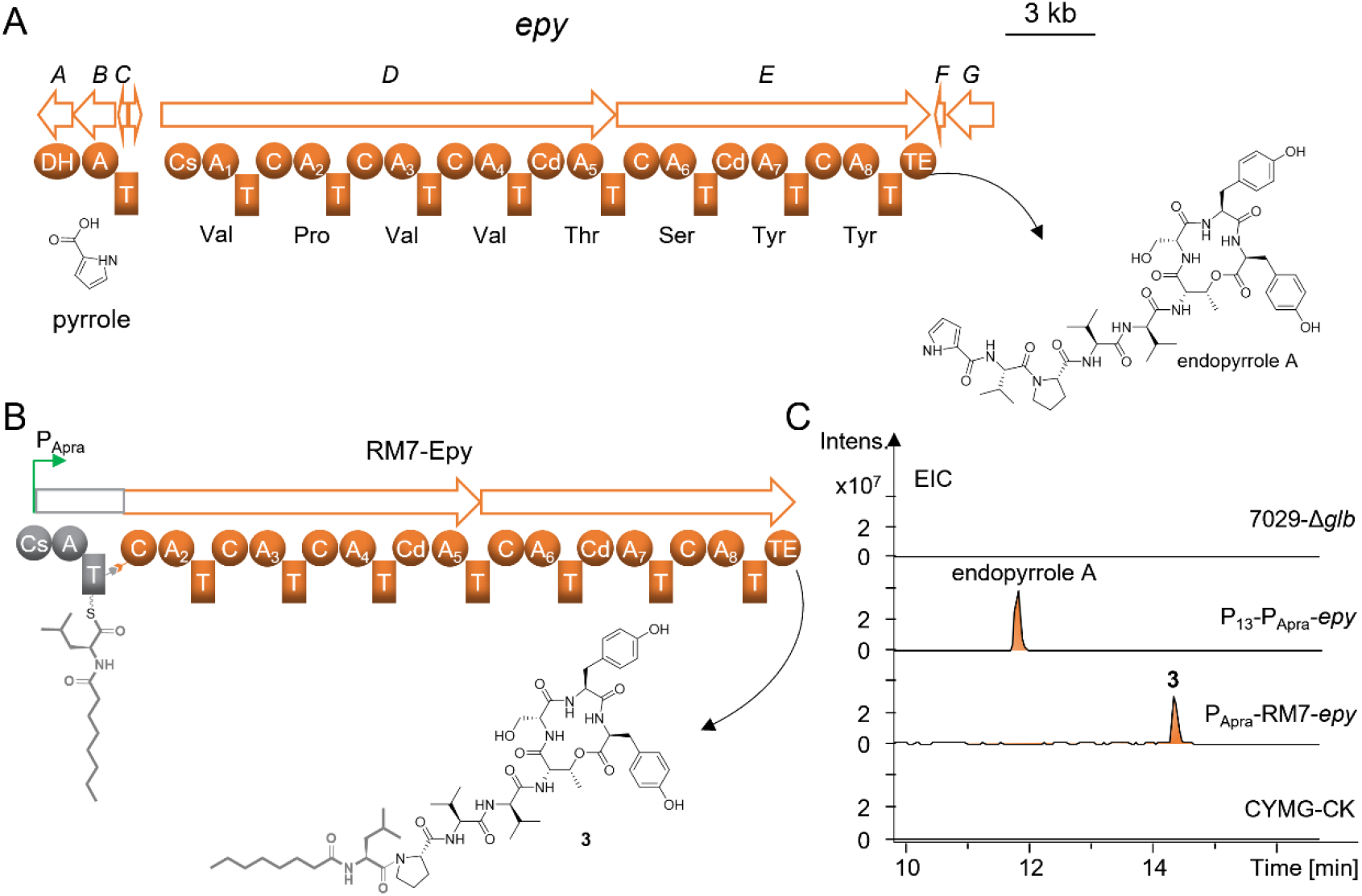
Changing the initiation module of endopyrrole pathway in *M. rhizoxinica* HKI 454. **A**. The *epy* BGCs from *M. rhizoxinica* HKI 454 and its main product endopyrrole A. **B**. Using the fusion site (FS7) of RzmA -Cs*AT to replace the initiation module, and produce compound **3. C**. comparative HPLC-MS analysis (Extracted ion chromatograms at *m/z* 1002.4931 [M+H] ^+^ for endopyrrole A, *m/z* 1049.5918 [M+H] for **3**) of crude extracts from the heterologous host 7029-Δ*glb* harboring the *epy* recombinants obtained by promoter insertion (P_13_-P_Apra_-*epy*) and RM7 replacement (P_Apra_-RM7-*epy*) using the 7029-Δ*glb* and medium (CYMG) as controls.

We directly cloned *epy* BGC and inserted the constitutive promoters P_Apra_ and P_13_ (52) to promote expression of the core gene and the gene responsible for the specific precursor synthesis, respectively (*SI Appendix*, Fig. S4). The *epy* BGC was functional expressed in heterologous host *C. brevitalea* DSM 7029 and produce the endopyrrole A with *m/z* 1001.4858 [M+H] ^+^(Fig. 3C and *SI Appendix*, Fig. S4). Next, we used RzmA-Cs*A_Leu_T (RM7) to replace the EpyD-CsA_Val_T as a new initiation module, the pyrrole precursor section and the first Val will be replaced with an octanoyl chain and Leu (Fig. 3B). Cs* is a Cs R148A mutant of the Arg_148_ point to Ala in RzmA-Cs domain, which can enhance the binding affinity for the octanoyl chain without affecting its ability to recognize the acetyl chain (41). By HPLC-HRESIMS analysis, we obtained a compound **3** with *m/z* 1049.5896 [M+H] ^+^ (C_54_H_80_N_8_O_13_) containing a signature fragment ion peak at *m/z* 240.1957 [M+H] ^+^ (calc 240.1958, octanoyl-Leu). By comparative analysis of the MS/MS fragmentations with endopyrrole A suggested compound **3** is the derivative of endopyrrole A with an *N*-terminal octanoyl chain and the Val was substituted as Leu, which is consistent with the design of initiation unit exchange (Fig. 3B and C, *SI Appendix*, Fig. S12). The findings indicate that changing the initiation module with the Cs-containing unit to activate silent NRPS pathways is promising.

### Changing initiation unit to access silent NRPSs

We next aimed to access the six silent NRPS in *M. rhizoxinica* HKI 454 by applying the selected sites with initiation units (Fig. 4A and *SI Appendix*, Table S4). Using recombineering (53), we first performed the initiation modules from RzmA to exchange the starter regions of NRPSs in *M. rhizoxinica* HKI 454 Δ*rhi* with the rhizoxin BGC was inactivated. Since the amino acids recognized by the first A domain of these NRPSs are unknown, we mainly chosen the initiation units from RM5 to RM10 containing Cs and A domains, with a specific emphasis on RM7 (Fig. 2D, 2E and 4A). (*SI Appendix*, Fig. S5c).

**Figure 4.**
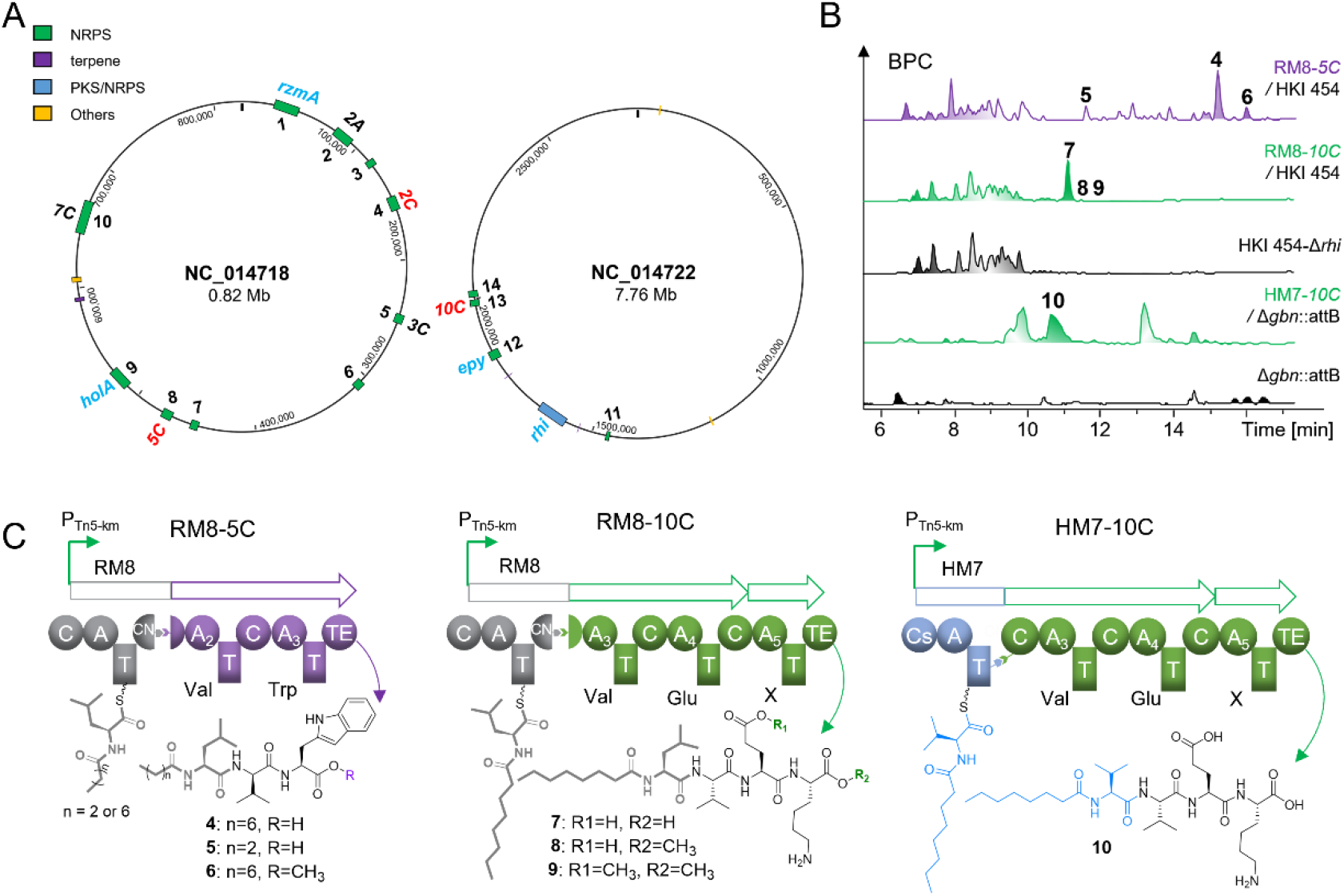
Changing the initiation unit to access previously silent NRPS BGCs in *M. rhizoxinica* HKI 454. **A**. Prediction of NRPS BGCs in HKI 454 by anti-SMASH. The blue font represent the previously reported BGCs, and the red font represents the BGCs that products were successfully obtained in this study. **B**. HPLC-MS analysis of the crude extracts of wild-type strains and the recombinant strains obtained by initiation unit exchanges. HKI 454-Δ*rhi* represents the HKI 454 with the inactivation of rhizoxin BGCs, Δ*gbn*::*attB*: represents the chassis strain *B. gladioli* Δ*gbn::attB* for engineered BGCs heterologous expression. Compounds **4, 5** and **6** are products derived from RM8-5C, **7** and **10** are products activated from RM8-10C and HM7-10C, respectively. **C**. Schematic representation of product synthesis after modification of *5C* gene cluster by RzmA initiation module RM8. **D** and **E**. Schematic representation of product synthesis after modification of *10C* gene cluster by the initiation units, RM8 of RzmA and HM7 from HolA, respectively.

We constructed HKI 454 recombinant strains of RM7-2A, RM7-2C, RM7-3C, RM8-5C, RM7-7C, and RM8-10C. The C domain in the second module of *10C* is a C_dual_ domain that can alter the configuration of upstream amino acids. To prevent the effect of this donor specificity, we replaced the CsAT-C_2_A_2_T_2_-C_3_N of *10C* with RM8 (CsATC_2_N). The new peaks were detected in three recombinant strains (RM8-2C, RM8-5C and RM8-10C) of them compared to the wild type by LC-MS analysis (Fig. 4B). We also performed “initiation unit engineering” on the directly cloned BGCs *5C* and *10C*, and conducted heterologous expression in strain DSM 7029 Δ*glb* (Fig. 4C-4F and SI Appendix, Fig. S5b). The results were consistent with those observed in HKI 454, the same product was detected, indicating that these three silent NRPSs were successfully activated (Fig. 4A and SI Appendix, Fig. S7a).

The anti-SMASH analysis of gene cluster *5C* predicted that it contains three C-A-T modules, starting with a Cs domain. However, the amino acid recognized by the first A domain is unknown, while the second and third A domains are predicted to recognize Val and Trp, respectively. We selected and compared four fusion sites—FS5, FS7, FS8 and FS9—using the starting modules RM5, RM7, RM8, and RM9 from RzmA for substitution, and transferred into different heterologous hosts for product detection (Fig. 4C and SI Appendix, Fig. S6a). All the mutants can produce two compounds **4, 5** except the RM5-5C mutant strain (Fig. 4C). Additionally, our experiments revealed that the *Burkholderia gladioli* Δ*gbn::attB* with cleaner metabolic background and the improved yields compared to the DSM 7029 Δ*glb* chassis (SI Appendix, Fig. S6b). The products were purified, and elucidated through HRESIMS and NMR spectroscopy. Compound **4** was isolated as colorless oil with the chemical formula established as C_30_H_46_N_4_O_5_ with *m/z* 543.3519 [M+H] ^+^(calc 543.3541) though MS (*SI Appendix*, Fig. S13 and Tables S5). The structure was determined on the data of DEPT (90 and 135) spectrum, the amino acid predicted by A domain, as well as 2D correlation NMR spectroscopies including ^1^H-^1^H correlation spectroscopy (^1^H-^1^H COSY), heteronuclear multiple bond correlation (HMBC) and heteronuclear single quantum coherence spectroscopy (HSQC). Interpretation of the NMR data established three proteinogenic amino acid residues including Leu, Val and Trp and an octanoic acid residue (*SI Appendix*, Fig. S19-S24 and Tables S5). These residues were linked together to form a linear lipopeptide based on HMBC correlations (*SI Appendix*, Fig. S18). The configurations of the amino acid residues were determined to be L-Leu, D-Val and L-Trp by the Marfey’s method in combination with the presence of a dual condensation domain (*SI Appendix*, Fig. S10). In addition to **4** with an octenyl chain, we also identified derivatives **5** and **6**, molecular formula of C_24_H_34_N_4_O_5_ and C_31_H_48_N_4_O_5_, with *m/z* 459.2602 [M+H] ^+^ (calc 459.2594) and *m/z* 557.3674 [M+H] ^+^ (calc 557.3697) (*SI Appendix*, Fig. S9 and Tables S5). Compared to the NMR data of **4, 5** possesses an acetyl chain instead of the octanoyl chain in **4**, and **6** has a *O*-methylation modification of Trp at the C-terminus, respectively (*SI Appendix*, Fig. S18 and S25-S36).

HRESIMS analysis of the crude extract from RM8-*10C*/ HKI 454 Δ*rhi* revealed that three more peaks **7,8,9** compared to the control (Fig. 4D and 4E) (SI Appendix, Fig. S15). The chemical formula of **7** was identified to C_30_H_55_N_5_O_8_ at *m/z* 614.4149 [M+H] ^+^ (calc 614.4132), accompanied by a signature fragment ion peak at *m/z* 240.1959 [M+H] ^+^ (calc 240.1958, octanoyl-Leu) (*SI Appendix*, Fig. S14 and Tables S6). The NMR data revealed the presence of four proteinogenic amino acids, and the HMBC correlations confirmed their order in the formation of the linear lipopeptide (*N*-octanoyl-Leu-Val-Glu-Lys) (*SI Appendix*, Fig. S37-S42 and Tables S6). The absolute configuration of the amino acids, L-Leu, L-Val, L-Glu and L-Lys, was determined using the Marfey’s method along with bioinformatics analysis of the C domains (SI Appendix, Fig. S10). (Fig. 4D and 4E, SI Appendix, Fig. S18). The HR-ESI-MS analysis of **8** and **9** suggested their molecular formulas are C_31_H_57_N_5_O_8_ and C_32_H_59_N_5_O_8_, with *m/z* values of 628.4288 [M+H] ^+^ (calc 628.4280) and 642.4439 [M+H] ^+^ (calc 614.4436), respectively (SI Appendix, Fig. S15), calculating one (**8**) or two (**9**) more -CH_2_ groups than **7**. According to the MS/MS fragmentation analysis, **8** contains a *O*-methyl group on the carbonyl of Lys, and **9** harbors two *O*-methyl groups on the carbonyl moieties of Lys and Glu, respectively. (SI Appendix, Fig. S14 and S15). *O*-methylation of C-terminal amino acids in non-ribosomal peptides are not rare in *Burkholderiales* (45). This may occur either due to spontaneously during the preparation of peptide compounds in acidified methanol/aqueous solutions, special type of TE domain in the NPRS(54), or the *O*-methyltransferase present in the genome (55). The second and third A domains are predicted to recognize Val, which is same as the first activated amino acid of the HolA (Fig. 2A and 4A). Therefore, we tried to change the initiation CsAT-C_2_AT region of 10C using the HM7 module from HolA for heterologous expression in the *B. gladioli* Δ*gbn::attB*. An expected product **10** was detected in the crude extract with *m/z* 600.3978 [M+H] ^+^ (calc 600.3967, cacl C_29_H_53_N_5_O_8_) (Fig. 4F and SI Appendix, Fig. S16). MS/MS analysis showed the signature fragment ion peak was changed from *m/z* 240.1959 [M+H] ^+^ (octanoyl-Leu) in **7** to *m/z* 226.1805 [M+H] ^+^ (octanoyl-Val) in **10** (*SI Appendix*, Fig. S11, S16 and S18). The NMR spectra suggested **10** is a linear lipopeptide *N*-octanoyl-Val-Val-Glu-Lys, the first amino acid Val in **10** is different to the first amino acid Leu in **7** (SI Appendix, Fig. S43-S48 and Tables S6). Compounds **7** and **10** are both activated derivatives by altering the initiation region of *10C* using functional initiation modules from two NRPSs, suggesting that the strategy of changing the initiation modules can explore the potential biosynthetic capacity and enrich the structural diversity of non-ribosomal peptides.

Unfortunately, the structure of the product activated from BGC *2C* failed to be identified due to poor solubility for NMR recording (*SI Appendix*, Fig. S7). We assessed the anti-inflammatory activity of four compounds (**4, 5, 7** and **10**), the results determined that compound **7** exhibited potent effect, reducing NO levels by approximately 30% at 40 μM (*SI Appendix*, Fig. S9). All of them didn’t show any antibacterial activity or cytotoxicity.

For other three BGCs, we attempted various FSs for swapping the initiation regions, but all failed to produce any desired compounds. We speculate that the recombination of genes may have disrupted the normal interactions between domains or other factors are needed for its biosynthesis. Moreover, there are many reasons about silent BGCs and many obstacles to the proper functioning of the initiation module. And the possibilities that elongation modules inactivation cannot be ignored. The strategy proposed in this study aims to address two main issues: the loss of activity of the initiation module and the insufficient supply of substrates required for initiation. Successfully activating three of six silent gene clusters in this strain that had previously failed in genome mining demonstrates the potential applicability of this novel strategy to recover silent NRPS pathways.

### Adding a lipid chain into NRPs by heterologous initiation unit exchange

Lipid chains play a crucial role in the biological activity of NRPs (29). The lipophilic fatty acid chain facilitates the anchoring of lipopeptides to the phospholipid bilayer of cell membranes, increasing interaction time with targets (56, 57). Additionally, the length of the lipid chain in non-ribosomal lipopeptides can balance their biological activity and cytotoxicity (58). Engineering the “initiation unit” can also be applied to optimize yields of targeted or even converting the non-ribosomal peptides without fatty chains into lipopeptides. Adding a fatty acid chain at the *N*-terminus of NRPs by replacing their initiation module that containing the Cs domain can alter their chemical properties, enhancing activity and drug potential.

We extend this strategy to more BGCs in other species. Our previous study showed heterologous expression of a cryptic *chm* BGC from *Chitinimonas koreensis* DSM 17726 (family *Burkholderiaceae*) in *C. brevitalea* DSM 7029 leading to the discovery of new NRPs, chitinimides (45). The initiation ChmA is in charge of the fragment containing the ureido-bond between the first Phe and the second dehydrobutyrine (Dhb) by a head-to-head manner which performed via the first ureido-bond formation C_u_ domain (Fig. 5A). Dhb residue is formed from the dehydration of Thr catalyzed by the downstream C domain (27). To add a *N*-terminal lipid chain into the chitinimides, the initiation A_Phe_-T-C_u_ region should be removed meanwhile a Cs-containing initiation unit with preferentially load downstream Dhb residue should be introduced, aiming to avoid the formation of urea bond and improve the adaptability between the exogenous Cs and the targeted NRPS. We found that the first amino acid residue in the lipopeptide burriogladiodin (bgdd) from *Burkholderia gladioli* ATCC 10248 is Dhb (59), so the Cs domain in this pathway was selected to replace the initiation A_Phe_-T-C_u_ region in chitinimide biosynthetic pathway using the efficient fusion site FS4 screened by the above experiment (Fig. 2B and 5B) (Fig. 5A). The *chm* BGC plasmid was engineered and the constructed plasmid BM4-*chm* was integrated into the genome of *C. brevitalea* DSM 7029 Δ*glb*, and the correct mutant strains were screened for fermentation detection.

**Figure 5.**
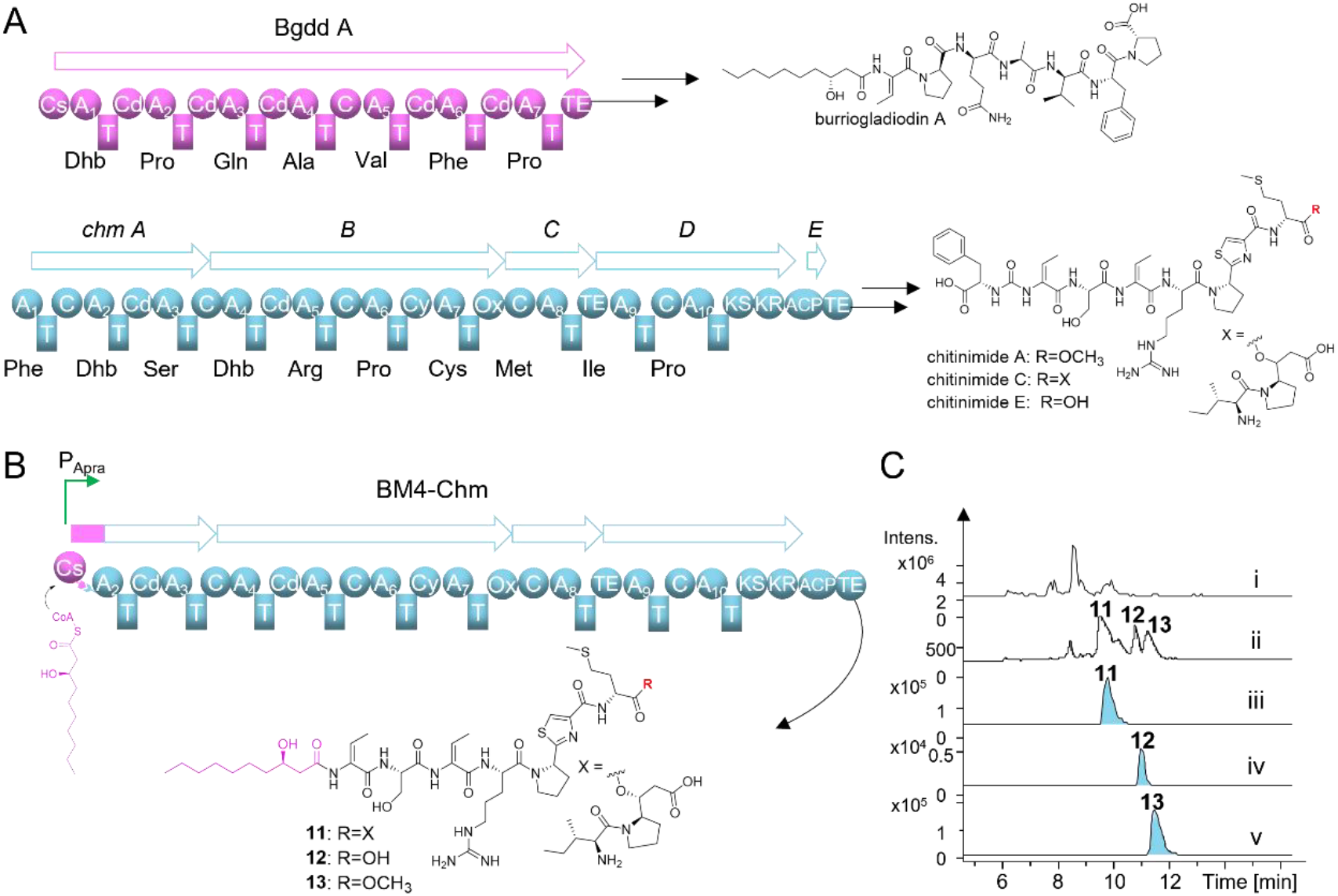
Introducing *N*-terminal fatty acid chains by engineering the initiation region of ChmA. **A**. The *bgdd* and *chm* BGCs and their main products burriogladiodin A and chitinimides A, C and E from *B. gladioli* ATCC 10248 and *C. koreensis* DSM 17726, respectively. **B**. Schematic representation of the products after engineering of *chm* BGC by the initiation unit BM4 of Bgdd. **C**. The UV spectrum (254nm, ii) and HPLC-MS analysis (base peak chromatograms at *m/z* 1163.5951 [M+H] ^+^for **11**, *m/z* 909.4321 [M+H] ^+^ for **12** and *m/z* 923.4478 [M+H] ^+^ for **13**, i, iii, iv, v) of crude extracts from strain DSM 7029Δ*glb*/BM4-*chm*.

Metabolic profiling of the fermentation crude extract revealed three distinct absorption peaks at 254 nm, corresponding to products **11, 12**, and **13**, with compound **13** having the highest yield (Fig. 5C). HRESIMS determined that the molecular formula of **13** was C_41_H_66_N_10_O_10_S_2_ at *m/z* 923.4473 [M+H] ^+^ (calc 923.4478) (*SI Appendix*, Fig. S13, S43-S51 and Tables S7). Further detailed analysis NMR data combined with MS/MS spectrum showed that **13** is composed of a *β*-OH decanoate chain and seven amino acid residues (Dhb, L-Ser, Dhb, L-Arg, L-Pro, L-Cys and L-Met), which is a derivative of chitinimide A with a lipid chain (*SI Appendix*, Fig. S52-S54). HRESIMS and MS/MS analysis suggested that the chemical formulas of **11** (C_53_H_86_N_12_O_13_S_2_ with *m/z* 582.3033 [M+2H] ^2+^ (calc 582.3012)) and **12** (C_40_H_64_N_10_O_10_S_2_ at *m/z* 909.4319 [M+H] ^+^ (calc 909.4321)) are derivatives of chitinimides C and E with a lipid chain, respectively (*SI Appendix*, Fig. S17). (Fig. 5 and *SI Appendix*, Fig. S17).

Cytotoxicity assays demonstrated that **12** exhibited moderate cytotoxicity against HepG2 and A549 tumor cell lines with IC_50_ values of 8.36 μM and 12.02 μM, respectively, which is higher than that of **11, 13**, and chitinimide A. Due to the extremely low yield of chitinimide E, it failed to purify for comparative test (*SI Appendix*, Table S8). The results indicate that the initiation unit exchange strategy can be applied not only to activate silent BGCs but also to structurally optimize the NRPs, facilitating the access underexplored biosynthetic potential.

## Discussion

Bacterial genomes contain numerous cryptic or silent NRPS gene clusters that potentially encode for the synthesis of non-ribosomal peptides with biologically active (7, 11, 22). Genome mining methods have released the secondary metabolic potential of part cryptic BGCs, but these only are a very small proportion of the large-scale unexplored genomic data. In this study, we introduce an alternative to the current genome mining strategy by the exchange of the initiation units of the NRPSs, which is a feasible way to access silent biosynthetic pathways and to generate new-to-nature nonribosomal peptide derivatives.

Current genome mining methods like OSMAC, promoter replacement, and regulatory gene modification can be attempted to cope with some silent or low-expressed BGCs for the reason of substrate deficiency, weak of original promoter, or strict regulatory network. (11, 34, 60). However, if the core biosynthetic enzyme responsible for biosynthesis is low in activity or loses function during evolution, the biosynthetic pathway will remain silent even with a favorable external environment or high expression level. In most cases, after replacing the promoter, the products cannot be gained in the crude extracts for the silent NRPS, including intermediates. Therefore, the initiation module may be the key to activating NRPS expression, serving as a potential “one-size-fits-all solution” to unlock the biosynthetic potential of NRPSs.

When swapping the initial Cs domains of two known gene clusters, the corresponding derivatives were detected (41). However, replacing RzmA-Cs with Cs domain of silent NRPS in this work resulted in no product formation, further supporting the possibility that the initiation module problem caused the silent gene cluster. Here, we activate silent NRPS BGCs by designing initiation modules to recover the function of the start regions, thereby enabling to directly utilize substrates from primary metabolism or achieving detectable levels of product yield. The fusion sites for swapping of initiation module in this study can serve as a reference for various types of NRPSs. With further analysis and optimization, these sites could be universally applicable in different contexts.

The Cs domains are responsible for lipid chain loading in non-ribosomal lipopeptides synthesis (SI Appendix, Fig. S8a), and exhibits variability in substrate selection and specificity across various NRPS gene clusters. Some are able to exclusively recognize and condense a single fatty acyl substrate from primary metabolism or with different modifications, whereas others are substrate tolerance and have the ability to synthesize diverse lipid chain (34, 61, 62). Recent studies have also focus on engineering Cs domain to affect selectivity for lipid chain substrates (30, 40, 41). In this study, we incorporated lipid chains into the *chm* gene cluster, which originally lacked them, offering a new perspective for optimizing NRPSs.

The initial synthesis process of Cs-A-T module in NRPS is dynamic, with the A_core_ subunit of A domain binding to the amino acid substrate to form aminoacyl-AMP, and interacts with the thiol of 4’-phosphopantetheine (Ppant) arm of T domain to create aminoacyl-T (SI Appendix, Fig. S8). Then, the flexible A_sub_ subunit swings to the Cs domain with aminoacyl-T, where the Cs domain condenses the fatty acyl chain with aminoacyl-T to form a peptide bond (28, 63, 64). For the failed experiments or the different yields of recombinant products in this study may be due to the replaced modules forming fusion proteins with the original NRPS, disrupting the interaction interface of protein. The highest yield of HM7 (CsAT with full-length linker) suggests that this replacement preserves the effective interactions among the original C, A and T domains within the initiation module. An in-depth study of the dynamic interactions between domains can be crucial for designing NRPS to produce lipopeptides with novel structures and functions. Activating and optimizing NRPSs using functional initiation unit exchange from native or phylogenetically-closed strain can serve as a more preferentially efficient strategy.

## Materials and Methods

Detailed description of methods covered in the text including bacterial strains, reagents and growth conditions, plasmid construction, bioinformatics analysis, detailed design and replacement of the initiation unit, *in situ* modification of the NRPSs initiation units in *M. rhizoxinica* HKI 454, fermentation and HPLC-MS analysis of metabolic crude extracts, purification and characterization of compounds, Marfey’s analysis and biological activity screening, are given in *Supporting Information, Supplementary Materials and Methods*.

## Acknowledgments

We thank Guannan Lin, Zhifeng Li, Jingyao Qu, Jing Zhu, Haiyan Sui and Xiangmei Ren, from State Key Laboratory of Microbial Technology of Shandong University for help and guidance in HRMS, NMR and LC. This work was supported by National Key R&D Program of China (2019YFA0905700), National Natural Science Foundation of China (32070060), Fundamental Research Funds of Shandong University (2023QNTD001), SKLMT Frontiers and Challenges Project (SKLMTFCP-2023-05), and Shandong Provincial Natural Science Foundation (ZR2023ZD29).

## References

1. J. D. Hegemann, J. Birkelbach, S. Walesch, R. Müller, Current developments in antibiotic discovery: Global microbial diversity as a source for evolutionary optimized anti -bacterials. EMBO Rep 24, e56184 (2023).

2. A. K. Shakya, R. R. Naik, The Chemotherapeutic Potentials of Compounds Isolated from the Plant, Marine, Fungus, and Microorganism: Their Mechanism of Action and Prospects. J Trop Med 2022, 1–17 (2022).

3. A. G. Atanasov, S. B. Zotchev, V. M. Dirsch, C. T. Supuran, Natural products in drug discovery: advances and opportunities. Nat Rev Drug Discov 20, 200–216 (2021).

4. K. Blin, et al., antiSMASH 7.0: new and improved predictions for detection, regulation, chemical structures and visualisation. Nucleic Acids Res 51, W46–W50 (2023).

5. S. A. Kautsar, K. Blin, S. Shaw, T. Weber, M. H. Medema, BiG-FAM: the biosynthetic gene cluster families database. Nucleic Acids Res 49, D490–D497 (2021).

6. P. Cimermancic, et al., Insights into Secondary Metabolism from a Global Analysis of Prokaryotic Biosynthetic Gene Clusters. Cell 158, 412–421 (2014).

7. X. Zhang, Hindra, M. A. Elliot, Unlocking the trove of metabolic treasures: activating silent biosynthetic gene clusters in bacteria and fungi. Curr Opin Microbiol 51, 9–15 (2019).

8. K. Scherlach, C. Hertweck, Mining and unearthing hidden biosynthetic potential. Nat Commun 12, 3864 (2021).

9. M. G. Chevrette, et al., The confluence of big data and evolutionary genome mining for the discovery of natural products. Nat. Prod. Rep. 38, 2024–2040 (2021).

10. K. D. Bauman, K. S. Butler, B. S. Moore, J. R. Chekan, Genome mining methods to discover bioactive natural products. Nat. Prod. Rep. 38, 2100–2129 (2021).

11. B. Baral, A. Akhgari, M. Metsä-Ketelä, Activation of microbial secondary metabolic pathways: Avenues and challenges. Synth Syst Biotechnol 3, 163–178 (2018).

12. R. Pan, X. Bai, J. Chen, H. Zhang, H. Wang, Exploring Structural Diversity of Microbe Secondary Metabolites Using OSMAC Strategy: A Literature Review. Front. Microbiol. 10, 294 (2019).

13. D. Montiel, H.-S. Kang, F.-Y. Chang, Z. Charlop-Powers, S. F. Brady, Yeast homologous recombination-based promoter engineering for the activation of silent natural product biosynthetic gene clusters. Proc. Natl. Acad. Sci. U.S.A. 112, 8953–8958 (2015).

14. B. Wang, F. Guo, S.-H. Dong, H. Zhao, Activation of silent biosynthetic gene clusters using transcription factor decoys. Nat Chem Biol 15, 111–114 (2019).

15. S. P. Niehs, et al., Genome Mining Reveals Endopyrroles from a Nonribosomal Peptide Assembly Line Triggered in Fungal–Bacterial Symbiosis. ACS Chem. Biol. 14, 1811–1818 (2019).

16. D. Mao, A. Yoshimura, R. Wang, M. R. Seyedsayamdost, Reporter-Guided Transposon Mutant Selection for Activation of Silent Gene Clusters in Burkholderia thailandensis. ChemBioChem 21, 1826–1831 (2020).

17. F. Stevenson-Jones, J. Woodgate, D. Castro-Roa, N. Zenkin, Ribosome reactivates transcription by physically pushing RNA polymerase out of transcription arrest. Proc. Natl. Acad. Sci. U.S.A. 117, 8462–8467 (2020).

18. T. Hosaka, et al., Antibacterial discovery in actinomycetes strains with mutations in RNA polymerase or ribosomal protein S12. Nat Biotechnol 27, 462–464 (2009). 1.

19. M. Komatsu, T. Uchiyama, S. Ōmura, D. E. Cane, H. Ikeda, Genome-minimized Streptomyces host for the heterologous expression of secondary metabolism. Proc. Natl. Acad. Sci. U.S.A. 107, 2646–2651 (2010).

20. J. Ke, Y. Yoshikuni, Multi-chassis engineering for heterologous production of microbial natural products. Curr Opin Biotechnol 62, 88–97 (2020).

21. L. Huo, et al., Heterologous expression of bacterial natural product biosynthetic pathways. Nat. Prod. Rep. 36, 1412–1436 (2019).

22. A. Gavriilidou, et al., Compendium of specialized metabolite biosynthetic diversity encoded in bacterial genomes. Nat Microbiol 7, 726–735 (2022).

23. B. C. Covington, F. Xu, M. R. Seyedsayamdost, A Natural Product Chemist’s Guide to Unlocking Silent Biosynthetic Gene Clusters. Annu. Rev. Biochem. 90, 763–788 (2021).

24. Z. Liu, Y. Zhao, C. Huang, Y. Luo, Recent Advances in Silent Gene Cluster Activation in Streptomyces. Front Bioeng Biotechnol 9, 632230 (2021).

25. M. Stasiak, E. Maćkiw, J. Kowalska, K. Kucharek, J. Postupolski, Silent Genes: Antimicrobial Resistance and Antibiotic Production. Pol J Microbiol 70, 421–429 (2021).

26. M. A. Fischbach, C. T. Walsh, Assembly-Line Enzymology for Polyketide and Nonribosomal Peptide Antibiotics: Logic, Machinery, and Mechanisms. Chem. Rev. 106, 3468–3496 (2006).

27. S. Dekimpe, J. Masschelein, Beyond peptide bond formation: the versatile role of condensation domains in natural product biosynthesis. Nat. Prod. Rep. 38, 1910–1937 (2021).

28. R. D. Süssmuth, A. Mainz, Nonribosomal Peptide Synthesis—Principles and Prospects. Angew Chem Int Ed 56, 3770–3821 (2017).

29. Y.-H. Chooi, Y. Tang, Adding the Lipo to Lipopeptides: Do More with Less. Chem Biol 17, 791–793 (2010).

30. Q. Liu, W. Fan, Y. Zhao, Z. Deng, Y. Feng, Probing and Engineering the Fatty Acyl Substrate Selectivity of Starter Condensation Domains of Nonribosomal Peptide Synthetases in Lipopeptide Biosynthesis. Biotechnol J 15, 1900175 (2020).

31. F. I. Kraas, T. W. Giessen, M. A. Marahiel, Exploring the mechanism of lipid transfer during biosynthesis of the acidic lipopeptide antibiotic CDA. FEBS Letters 586, 283–288 (2012).

32. D. C. Alexander, et al., Production of novel lipopeptide antibiotics related to A54145 by Streptomyces fradiae mutants blocked in biosynthesis of modified amino acids and assignment of lptJ, lptK and lptL gene functions. J Antibiot 64, 79–87 (2011).

33. K. T. Nguyen, et al., Combinatorial biosynthesis of novel antibiotics related to daptomycin. Proc. Natl. Acad. Sci. U.S.A. 103, 17462–17467 (2006).

34. R. H. Baltz, Genome mining for drug discovery: cyclic lipopeptides related to daptomycin. J Ind Microbiol Biotechnol 48, kuab020 (2021).

35. F. I. Kraas, V. Helmetag, M. Wittmann, M. Strieker, M. A. Marahiel, Functional Dissection of Surfactin Synthetase Initiation Module Reveals Insights into the Mechanism of Lipoinitiation. Chem Biol 17, 872–880 (2010).

36. C. Wang, C. J. Flemming, Y.-Q. Cheng, Discovery and activity profiling of thailandepsins A through F, potent histone deacetylase inhibitors, from Burkholderia thailandensis E264. Med. Chem. Commun. 3, 976 (2012).

37. H. Chen, et al., Biosynthesis and engineering of the nonribosomal peptides with a C-1. terminal putrescine. Nat Commun 14, 6619 (2023).

38. H. Chen, et al., Biosynthesis of Glidomides and Elucidation of Different Mechanisms for Formation of β-OH Amino Acid Building Blocks. Angew Chem Int Ed 61, e202203591 (2022).

39. D. B. Hansen, S. B. Bumpus, Z. D. Aron, N. L. Kelleher, C. T. Walsh, The Loading Module of Mycosubtilin: An Adenylation Domain with Fatty Acid Selectivity. J. Am. Chem. Soc. 129, 6366–6367 (2007).

40. Y. Yuan, et al., Control of the polymyxin analog ratio by domain swapping in the nonribosomal peptide synthetase of Paenibacillus polymyxa. J Ind Microbiol Biotechnol 47, 551–562 (2020).

41. L. Zhong, et al., Engineering and elucidation of the lipoinitiation process in nonribosomal peptide biosynthesis. Nat Commun 12, 296 (2021).

42. C.-H. Ji, S. Park, K. Lee, H.-W. Je, H.-S. Kang, Lipidation Engineering in Daptomycin Biosynthesis. J. Am. Chem. Soc. 146, 30434–30442 (2024).

43. X. Wang, et al., Discovery of recombinases enables genome mining of cryptic biosynthetic gene clusters in Burkholderiales species. Proc. Natl. Acad. Sci. U.S.A. 115, E4255–E4263 (2018).

44. H. Chen, et al., Identification of Holrhizins E–Q Reveals the Diversity of Nonribosomal Lipopeptides in Paraburkholderia rhizoxinica. J. Nat. Prod. 83, 537–541 (2020).

45. J. Liu, et al., Rational construction of genome-reduced Burkholderiales chassis facilitates efficient heterologous production of natural products from proteobacteria. Nat Commun 12, 4347 (2021).

46. K.A.J. Bozhüyük, et al., Evolution-inspired engineering of nonribosomal peptide synthetases. Science 383, eadg4320 (2024).

47. K.A.J. Bozhüyük, et al., De novo design and engineering of non-ribosomal peptide synthetases. Nature Chem 10, 275–281 (2018).

48. K.A.J. Bozhüyük, et al., Modification and de novo design of non-ribosomal peptide synthetases using specific assembly points within condensation domains. Nat. Chem. 11, 653–661 (2019).

49. M. J. Calcott, J. G. Owen, D. F. Ackerley, Efficient rational modification of non-ribosomal peptides by adenylation domain substitution. Nat Commun 11, 4554 (2020).

50. R. He, et al., Knowledge-guided data mining on the standardized architecture of NRPS: Subtypes, novel motifs, and sequence entanglements. PLoS Comput Biol 19, e1011100 (2023).

51. X. Bian, et al., Direct Cloning, Genetic Engineering, and Heterologous Expression of the Syringolin Biosynthetic Gene Cluster in E. coli through Red/ET Recombineering. ChemBioChem 13, 1946–1952 (2012).

52. Q. Ouyang, et al., Promoter Screening Facilitates Heterologous Production of Complex Secondary Metabolites in Burkholderiales Strains. ACS Synth. Biol. 9, 457–460 (2020).

53. J. Yin, et al., Single-Stranded DNA-Binding Protein and Exogenous RecBCD Inhibitors Enhance Phage-Derived Homologous Recombination in Pseudomonas. iScience 14, 1– 14 (2019).

54. W. Niu, et al., Biosynthesis of Nonribosomal Peptides Chitinimides Reveal a Special Type of Thioesterase Domains. Chemistry 30, e202402763 (2024). 1.

55. A. E. Atik, M. Z. Guray, T. Yalcin, Observation of the side chain O -methylation of glutamic acid or aspartic acid containing model peptides by electrospray ionization-mass spectrometry. J Chromatogr B Analyt Technol Biomed Life Sci 1047, 75–83 (2017).

56. G. Pirri, A. Giuliani, S. Nicoletto, L. Pizzuto, A. Rinaldi, Lipopeptides as anti-infectives: a practical perspective. Open Life Sciences 4, 258–273 (2009).

57. S. Götze, P. Stallforth, Structure, properties, and biological functions of nonribosomal lipopeptides from pseudomonads. Nat. Prod. Rep. 37, 29–54 (2020).

58. R. H. Baltz, V. Miao, S. K. Wrigley, Natural products to drugs: daptomycin and related lipopeptide antibiotics. Nat Prod Rep 22, 717–741 (2005).

59. H. Chen, et al., Genomics-Driven Activation of Silent Biosynthetic Gene Clusters in Burkholderia gladioli by Screening Recombineering System. Molecules 26, 700 (2021).

60. K. L. Dunbar, et al., Resistance gene–guided genome mining reveals the roseopurpurins as inhibitors of cyclin-dependent kinases. Proc. Natl. Acad. Sci. U.S.A. 120, e2310522120 (2023).

61. Y. Wang, et al., Facile Method for Site-specific Gene Integration in Lysobacter enzymogenes for Yield Improvement of the Anti-MRSA Antibiotics WAP-8294A and the Antifungal Antibiotic HSAF. ACS Synth. Biol. 2, 670–678 (2013).

62. G.-L. Ma, et al., Biosynthesis of Tasikamides via Pathway Coupling and Diazonium-Mediated Hydrazone Formation. J. Am. Chem. Soc. 144, 1622–1633 (2022).

63. J. Rüschenbaum, W. Steinchen, F. Mayerthaler, A. Feldberg, H. D. Mootz, FRET Monitoring of a Nonribosomal Peptide Synthetase Elongation Module Reveals Carrier Protein Shuttling between Catalytic Domains. Angew Chem Int Ed 61, e202212994 (2022).

64. A. Tanovic, S. A. Samel, L.-O. Essen, M. A. Marahiel, Crystal Structure of the Termination Module of a Nonribosomal Peptide Synthetase. Science 321, 659–663 (2008).

