## Supporting Information for "Changing the initiation unit of nonribosomal peptide synthetases to access underexplored biosynthetic potential"

### Supplementary Materials and Methods

**Bacterial strains, reagents and growth conditions.** The bacterial strains, mutants and plasmids used and created in this study were listed in Supplementary Tables S1 and S2. Primers used in this research were listed in Supplementary Tables S3. All of *E. coli* strains were cultured in low-salt Luria - Bertani (LB) medium at 37°C or 30°C. The plasmid constructs in this study were obtained by linear- linear homologous recombination (LLHR) recombination in *E. coli* GB05-dir or linear-circular homologous recombination (LCHR) in *E. coli* GB08-red(1). Antibiotics were added to both LB medium plates and liquid at the following concentrations: chloramphenicol (Cm) 15 μg/mL or 10 μg/mL, kanamycin (Km) 15 μg/mL or 10 μg/mL, gentamycin (Genta) 6 μg/mL or 3 μg/mL, apramycin (Apra) 20 μg/mL or 10 μg/mL, ampicillin (Amp) 100 μg/mL or 50 μg/mL and tetracyline (Tet) 10 μg/mL or 5 μg/mL.

The strains of *Mycetohabitans rhizoxinica* HKI 454 (previous known as *Paraburkholderia rhizoxinica* HKI 454), *Caldimonas brevitalea* DSM 7029 (previous known as *Schlegelella brevitalea* DSM 7029) and *Burkholderia gladioli* ATCC 10248 were grown on CYMG medium at 30°C according to our previous work(2–4). The appropriate concentrations of antibiotics for *M. rhizoxinica* HKI 454 in liquid or plates were Kanamycin (Km) 30 μg/mL or 40 μg/mL, gentamycin (Genta) 20 μg/mL or 30 μg/mL and apramycin (Apra) 30 μg/mL or 50 μg/mL, respectively. The antibiotic of Apra of *C. brevitalea* DSM 7029 and *B. gladioli* ATCC 10248 in liquid or plates were 30 μg/mL or 40 μg/mL and 50 μg/mL or 300 μg/mL. The fermentation medium consists of M9 medium (Glucose 10 g/L, $K_2HPO_4$ 7 g/L, $KH_2PO_4$ 2 g/L, $(NH_4)_2SO_4$ 1 g/L, Sodium Citrate 0.5 g/L, $MgSO_4$ 0.1 g/L) for the mutants of *B. gladioli* ATCC 10248, low-salt LB medium for GB05-MtaA(5), and CYMG medium for *M. rhizoxinica* HKI 454 and *C. brevitalea* DSM 7029 at 30°C. The transformations of linear-DNA or plasmids into *M. rhizoxinica* HKI 454 and *C. brevitalea* DSM 7029 were performed by electroporation (Eppendorf® Electroporator 2510, 1300V) according to our previous works(6). The transformations of linear-DNA and plasmid containing gene cluster into *B. gladioli* ATCC 10248 were performed by electroporation and conjugation mediated methods, respectively(4).

PCR performed with ApexHF HS DNA Polymerase FS Master Mix (Accurate Biology, AG, cat. no. AG12202), and the resulting dsDNA was purified using the New Gel Mini Purification Kit (Beijing Zoman Biotechnology, cat. no. zpn202-3), following the manufacturer's instructions. Restriction enzymes and 1 kb DNA marker were obtained from New England Biolabs, while 5000 or 2000 DNA markers were acquired from AG. Antibiotics were purchased from MDBio, Inc., SIGMA, and Sangon (Shanghai), China.

**Bioinformatics analysis.** Firstly, some biosynthesis gene clusters of NRPSs derived from *Burkholderiales* were annotated by anti-SMASH(7) to provide detailed results of sequence analysis. The annotated initiation region sequence of NRPS was then uploaded to the"NRPS Motif Finder"database(8) for resolving the conserved motif and inter-motif structures in the NRPS. At the same time, using the free software MEGA11(9) for comparative analysis of homologous gene or protein sequences, align by ClustalW2. Finally, combined with the crystal structure of RzmA-Cs R148A/C8-CoA(10), as well as the current reports on the relevant sites of NRPS combinatorial biosynthesis, ten fusion sites were selected for the experiment.

**Direct cloning and heterologous expression of biosynthetic gene clusters.** The plasmids of *rzmA* and *holA* from *M. rhizoxinica* HKI 454, *chm* from *Chitinimonas koreensis* DSM 17726 were constructed by our group(3, 10). Firstly, BGCs of NRPSs derived from *Burkholderiales* were annotated by anti-SMASH and determine the boundaries of possible functional genes. The BGCs of *2A* (22.4 kb), *2C* (19.4 kb), *5C* (18.2 kb), *7C* (40.9 kb), *epyA* (34.4 kb) and *10C* (33.5 kb) from the genomic DNA of *M. rhizoxinica* HKI 454 were directly cloned via Red/ET recombineering in *E. coli* according to the previous protocols(11, 12) (Supplementary Tables S4).

The extracted genome DNA was digested by appropriated restriction enzymes, and the restriction enzymes of six BGCs were *Sca*I, *Xmn*I, *Sal*I-*Asc*I, *Asc*I, *Not*I and *Xba*I-*Avr*II in order. Then, the digested genomic DNA purified by ethanol precipitation, p15A vector with homologous arm (HAs: ~80 bp), T4 polymerase and buffer reacted in PCR program: 30°C, 60 min; 75°C, 20 min; 50°C, 30 min. The obtained PCR products then electroporated into *E. coli* GB05-dir that expressing the pSC101-BAD-ETgA-tet plasmid (L-arabinose induced) to resulting plasmids p15A-cm-*2A*, p15A-cm-*2C*, p15A-cm-*5C*, p15A-cm-*7C*, p15A-cm-*epy* and p15A-cm-*10C* (Supplementary Tables S1, S2 and S3).

In order to enhance the expression, the BGCs are under the regulation of exogenous constitutive promoters of $P_{Tn5\text{-}km}$ or $P_{Apra}$. A specific integration site *oriT*-phiC31-*attP* (only the BGC of *epy* use transposable element *oriT-tnpA-IR* cassette) was inserted by recombineering for the integration of gene clusters into the genome of heterologous hosts, resulting in constructs of plasmids p15A-phiC31-*attP*-apra-$P_{Tn5\text{-}km}$-*2A*, p15A-phiC31-*attP*-apra-$P_{Tn5\text{-}km}$-*2C*, p15A-phiC31-*attP*-apra-$P_{Tn5\text{-}km}$-*5C*, p15A-phiC31-*attP*-apra-$P_{Tn5\text{-}km}$-*7C*, p15A-cm-TnpA-km-$P_{13}$-$P_{Apra}$-*epyA* and p15A-phiC31-*attP*-apra-$P_{Tn5\text{-}km}$-*10C*, respectively (primers in Supplementary Tables S3, diagram in Fig. S2a). More detailed methods for direct cloning and engineering can be found in previous studies(1, 3, 5, 11, 13).

In this study, the heterologous hosts used as needed were *E. coli* GB05-MtaA(5), *C. brevitalea* DSM 7029Δ*glb*(3) and *B. gladioli* Δ*gbn*::*attB*(4). The series plasmids of *rzmA* were electroporated into the heterologous host of *E. coli* GB05-MtaA, and *epy*, *chm*, *2A*, *2C*, *3C*, *5C*, *7C* and *10C* were electroporated into *C. brevitalea* DSM 7029Δ*glb* for functional expression with reference to the previous method. Some modified 10C plasmids were transduced into 10248 in a binding transfer manner. Some modified plasmids like *10C* were transferred into chassis of *B. gladioli* Δ*gbn*::*attB* with method of conjugation mediated.

**Design and replacement of the initiation unit.**

**Cs domain swapping.** The Cs domain of the silent gene cluster in *M. rhizoxinica* HKI 454 was exchanged to the start of the *rzmA* gene cluster that had already obtained the activated product, and then fermentation detection was performed to check the product production. The experiment used the genome as a template, and the full-length Cs domain from the BGCs of *2C*, *3C*, *5C*, *epy* and *10C* were obtained by PCR, and overlap extension PCR was performed with TnpA-$P_{Tn5\text{-}km}$, and then the plasmids (p15A-TnpA-$P_{Tn5\text{-}km}$-2C/3C/5C/Epy/10C-Cs-RzmA) were obtained by LLHR recombineering with linearized p15A-cm-rzmA by *Sac*I and *Nde*I (primers in Tables S3, diagram in Fig. S3a). The resulting plasmids were validated and transformed into *C. brevitalea* DSM 7029Δ*glb* for heterologous expression and metabolic analysis. Detailed methods can be found in our previous studies of the Cs domain swapping(10).

**Replace the initiation regions of RzmA with M1~M10 of HolA.** The experiment was divided into two steps, the initiation regions of RzmA containing the Cs domain were replaced by amp-ccdB cassette, and then linearized DNA was obtained by digestion of *Pac*I. Ten sets of recombinant plasmids were conducted by LCHR or LLHR of recombineering. Detailed experimental process is as follows: Firstly, the amp-ccdB cassettes of three homology arms were LCHR recombineering with plasmid p15A-cm-rzmA to obtain three plasmids, p15A-*Pac*I-*amp-ccdB*-ΔCs-*Pac*I-*rzmA*, p15A-*Pac*I-*amp-ccdB*-ΔCsAT-*Pac*I-*rzmA* and p15A-*Pac*I-*amp-ccdB*-ΔCsAT-*Pac*I-*rzmA*, respectively. Secondly, the above three plasmids are digested by *Pac*I and purified to obtain linear DNA fragments p15A-ΔCs/ΔCsAT/ΔCsATC$_2$-RzmA, respectively. Then, the plasmid p15A-phiC31-*attP*-*apra*-P$_{Tn5\text{-}km}$-*holA* was used as a template for PCR amplification of ten fragments, *attP*-*apra*-P$_{Tn5\text{-}km}$-HolA-M1 to HolA-M10 (abbreviated as HM1 to HM10). Finally, approximately 600 ng of five fragments *attP*-*apra*-P$_{Tn5\text{-}km}$-HolA-HM2 to HM6 DNA and 300 ng of p15A-ΔCs-*rzmA* were electroporated into Ara-inducible recombinase-expressing GB05-dir for LLHR, resulting in p15A-phiC31-*attP*-*apra*-P$_{Tn5\text{-}km}$-RzmA-HM2 to HM6, respectively. LLHR recombineering between the DNA of HM7 to HM8 and p15A-ΔCsAT-RzmA was used to construct p15A-phiC31-*attP*-*apra*-P$_{Tn5\text{-}km}$-RzmA-HM7 and p15A-phiC31-*attP*-*apra*-P$_{Tn5\text{-}km}$-RzmA-HM8. Recombination of HM9 to HM10 and p15A-ΔCsAT-RzmA resulted in plasmids of p15A-phiC31-*attP*-*apra*-P$_{Tn5\text{-}km}$-RzmA-HM9 and p15A-phiC31-*attP*-*apra*-P$_{Tn5\text{-}km}$-RzmA-M10, respectively. P15A-phiC31-*attP*-*apra*-P$_{Tn5\text{-}km}$-RzmA -HM1 was obtained directly by LCHR of *attP*-*apra*-P$_{Tn5\text{-}km}$-HolA-M1 with p15A-*cm-rzmA*. Detailed description for the construction of *amp-ccdB* plasmids were performed according to our previous protocols(4, 14), and related strains/plasmids/primers were given in Supplementary Tables S1, S2 and S3. The resulting plasmids were validated and transformed into *E. coli* GB05-MtaA for heterologous expression and production comparison.

**Change the initiation region of BGC *5C* and *10C*.** The initiation regions of BGC *5C* were replaced by M5, M7, M8, and M9 of RzmA, respectively. The initiation regions of BGC *10C* were replaced by the initiation units of RzmA-M8 and HolA-M7, respectively. The detailed method is illustrated by BGC*10C*: the directly cloned plasmid p15A-*cm-10C* was digested with various enzymes, *Spe*I, *Apa*I and *Xmn*I (*Dra*I, *Sca*I and *Asc*I or *Sna*BI and *Bsr*GI for *5C* to obtain the linearized fragment p15A-*10C*-ΔCs-ΔA$_1$(750bp). Fragments *attP*-*apra*-P$_{Tn5\text{-}km}$-RM8 and *attP*-*apra*-P$_{Tn5\text{-}km}$-HM7 containing the corresponding homology arms were obtained by PCR in the same way as described above. P15A-phiC31-*attP*-*apra*-P$_{Tn5\text{-}km}$-RM8-10C and P15A-phiC31-*attP*-*apra*-P$_{Tn5\text{-}km}$-HM7-10C were obtained by subsequent LLHR recombineering (primers in Tables S3, diagram in Fig. S5b).

**Change the initiation region of BGC *epy* and *chm*.** The initiation module CsAT of Epy was replaced by the initiation unit M7 of RzmA; the initiation module ATC of gene cluster chm was replaced by M4 of Bgdd; The process of chm BGC as an example was as follows, firstly, the fragment Bgdd-Cs and promoter P$_{Apra}$ were obtained by PCR, and after that, overlapping extension PCR was performed to obtain the fragment P$_{Apra}$-BM4. Then, the original plasmid p15A-TnpA-*km-chm* was transferred into GB08-red. Finally, about 300 ng of DNA fragment P$_{Apra}$-BM4 was electroporated into Ara-inducible recombinase-expressing GB08-red/TnpA-*km-chm* for LCHR recombineering, which resulted in p15A-TnpA-km-P$_{Apra}$-Bgdd-Cs-Chm (strains/plasmids/primers in Supplementary Tables S1, S2 and S3, diagram in Fig. S4b). The resulting plasmids were validated and transformed into *C. brevitalea* DSM 7029Δ*glb* for heterologous expression and metabolic analysis.

***In situ* modification of the NRPS initiation units in *M. rhizoxinica* HKI 45.** Due to the high homology and multiple repeats among the gene clusters, we constructed plasmids containing

2-3 kb homology arms (HA), promoters and different exchange initiation units by LLHR recombineering or Gibson assembly, and then obtained linear DNA fragments by *Pac*I digested (Supplementary Fig.S5c). Homologous recombination occurred by electroporation of the 2μg linear DNA into *M. rhizoxinica* HKI 454 that induced recombinase expression. The correct mutants were confirmed through colony PCR and subsequently cultured in CYMG liquid medium for metabolic analysis. Detailed description for the editing of the HKI genome can be found in the previous protocols(6).

**Fermentation and HPLC-MS analysis of metabolic crude extracts.** The fermentation for BGCs in *E. coli* GB05-MtaA, *C. brevitalea* DSM 7029Δ*glb* and *B. gladioli* Δ*gbn*::*attB* were performed in low-salt LB, CYMG and M9 liquid medium, as described above. Seed cultures of heterologous expression mutants of different BGCs were incubated overnight in 1.5 mL tubes containing 1.3 mL liquid culture on a mixture Mix at 30°C 950 rpm. The seed culture was diluted at the ratio of 1:50 into a 250 mL flask containing 50 mL of LB broth at 30°C with 200 rpm shaking for 3 days. Then, the metabolites were adsorbed overnight for 18-24 hours by adding 2% of XAD-16 resin, and incubated continuously for an additional 24 hours. The resin and bacterial cells were then collected at 8300 rpm and the product extracted with 35 mL of methanol at 30°C, 200 rpm for 2 h. The methanol extract mixture was filtered through filter paper and evaporated to dryness and the crude extract was re-suspended in 1 mL MeOH for further HPLC analysis. To compare the secondary metabolites of different engineering mutants of 05-MtaA/ *rzmA*, the seed cultures were uniformly $OD_{600}$ values and transferred to 50 mL of fermentation medium containing antibiotics in a 250 mL shaker at 2% inoculation, with three parallel experimental groups in each.

The HPLC-MS was performed on an ODS column (Luna RP-C18, 4.6 × 250 mm, 5 μm, 0.3 mL/min) with gradient elution (solvent A, $H_2O$ + 0.1% FA, formic acid, solvent B, ACN, acetonitrile +0.1% FA), UV spectroscopy detection range 190-400 nm. The elution program was 25 min, 0-3 min, 5% B, 3-19 min, 5-95% B, 19-22 min, 95% B, 23-25 min, 5% B). HRMS detection was performed using a Bruker Impact HD High Resolution Q-TOF mass spectrometry (Bruker Daltonics, Bremen, Germany) with a standard ESI ion source in the range of 400-1600 m/z, in positive electrospray ionization mode, after the confirmation of the presence of product. The HRESIMS was performed on a short column (Thermo Scientific™ Acclaim™ 120 C18, 2.2 μm, 100 × 2.1 mm, 0.3 mL /min ) was used with an elution time and gradient of 0-3 min, 5% B, 3-19 min, 5-95% B, 19-22 min, 95% B, 23-25 min, 5% B.

**Purification and characterization of compounds.** The derivatives of rhizomide A were identified according to our previous description with minor modifications(6, 10). For compound **1**, the extract of mutants GB05-MtaA/HM1-rzmA to HM4-rzmA at retention time in 14.0 min, with molecular formula of $C_{41}H_{65}N_7O_{10}$ at *m/z* 816.4834 $[M+H]^+$ (calc 816.4866). Compound **2** (RT 13.5 min) at *m/z* 802.4689 $[M+H]^+$ (calc 802.4709) in the extract of mutants GB05-MtaA/RzmA-HM5 to RzmA-HM10, with chemical formulas of $C_{40}H_{63}N_7O_{10}$ (Fig. 2B and 2D, *SI Appendix*, Fig. S11). The *epy* BGC was functional expressed in heterologous host *C. brevitalea* DSM 7029 and produce the endopyrrole A. HRESIMS analysis compound **3** (RT 14.4 min) is the derivatives of endopyrrole A at *m/z* 1049.5896 $[M+H]^+$ (calc 1049.5918) in the extract of mutant 7029Δ*glb*/*epy*-RM7, with chemical formulas of $C_{54}H_{80}N_8O_{13}$ (column: Thermo Scientific™ Acclaim™ 120 C18, 2.2 μm, 100 × 2.1 mm, 0.3 mL /min; gradient elution: 0-3 min, 5% B, 3-19 min, 5-95% B, 19-22 min, 95% B, 23-25 min, 5% B) (Fig. 3B and C, *SI Appendix*,

Fig. S12).

For compound **4** and **6**, the crude extract of 8 L culture of mutant Δ*gbn*::*attB*/*5C*-RM8 was eluted by a normal-phase silica gel column resulting in fraction of $CH_2Cl_2$:MeOH = 20:1, which was further purified by semipreparative RP-HPLC (column: Agilent ZORBAX Stable Bond-C18; 5 μm, 9.4×250 mm; gradient elution: solvent A, $H_2O$ with 0.5% TFA, solvent B, ACN. 0-3 min, 40% B; 3-10 min, 40%-50%B; 10-20 min, 50%-62% B; 20.1 min, 72% B; 20.1-30 min, 72%-95% B; 30-40 min, 95% B, flow rate of 2 ml/min) to yield **4** (5 mg) at retention time 30 min and **6** (3 mg) at retention time 32 min, respectively. Using the same parameters in a ratio of $CH_2Cl_2$: MeOH = 10:1, we detected and purified the compound **5** (4 mg) at retention time 16 min.

For compound **7**, the extract of 8 L culture of mutant HKI 454/*10C*-RM8 was also isolated by normal phase silica gel column in fraction of $CH_2Cl_2$: MeOH = 10:1, which was further isolated and purified by semipreparative RP-HPLC. The parameters were as follows: solvent A, $H_2O$ with 0.5% TFA, solvent B, ACN. 0-3 min, 28% B; 3-20 min, 28%-45%B; 20-35 min, 45%-50% B; 35.1-42 min, 95% B to obtain **7** (4 mg) with retention time at 28 min. As for compound **10**, the cleaner crude extract was obtained by 8 L fermentation of mutant Δ*gbn*::attB/*10C*-HM7 in M9 medium, which can be directly separated and purified by semipreparative RP-HPLC. The gradient elution was as follows: 0-3 min, 30% B; 3-15 min, 30%-42%B; 15-30 min, 42%-50% B; 30-35 min, 50%-95% B; 35-40 min,95% B. At a flow rate of 2 mL/min, we obtained the compound **10** (5 mg) with retention time at 19 min.

For compound **11, 12** and **13**, the crude extract of 12 L culture of mutant DSM 7029Δ*glb*/BM4-*chm* was firstly eluted on a normal-phase silica gel column via different ratios of $CH_2Cl_2$: MeOH. The target compound is present in in fraction of 10:1 and 8:1, gradient elution of extracts was performed using semipreparative RP-HPLC (column: Thermo Fisher Basic 120-C18; 5 μm, 10×250 mm) with the program as follows: 0-3 min, 25% B; 3-15 min, 25%-50%B; 15-25 min, 50%-70% B; 25-35 min, 70%-75% B; 35-40 min, 70%-95% B; 40-45 min, 95% B, DAD at 254 nm to acquire compound **13** (5 mg) at 21.5 min, while compounds **11** and **12** were detected at 18 min and 19.5 min, respectively.

For compound **14** and **15**, the crude extract of 6 L culture of mutant HKI 454/*2C*-RM7 was de detected at ratios of $CH_2Cl_2$: MeOH = 5:1, which was further purified by RP-HPLC (column: Thermo Fisher Basic 120-C18; 5 μm, 10×250 mm; gradient elution: solvent A, $H_2O$ with 0.5% TFA, solvent B, ACN. 0-3 min, 30% B; 3-15 min, 30%-40%B; 15-30 min, 40%-44% B; 30-35 min, 44%-95% B; 35-40 min, 95% B) to yield **14** (8 mg) with retention time at 11.5 min and **15** (10 mg) at 20.5 min, respectively. Due to the low solubility of the product, it is difficult to show the signal in the $^1$H and $^{13}$C NMR spectrum with DMSO-$d_6$ as the solvent, HRESIMS (60 min program) *m/z* 530.3178 $[M+H]^+$ of **14** with retention time at 21.8 min and 558.3485 $[M+H]^+$ of **15** with retention time at 25.1 min, respectively.

**Marfey's analysis of the amino acid constituents of compounds.** Using Marfey's analysis to determine amino acid configuration of the product as reported previously(15). In brief, approximately 1 mg of the compound was dissolved in 500 μL of 6M HCl at 60°C overnight. The overnight reaction solutions were dried by nitrogen and redissolve with 200 μL of dd$H_2O$. Approximately 1 mg of standard amino acids (L- and D-) were also dissolved in 200 μL of dd$H_2O$, respectively. Then, added 25 μL of 1 M $NaHCO_3$ and 200 μL of 1-fluoro-2,4-dinitrophenyl-5-L-alanine amide (L-FDAA) solution to the above compounds and standard amino acids, and the mixture was vortexed and heated at 40 °C for 60 min. Finally, 100 μL of 2M HCl solution were added to quench the reaction, and the resulting mixture solution was filtered through a small 2.5 μm filter for HPLC-MS detection. Using the Luna RP-C18 column (4.6 × 250 mm, 5 μm) for LC–MS analyzed with a linear gradient of ACN and 0.1% aqueous formic acid solution with different elution conditions (0-3 min, 5% B, 3-19 min, 5-95% B, 19-22 min, 95% B, 23-25 min, 5% B or 0-3 min, 5%B, 3-38 min, 5%-40%B, 38.1 min, 95%B, 38.1-42 min, 95%B, 42.1 min, 5%B, 42.1 min-45 min, 5%B) at a flow rate of 0.3 mL/min and UV detection at 330 nm. Each chromatographic peak was identified by independent comparison of their retention time and molecular weight with the Marfey reagent derivative of the L- and D-amino acid standard (Supplementary Fig. S14).

**Biological Activity Screening.** The anti-inflammatory activity of the compounds was evaluated by measuring the inhibitory effect of the compounds on nitric oxide (NO) generation in LPS-stimulated RAW 264.7 macrophages using the classic Griess method(16). RAW 264.7 cells were seeded in a 96-well plate at $10^5$ cells/well and incubated for 24 h. Then, cells were treated with different concentrations of the compounds were added and incubated for 30 min, followed by LPS (10 mg/mL) and incubated for another 24 h. At this time, the cellular supernatant was tested for nitrite accumulation using a NO detection kit containing Griess reagent (Beyotime, batch number: S0021, Shanghai, China), which represents the cellular NO level, and the OD value of the absorbance at 540 nm on the microplate was read.

The cytotoxic activity of the new compounds was evaluated by the MTT method(6). Five cell lines were selected for detection: human liver cancer cell HepG2, human renal clear cancer cell ACHN, human non-small cell lung cancer cell H1299, human colon cancer cell HT-29, human non-small cell lung cancer cell A549, human gastric adenocarcinoma cell AGS and human cardiomyocyte cell AC-16 were transferred to 96-well plates for inoculation, with approximately 4 to 5 × $10^4$ cells in each well, and incubated overnight at 37°C in 5% CO2/air. When the cells grew and adhered to the wall, different concentrations of the compound to be tested were added to each well, with a solution volume of 100 μL in each well, and incubated in a cell culture incubator for 24 h. The negative control was DMSO with the same concentration, and the positive control was doxorubicin. 20 μL of MTT (5 mg/mL) was added to each well and incubated for another 4 h. After the supernatant was aspirated, 150 μL of DMSO was added to each well and allowed to stand in an incubator overnight. The OD value (vertical axis) of each well was detected at a wavelength of 570 nm using an enzyme-labeled instrument, and a correlation curve was drawn with the compound concentration (horizontal axis) to calculate the corresponding $IC_{50}$. Each experimental group had three parallels, and the experiment was repeated three times.

### Supplementary Tables

**Table S1.** The strains and mutants used in this study

| Strains | Descriptions | Sources |
|---|---|---|
| ***E. coli*** | | |
| GB2005 | HS996, Δ*rec*ET, Δ*ybc*C, used for retransformation of plasmids to avoid background. | Our lab(17) |
| GB05-dir | GB2005, *ara*C-BAD-*ETgA*. Used for LLHR. | Our lab(1) |
| GB05-red | GB2005, *ara*C-BAD-*γαβA*. Used for LCHR. | Our lab(1) |
| GB05-red-gyrA462 | GB2005-red with GyrA mutation of Arg462Cys. Used for LCHR which contained *amp-ccdB* cassette. | Our lab(14) |
| GB2005-dir-pSC101-BAD-*ETgA*-tet | Generally used for direct cloning of gene clusters. Contains the araC-BAD-ETgA expression cassette for LLHR, together with the temperature-sensitive recombinase plasmid pSC101-BAD-ETgA-tet | Our lab(1, 11) |
| GB05-MtaA | GB2005 carries MtaA, a phosphopantetheine transferase from *Stigmatella aurantiaca*. Used for heterologous expression of *rzmA* and *holA* gene clusters and related mutants. | Our lab(5) |
| WM 3064 | The donor strain for conjugate transfer with *Burkholderia gladioli* ATCC10248. | Our lab(18) |
| 05 MtaA/$P_{Tn5\text{-}km}$-*rzmA* | *rzmA* gene cluster integrated into GB05-MtaA | This work |
| 05 MtaA/$P_{Tn5\text{-}km}$-*holA* | *holA* gene cluster integrated into GB05-MtaA | This work |
| 05 MtaA/HM1-*rzmA* | Swapping N terminal of RzmA-Cs with HolA-Cs | This work |
| 05 MtaA/HM2-*rzmA* | Swapping RzmA-Cs with HolA-Cs (short linker) | This work |
| 05 MtaA/HM3-*rzmA* | Swapping RzmA-CsXU with HolA-CsXU (W: N) | This work |
| 05 MtaA/HM4-*rzmA* | Swapping RzmA-Cs with HolA-Cs (full-length linker) | This work |
| 05 MtaA/HM5-*rzmA* | Swapping RzmA-CsA with HolA-CsA (short linker) | This work |
| 05 MtaA/HM6-*rzmA* | Swapping RzmA-CsA with HolA-CsA (full-length linker) | This work |
| 05 MtaA/HM7-*rzmA* | Swapping RzmA-CsAT module with HolA-CsAT (full-length linker) | This work |
| 05 MtaA/HM8-*rzmA* | Swapping N terminal of RzmA-$C_2$ with HolA-$C_2$ (CsAT-$C_2$N) | This work |
| 05 MtaA/HM9-*rzmA* | Swapping RzmA-CsAT$C_2$XU with HolA- CsAT$C_2$XU | This work |
| 05 MtaA/HM10-*rzmA* | Swapping RzmA-CsAT$C_2$ with HolA- CsAT$C_2$ (full-length linker) | This work |
| ***Burkholderiales*** | | |
| *Caldimonas brevitalea* DSM 7029 (Δ*glb*-attB) | Used for heterologous expression of *epyA* and *chm* gene clusters and related mutants. | Our lab(3) |
| 7029Δ*glb*/*epy* | *epy* gene cluster integrated into 7029Δ*glb* with $P_{13}$, $P_{Apra}$ and oriT-TnpA-km. | This work |
| 7029Δ*glb*/*epy*-RM7 | Replace Ep*y*-CAT with RzmA-Cs*AT in 7029Δ*glb*. | This work |

| Strains | Descriptions | Sources |
|---|---|---|
| 7029Δ*glb*/ 2C-Cs-RzmA | Replace RzmA-Cs (RM4) with 2C-Cs in 7029Δ*glb*. | This work |
| 7029Δ*glb*/ 3C-Cs-RzmA | Replace RzmA-Cs (RM4) with 3C-Cs in 7029Δ*glb*. | This work |
| 7029Δ*glb*/ 5C-Cs-RzmA | Replace RzmA-Cs (RM4) with 5C-Cs in 7029Δ*glb*. | This work |
| 7029Δ*glb*/ 10C-Cs-RzmA | Replace RzmA-Cs (RM4) with 10C-Cs in 7029Δ*glb*. | This work |
| 7029Δ*glb*/ EpyD-Cs-RzmA | Replace RzmA-Cs (RM4) with EpyD-Cs in 7029Δ*glb*. | This work |
| 7029Δ*glb*/ $P_{Tn5\text{-}km}$-*2A* | *2A* gene cluster integrated into 7029Δ*glb* with $P_{Tn5\text{-}km}$ and oriT-phiC31-*attP*. | This work |
| 7029Δ*glb*/ $P_{Tn5\text{-}km}$-*10C* | *10C* gene cluster integrated into 7029Δ*glb* with $P_{Tn5\text{-}km}$ and oriT-phiC31-*attP*. | This work |
| 7029Δ*glb*/ $P_{Tn5\text{-}km}$-*5C* | *5C* gene cluster integrated into 7029Δ*glb* with $P_{Tn5\text{-}km}$ and oriT-phiC31-*attP*. | This work |
| 7029Δ*glb*/ $P_{Tn5\text{-}km}$-*7C* | *7C* gene cluster integrated into 7029Δ*glb* with $P_{Tn5\text{-}km}$ and oriT-phiC31-*attP*. | This work |
| 7029Δ*glb*/RM7-2A | Replace 2A-AT with RzmA-Cs*AT in 7029Δ*glb*. | This work |
| 7029Δ*glb*/RM7-2C | Replace 2C-CsAT with RzmA-Cs*AT in 7029Δ*glb*. | This work |
| 7029Δ*glb*/RM7-3C | Replace 3C-CsAT with RzmA-Cs*AT in 7029Δ*glb*. | This work |
| 7029Δ*glb*/RM5-5C | Replace 5C-CsA with RzmA-Cs*A in 7029Δ*glb* (short linker). | This work |
| 7029Δ*glb*/RM7-5C | Replace 5C-CsAT with RzmA-Cs*AT in 7029Δ*glb*. | This work |
| 7029Δ*glb*/RM8-5C | Replace 5C-CsATC$_2$N with RzmA-Cs*ATC$_2$N in 7029Δ*glb*. | This work |
| 7029Δ*glb*/RM9-5C | Replace 5C-CsATC$_2$XU with RzmA-Cs*ATC$_2$XU in 7029Δ*glb*. | This work |
| 7029Δ*glb*/RM7-7C | Replace 7C-CsAT with RzmA-Cs*AT in 7029Δ*glb*. | This work |
| 7029Δ*glb*/HM7-10C | Replace 10C-CsAT-C$_2$AT with HolA-CsAT in 7029Δ*glb*. | This work |
| 7029Δ*glb*/BM4-Chm | Replace Chm-ATC$_2$ with Bgdd-Cs (full-length linker) in 7029Δ*glb*. | This work |
| *B. gladioli* Δ*gbn*::attB | Used for heterologous expression of *waps* gene clusters and related mutants. | Our lab(4) |
| Δ*gbn*::attB /RM8-5C | Replace 5C-CsATC$_2$N with RzmA-Cs*ATC$_2$N inΔ*gbn*::attB. | This work |
| Δ*gbn*::attB /HM7-10C | Replace 10C-CsAT-C$_2$AT with HolA-CsAT in Δ*gbn*::attB. | This work |
| *Mycetohabitans rhizoxinica HKI 454* (Δ*rhi*) | An Endosymbiont of Rhizopus microspores, DSM 19002. The original producer of rhizomide, holrhizin and endopyrrole. | Our lab(2, 6) |
| HKI 454/Epy-RM7 | Replace Ep*y*-CAT with RzmA-Cs*AT in HKI 454. | This work |

| **Strains** | **Descriptions** | **Sources** |
|---|---|---|
| HKI 454/2A-RM7 | Replace 2A-AT with RzmA-Cs*AT in HKI 454. | This work |
| HKI 454/2C-RM7 | Replace 2C-CsAT with RzmA-Cs*AT in HKI 454. | This work |
| HKI 454/3C-RM7 | Replace 3C-CsAT with RzmA-Cs*AT in HKI 454. | This work |
| HKI 454/5C-RM7 | Replace 5C-CsAT with RzmA-Cs*AT in HKI 454. | This work |
| HKI 454/5C-RM8 | Replace 5C-CsATC$_2$N with RzmA-Cs*ATC$_2$N in HKI 454. | This work |
| HKI 454/7C-RM7 | Replace 7C-CsAT with RzmA-Cs*AT in HKI 454. | This work |
| HKI 454/10C-RM7 | Replace10C-CsATC$_2$AT with RzmA-Cs*AT in HKI 454. | This work |
| HKI 454/10C-RM8 | Replace 10C-CsATC$_2$ATC$_3$N with RzmA-Cs*ATC$_2$N in HKI 454. | This work |
| *Chitinimonas koreensis* DSM 17726 | The original producer of chitinimides. | Our lab(3) |

**Table S2.** Plasmids used in this study

| Plasmids | Descriptions | Sources |
|---|---|---|
| pBR322-amp-ccdB-rpsL | Template for amplification of DNA fragment pBR322-amp-ccdB | Our lab(11, 14) |
| p15A-cm-*rzmA* | The plasmid contains *rzmA* gene cluster with p15A vector and $cm^R$. | Our lab(10) |
| p15A-cm-*holA* | The plasmid contains *holA* gene cluster with p15A vector and $cm^R$. | Our lab(10) |
| pBAC-cm | Template for amplification vector pBAC-cm | Our lab(11) |
| p15A-phiC31-attP-apra-$P_{Tn5\text{-}km}$-*rzmA** | Template for amplification of *oriT*-phiC31-*attP* and $P_{Tn5\text{-}km}$-RzmA-Cs* (R148A) | Our lab(10) |
| p15A-phiC31-attP-apra-$P_{Tn5\text{-}km}$-*holA* | Template for amplification of *oriT-attP*-phiC31 and $P_{Tn5\text{-}km}$-HolA-Cs | Our lab(10) |
| p15A-TnpA-$P_{Tn5\text{-}km}$-2C-Cs-RzmA | Swapping 2C-Cs with RzmA-Cs to constructed recombinant *rzmA* BGC | This work |
| p15A-TnpA-$P_{Tn5\text{-}km}$-3C-Cs-RzmA | Swapping 3C-Cs with RzmA-Cs to constructed recombinant *rzmA* BGC | This work |
| p15A-TnpA-$P_{Tn5\text{-}km}$-5C-Cs-RzmA | Swapping 5C-Cs with RzmA-Cs to constructed recombinant *rzmA* BGC | This work |
| p15A-TnpA-$P_{Tn5\text{-}km}$-EpyD-Cs-RzmA | Swapping EpyD-Cs with RzmA-Cs to constructed recombinant *rzmA* BGC | This work |
| p15A-TnpA-$P_{Tn5\text{-}km}$-10C-Cs-RzmA | Swapping 10C-Cs with RzmA-Cs to constructed recombinant *rzmA* BGC | This work |
| p15A-*Pac*I-amp-ccdB-ΔCs-*Pac*I-*rzmA* | Linearized ΔCs-RzmA was obtained for the construction of plasmid R-HM2-M6 | This work |
| p15A-*Pac*I-amp-ccdB-ΔCsAT-*Pac*I-*rzmA* | Linearized ΔCsAT-RzmA was obtained for the construction of plasmid R-HM7-M8 | This work |
| p15A-*Pac*I-amp-ccdB-$ΔCsATC_2$-*Pac*I-*rzmA* | Linearized $ΔCsATC_2$-*rzmA* was obtained for the construction of plasmid R-HM9-M10 | This work |
| p15A-attP-apra-$P_{Tn5\text{-}km}$-HM1-*rzmA* (HM1-rzmA) | Swapping N terminal of RzmA-Cs with N terminal of HolA-Cs to constructed recombinant *rzmA* gene cluster | This work |
| p15A-attP-apra-$P_{Tn5\text{-}km}$-HM2-*rzmA* (HM2-rzmA) | Swapping RzmA-Cs with HolA-Cs (short linker) to constructed recombinant *rzmA* BGC | This work |
| p15A-attP-apra-$P_{Tn5\text{-}km}$-HM3-*rzmA* (HM3-rzmA) | Swapping RzmA-CsXU with HolA-CsXU (W: N) to constructed recombinant *rzmA* BGC | This work |
| p15A-attP-apra-$P_{Tn5\text{-}km}$-HM4-*rzmA* (HM4-rzmA) | Swapping RzmA-Cs with HolA-Cs (full-length linker) to constructed recombinant *rzmA* BGC | This work |
| p15A-attP-apra-$P_{Tn5\text{-}km}$-HM5-*rzmA* (HM5-rzmA) | Swapping RzmA-CsA with HolA-CsA (short linker) to constructed recombinant *rzmA* BGC | This work |
| p15A-attP-apra-$P_{Tn5\text{-}km}$-HM6-*rzmA* (HM6-rzmA) | Swapping RzmA-CsA with HolA-CsA (full-length linker) | This work |

| Plasmids | Descriptions | Sources |
|---|---|---|
| p15A-attP-apra-$P_{Tn5\text{-}km}$-HM7-*rzmA* (HM7-rzmA) | Swapping RzmA-CsAT module with HolA-CsAT (full-length linker) | This work |
| p15A-attP-apra-$P_{Tn5\text{-}km}$-HM8-*rzmA* (HM8-rzmA) | Swapping N terminal of RzmA-$C_2$ with HolA-$C_2$ (CsAT-$C_2$N) | This work |
| p15A-attP-apra-$P_{Tn5\text{-}km}$-HM9-*rzmA* (HM9-rzmA) | Swapping RzmA-CsAT$C_2$XU with HolA-CsAT$C_2$XU | This work |
| p15A-attP-apra-$P_{Tn5\text{-}km}$-HM10-*rzmA* (HM10-rzmA) | Swapping RzmA-CsAT$C_2$ with HolA- CsAT$C_2$ (full-length linker) | This work |
| p15A-cm-*epyA* | *epyA* gene cluster by direct cloning from *M. rhizoxinica* HKI 454 genomic DNA (digested by *Not*I) with BAC replicon, and $cm^R$. | This work |
| p15A-cm-TnpA-km-$P_{13}$-$P_{Apra}$-*epyA* | Added *IR-oriT-TnpA-IR-km* cassette, $P_{13}$ from *C. brevitalea* DSM 7029 and $P_{Apra}$ | This work |
| p15A-cm-TnpA-km-$P_{13}$-$P_{Apra}$-RM7-*epy* (RM7-epy) | Swapping EpyD-CsAT with RzmA-CsAT to constructed recombinant *epy* BGC | This work |
| p15A-cm-*Pac*I-*5C*HA1-apra-$P_{Tn5\text{-}km}$-RM7-*5C*HA2(long)-*Pac*I | Construct plasmids contain a 2-3 kb homology arm for replace 5C-CsAT with RzmA-Cs*AT in HKI 454 (Digested by *Pac*I) | This work |
| p15A-cm-*Pac*I-*10C*HA1-apra-$P_{Tn5\text{-}km}$-RM8-*10C*HA2(long)-*Pac*I | Construct plasmids contain a 2-3 kb homology arm for replace 10C-CsAT$C_2$N with RzmA-CsAT$C_2$N in HKI 454 (Digested by *Pac*I) | This work |
| p15A-cm-*2A* | *2A* gene cluster by direct cloning from *M. rhizoxinica* HKI 454 genomic DNA (digested by *Sca*I) with p15A vector, and $cm^R$. | This work |
| p15A-phiC31-attP-apra-$P_{Tn5\text{-}km}$-*2A* | *2A* BGC with *oriT-attP*-phiC31 and $P_{Tn5\text{-}km}$ | This work |
| p15A-attP-apra-$P_{Tn5\text{-}km}$-*2A*-RM7 | Swapping 2A-AT with RzmA-Cs*AT to constructed hybrid *2A* | This work |
| p15A-cm-*2C* | *2C* gene cluster by direct cloning from *M. rhizoxinica* HKI 454 genomic DNA (digested by *Xmn*I) with p15A vector, and $cm^R$. | This work |
| p15A-phiC31-attP-apra-$P_{Tn5\text{-}km}$-*2C* | *2C* BGC with *oriT-attP*-phiC31 and $P_{Tn5\text{-}km}$ | This work |
| p15A-cm-*3C* | Direct cloning of *3C* gene cluster from *M. rhizoxinica* HKI 454 (digested by *Ase*I) with p15A vector, and $cm^R$. | This work |
| p15A-cm-*5C* | *5C* gene cluster by direct cloning from *M. rhizoxinica* HKI 454 genomic DNA (digested by *Sal*I and *Asc*I) with p15A vector, and $cm^R$. | This work |

| **Plasmids** | **Descriptions** | **Sources** |
|---|---|---|
| p15A-phiC31-attP-apra-$P_{Tn5\text{-}km}$-*5C* | *5C* gene cluster with *oriT*-*attP*-phiC31 and $P_{Tn5\text{-}km}$ | This work |
| p15A-phiC31-attP-apra-$P_{Tn5\text{-}km}$-*5C*-RM5 | Swapping 5C-CsA (short linker) with RzmA-Cs*A to constructed recombinant *5C* | This work |
| p15A-phiC31-attP-apra-$P_{Tn5\text{-}km}$-*5C*-RM7 | Swapping 5C-CsAT with RzmA-Cs*AT to constructed recombinant *5C* | This work |
| p15A-phiC31-attP-apra-$P_{Tn5\text{-}km}$-*5C*-RM8 | Swapping 5C-CsATC$_2$N with RzmA-Cs*ATC$_2$N to constructed recombinant *5C* | This work |
| p15A-phiC31-attP-apra-$P_{Tn5\text{-}km}$-*5C*-RM9 | Swapping 5C-CsATC$_2$XU with HolA-CsATC$_2$XU to constructed recombinant *5C* | This work |
| p15A-cm-*7C* | *7C* gene cluster by direct cloning from *M. rhizoxinica* HKI 454 genomic DNA (digested by *Asc*I) with p15A vector, and cm$^R$. | This work |
| p15A-phiC31-attP-apra-$P_{Tn5\text{-}km}$-*7C* | *7C* gene cluster with *oriT*-*attP*-phiC31 and $P_{Tn5\text{-}km}$ | This work |
| p15A-phiC31-attP-apra-$P_{Tn5\text{-}km}$-*7C*-RM7 | Swapping 7C-CsAT with RzmA-Cs*AT to constructed recombinant *7C* | This work |
| p15A-cm-*10C* | *10C* gene cluster by direct cloning from *M. rhizoxinica* HKI 454 genomic DNA (digested by *Xba*I and AvrII) with p15A vector, and cm$^R$. | This work |
| p15A-phiC31-attP-apra-$P_{Tn5\text{-}km}$-*10C* | *10C* gene cluster with *oriT*-*attP*-phiC31 and $P_{Tn5\text{-}km}$ | This work |
| p15A-phiC31-attP-apra-$P_{Tn5\text{-}km}$-*10C*-HM7 | Swapping 10C-CsATC$_2$AT with HolA-CsAT to constructed recombinant *10C* | This work |
| p15A-TnpA-km-*chm* | p15A replicon, km$^R$, with o*riT*-*TnpA*-*IR* cassette and *chm* gene cluster | Our lab(3) |
| p15A-TnpA-km-$P_{Apra}$-*chm* | Under the control of constitutive promotor $P_{Apra}$ | Our lab(3) |
| p15A-TnpA-km-$P_{Apra}$-*bgdd*-Cs-*chm* (C-BM4) | Swapping Chm-ATC with Bgdd-Cs (full-length) to constructed recombinant *chm* | This work |

**Table S3.** Primers used in this study

| Primers | Sequence (5'-3') | Application |
|---|---|---|
| p15A-TnpA-HA1 (Reused) | **CTGAGGTCATTACTGGATCTATCAACAGGAGTCCAAGCGAGCTCGATATC**TGCATCCGATGCAAGTGTGT | For construction of p15A-TnpA-$P_{Tn5-km}$-2C-Cs-RzmA |
| TnpA-$P_{Tn5-km}$-RBS-2 (Reused) | AATCTGTACCTCCTTAAGTCAGAAGAACTCGTCAAGAAG | |
| $P_{Tn5-km}$-2C-Cs-HA1 | **ACGAGTTCTTCTGACTTAAGGAGGTACAGATT**ATGCCTGCTTGCGTTGCCCC | |
| 2C-Cs-rzmA-HA-2 | **ATGAGCTGGTGTGCCAAGCGGTTGGCCTGCGCATTCAGTTGCGTATAGCT**CACCTGGTGAATGCATTGGTGCG | |
| rzmA-A1-check | AATCGCCAGCAGGCCGACGAC | |
| $P_{Tn5-km}$-3C-Cs-HA1 | **GAGTTCTTCTGACTTAAGGAGGTACAGATT**ATGTCTGATAGCATCACATCGTCT | For construction of p15A-TnpA-$P_{Tn5-km}$-3C-Cs-RzmA |
| 3C-Cs-rzmA-HA-2 | **ATGAGCTGGTGTGCCAAGCGGTTGGCCTGCGCATTCAGTTGCGTATAGCT**AACCATCTGGTGCACGCACC | |
| $P_{Tn5-km}$-5C-Cs-HA1 | **GAGTTCTTCTGACTTAAGGAGGTACAGATT**ATGGATGTCAGCGTCATGCTCACT | For construction of p15A-TnpA-$P_{Tn5-km}$-5C-Cs-RzmA |
| 5C-Cs-rzmA-HA-2 | **ATGAGCTGGTGTGCCAAGCGGTTGGCCTGCGCATTCAGTTGCGTATAGCT**CAACTGGTGAATGCATAAGTGCGA | |
| $P_{Tn5-km}$-epy-Cs-HA1 | **GAGTTCTTCTGACTTAAGGAGGTACAGATT**ATGGATGCTAGCATCACGCCCG | For construction of p15A-TnpA-$P_{Tn5-km}$-epy-Cs-RzmA |
| epy-Cs-rzmA-HA-2 | **ATGAGCTGGTGTGCCAAGCGGTTGGCCTGCGCATTCAGTTGCGTATAGCT**GCAGATCGCCACCCGCGTGT | |
| $P_{Tn5-km}$-10C-Cs-HA1 | **CGAGTTCTTCTGACTTAAGGAGGTACAGATT**ATGGCTGTTAGCGCGACACCC | For construction of p15A-TnpA-$P_{Tn5-km}$-10C-Cs-RzmA |
| 10C-Cs-rzmA-HA-2 | **ATGAGCTGGTGTGCCAAGCGGTTGGCCTGCGCATTCAGTTGCGTATAGCT**CAATTGGTGAATACACCAGTGCGAT | |
| rzmA-CsAT-KO-amp-ccdB-HA1 | **ATCTCTTCAAATGTAGCACCTGAAGTCAGCCCCATACGATATAAGTTGTT**TTAATTAATTTGTTTATTTTTCTAAATAC | For construction of p15A-amp-ccdB-CsAT-KO-RzmA |
| rzmA-CsAT-KO-amp-ccdB-HA2 | **AACGATAGCGGCAAATGGCCTTCGCGTGAGACCGGTGTGATCTCGGGCAG**TTAATTAATTATATTCCCCAGAACATCAG | |
| p15A-phiC31-attP-HA1 (Reused) | **ATCTCTTCAAATGTAGCACCTGAAGTCAGCCCCATACGATATAAGTTGTT**TGATCATATGCGGATTAGAAAAACA | For construction of p15A-attP-$P_{Tn5-km}$-HM1~HM5-RzmA |
| HolA-M1-rzmA-HA2 | **ACCAATTAGCGCAGTGCTTGAGCCAATATGCTTCGTCTTGTGCTCGTTGGGCGGA**AGCTCGGTATTGCGCATCG | |
| HolA-M2-rzmA-HA2 | **TGAGCAAATTATGGTTTGTCGCCGAATACCCACCAAACGACAGATCAAGA**TCAAACGGCATCACGTTGACTGTG | |
| HolA-M3-rzmA-HA2 | **AAACAATTGGTGGACACACAGATGCGCTGGATAATCGCGCTGTGTTGCATT**CCATTCGATCAGCAGTCGATGCC | |
| HolA-M4-rzmA-HA2 | **GCTCGATGAGCTGGTGTGCCAAGCGGTTGGCCTGCGCATTCAGTTGCGTATAGCT**GAGCGTTTGCTCCTCGTAG | |
| HolA-M5-rzmA-HA2 | **CGATCGAGCTTGCCGTTGGGTGTCAGCGGCAACGCATCGAGCCGCACAAA**TGCACTGGGCACCATGTACTCGG | |

| Primers | Sequence (5'-3') | Application |
|---|---|---|
| HolA-M6-rzmA-HA2 | **ATGAGCTGGTGTGCCAAGCGGTTGGCCTGCGCATTCAGTTGCGTATAGCT**GAGCGTTTGCTCCTCGTAGACCAA | For construction of p15A-attP-$P_{Tn5\text{-}km}$-HM6~HM10-RzmA |
| HolA-M7-rzmA-HA2 | **ACACGTTCGACACCCAATAGCTCGGCCCAGATCGTCGCGAGCGTCGTCTC**GAGCTCGCCTTGGGGTGCTTCATA | |
| HolA-M8-rzmA-HA2 | **GCCAGCGTCGCGCGCCAATAGTCGCTTTGTGCGTTAAGCCGCTCGTCCGT**GAGCCACTGACGCTGCCACGCTG | |
| HolA-M9-rzmA-HA2 | **TCGAACAACTGGTGCACGCACAAGTGCGACGGATAGTCCCGCTGCGTCGC**GTTCCACGTCTGCAGCAACAACG | |
| HolA-M10-rzmA-HA2 | **GATCAACTGGTGCGCGAGGCGATTAGCTTGTGCGTTCAGCTGCGCATAGCT**GAGCGTTTGCTCCTCGTAGACCA | |
| 5C-apra-RM7-HA1 | **ATCAAAGCCGTCGTGGTCCCTGAAGCGCAACGATAGATTCTGATTATCGTATTAATATTTATTCGTACACCTATTAAACATCAGGCCGCTCGGTCGCATGAATCGGAGCTGTGTT**TCATCTCGTTCTCCGCTCATGAG | p15A-cm-*Pac*I-*5C*HA1-apra-$P_{Tn5\text{-}km}$-RM7-*5C*HA2(long)-*Pac*I construction |
| 5C-RM7-HA2 | **GAACCACAGCCGCTGTTGTGCGAACGACAGCGGCAAACTACCTTTGCGAGACACCGGTGTGATCTCGGA**CAGCGTATCGGTGCCTTGTT | |
| P15A-PacI-5C-2 | **TTGCGCTTCAGGGACCACGACGGCTTTGAT**TTAATTAAGACGTCGATATCTGGCGA | |
| 5C-longHA-PacI-M7-2 | **TGCCTGGAGATCCTTAAGATCTTAA**TTAATTAAACAAGCCAACGGTGAGCCACAAG | |
| P15A-1 | GATCTTAAGGATCTCCAGGCA | |
| 5C-M7-1 | TCCGAGATCACACCGGTGTCTCGCAAA | |
| 10C-apra-RM8-HA1 | **CAGGCCGCTGACGTACAATATGCCCCGAACCTTGACGGTACCTCAATACGTGGGGGAGGGGCGAACCACGTATCTTAAATCAAATAAATCTTTTATCTAAATTGGAGATAAATTG**TCATCTCGTTCTCCGCTCATGAG | p15A-cm-*Pac*I-*10C*HA1-apra-$P_{Tn5\text{-}km}$-RM8-*10C*HA2(long)-*Pac*I construction |
| 10C-RM8-HA2 | **CAGCAACACCGGGGCACCGGCCAGCTGCGTGCGCCAGTACGCGCTTTGCGCCTCAAGCCGCTCACCCGT**GAGCCACTGCCGTTGCCACG | |
| P15A-PacI-10C-2 | **GGCATATTGTACGTCAGCGGCCTG**TTAATTAAGACGTCGATATCTGGCGAAAATGA | |
| 10C-longHA-PacI-p15A-2 | **TGCCTGGAGATCCTTAAGATC**TTAATTAATTCGTCATCGGCCACTTGAATGCCGCA | |
| P15A-1 | GATCTTAAGGATCTCCAGGCA | |
| 10C-M8-1 | ACGGGTGAGCGGCTTGAGGCGCAAAGC | |
| p15A-cm-epy-HA1 | **CGCCAGCCAGATACCGAGCCCATCATACGACGAGCACCTTCATGCGCGTCGAGTCACGGTTAGCAAACAGATAGGCGTGATCTTAAG**GATCTCCAGGCA | For direct cloning of *epy* gene cluster |
| p15A-cm-epy-HA2 | **GCAACTGCTGGGCTTGCTTGTCGGTCAGCCCGAACTGCTGCATCAGTTGCGCATAGATCGCCGCCTCCAGCTTCGGAT**TAAGACGTCGATATCTGGCGA | |

| Primers | Sequence (5'-3') | Application |
|---|---|---|
| cm-$P_{13}$-epy-HA1 | **ACAAAAGCACGGAGTTTCACCAAAGTGTCAGACATGGGCTTCTCCTTTAA**GGTTATGTGTGGGAGGGCTAA | To construct plasmid of P15A-TnpA-km-cm-$P_{13}$-Epy |
| cm-2 | GACGTTGATCGGCACGTAAG | |
| $p_{13}$-cm-1 | **CTTACGTGCCGATCAACGTC**ATGTATATCTCCTGGCTCGCTCCTTGGGAAAACC | |
| cm-$P_{13}$-epy-HA2 | **ACGAGCGCAGGAGCAAAGAGGCAAAATTTTTTGAGTTTGTGAGCATCCAT**CGGCCATTATCATCTGCCTCGGCGTTGACACTC | |
| $P_{Apra}$-epy-HA1 | **CAGGTGCTAAGCGCTGCGGTTACCTGAATCTTTGCAGCGGCTCATGACAG**ACGCTCAGTGGAACGAGGTT | Construction of $P_{Apra}$ for promoter insertion |
| $P_{Apra}$-epy-HA2 | **GTTTGGGCAGTGGATAATGCATACGTAACGGGCGTGATGCTAGCATCCAT**AATCTGTACCTCCTTAAGTCAG | |
| apra-2 | AATCTGTACCTCCTTAAGTCAG | |
| $P_{Apra}$-RzmA-M7-HA1 | **CTGACTTAAGGAGGTACAGATT**ATGGACGCTAGCGTCATGTCCACT | |
| RzmA-M7-epy-HA2 | **AGCTGCGCGAGAAACCACAACCGCTGCTGCGCGAACGATAGCGGCAGACT**TTCGCGTGAGACCGGTGTGATCT | |
| P15A-cm-2A-HA1 | **CTTGATCGAGTCGGCAGTCAGTGGGCCTGCCGTGGCGCAAACGGCCGAACAACAGCAATCCAGTCACGGCGATTTCCC**GATCTTAAGGATCTCCAGGCA | For direct cloning of *2A* gene cluster |
| P15A-cm-2A-HA2 | **GTGCGCGAGATTCAAGGCCATTTGCGGGAACTGTATGGCCTTCAGGTGTCGCCCGATCTGATTTCGAGCGTCACCGAC**TAAGACGTCGATATCTGGCGA | |
| P15A-cm-2C-HA1 | **GATGCCAATGCTCAAATTCACCTTGTCGGACCGAGGGTCAACCTGGAAATCCTCGTTGAGCGACAGGATCGGATCACC**GATCTTAAGGATCTCCAGGCA | For direct cloning of *2C* gene cluster |
| P15A-cm-2C-HA2 | **CATCGATTGGTGGCTCATCGATGCTTGGCCCAGCTCAAACCAGTTGGCCGCTTCTTCAGCCAGTACGCTGGGTGCAAG**TAAGACGTCGATATCTGGCGA | |
| P15A-cm-5C-HA1 | **GGGGCTGAACTGCACCTGCGTCGATCAGCCGGTCATCTCGACGATGCAACGTGCGCGGGCCGGCGATTGCTTCGTCGC**GATCTTAAGGATCTCCAGGCA | For direct cloning of *5C* gene cluster |
| P15A-cm-5C-HA2 | **TCCACGTCAATCGCGCGGTGTCGGCGACGGCAGTGTTTCGAAGTCCCAAGCGATACTGGCATACGCACGTGCGGCCCG**TAAGACGTCGATATCTGGCGA | |
| P15A-cm-7C-HA1 | **TATCCTCGTGCGTAGTGCAGCAAACAACGTTGTTAAAAGCTTTGTGAAGTTTTCAATATGCTGCGCTGGCTGGTCAGG**GATCTTAAGGATCTCCAGGCA | For direct cloning of *7C* gene cluster |
| P15A-cm-7C-HA2 | **CGACACGCACCGGTGAGGATCCGACGGGGGACATTCGTTCGCTGCCGCGATAGCATAGGCATGAGCTCGCTAGAGCGC**TAAGACGTCGATATCTGGCGA | |

| Primers | Sequence (5'-3') | Application |
|---|---|---|
| P15A-cm-10C-HA1 | **TATTTGATACGCATTGCCTTGCCCTAGCTGATCCAGGAACCATAGACGCTCTTGCGCGAAAGACAAAGGCAGTGCATC**GATCTTAAGGATCTCCAGGCA | For direct cloning of *10C* gene cluster |
| P15A-cm-10C-HA2 | **GCGGACCGGATTGAAGGATGCCTGTTCGGCAACGGCGAGCGCACGGGTAACGTCGATCTCGTGACACTCGCGCTGAAC**TAAGACGTCGATATCTGGCGA | |
| p15A-phiC-apra-HA1 | **ATCTCTTCAAATGTAGCACCTGAAGTCAGCCCCATACGATATAAGTTGTT**TGATCATATGCGGATTAGAAAAACA | To construct plasmid of p15A-phiC31-apra-$P_{Tn5-km}$-2A |
| p15A-phiC-apra-$P_{Tn5-km}$-2A-HA2 | **CAGTCAGTTTGGGACGTACTAAGCGGAACTGGCATACTGTAAACAGACATAATCTGTACCTCCTTAAG**TCAGAAGAACTCGTCAAGAAG | |
| p15A-phiC-apra-HA1 | Repeat | To construct plasmid of p15A-phiC31-apra-$P_{Tn5-km}$-2C |
| p15A-phiC-apra-$P_{Tn5-km}$-2C-HA2 | **GTTTGCGCGGTGGATAACGGATGGGTGAGTGGGGCAACGCAAGCAGGCATAATCTGTACCTCCTTAAG**TCAGAAGAACTCGTCAAGAAG | |
| p15A-phiC-apra-$P_{Tn5-km}$-5C-HA2 | **GTTTGAGCAGCGGATAATGCATACGTAGTGAGCATGACGCTGACATCCATAATCTGTACCTCCTTAAG**TCAGAAGAACTCGTCAAGAAG | To construct plasmid of p15A-phiC31-apra-$P_{Tn5-km}$-5C |
| p15A-phiC-apra-$P_{Tn5-km}$-7C-HA2 | **GTTTGGGCAGTGGATAACGGATACGTAAGTGGCGTGACGCTAATAGACATAATCTGTACCTCCTTAAG**TCAGAAGAACTCGTCAAGAAG | To construct plasmid of p15A-phiC31-apra-$P_{Tn5-km}$-7C |
| p15A-phiC-apra-$P_{Tn5-km}$-10C-HA2 | **GTGCCGATATTGCCCGAGCCGATAATGGCGACCTTCACCTTTTGCGTCATAATCTGTACCTCCTTAAG**TCAGAAGAACT | To construct plasmid of p15A-phiC31-apra-$P_{Tn5-km}$-10C |

The homology arms are in bold. The sites of restriction enzyme are in red.

**Table S4.** Information of gene clusters involved in this study

| Gene clusters | Size (kb) | Accession numbers of core gene | Strains | Chromosome or plasmid |
|---|---|---|---|---|
| *bgdd* | | BM43_RS04185 | *B. gladioli* ATCC 10248 (4, 19) | NZ_CP009322.1 |
| *rzmA* | 23.0 | RBRH_RS12370 | *M. rhizoxinica* HKI 454 (2, 6, 10) | pBRH01 NC_014718.1 |
| *holA* | 19.7 | RBRH_RS16800 | | |
| *2A* | 18.6 | RBRH_RS12510 | | |
| *2C* | 13.5 | RBRH_RS12725 | | |
| *3C* | 7.5 | RBRH_RS13085 | | |
| *5C* | 10.3 | RBRH_RS13905 | | |
| *7C* | 30.0 | RBRH_RS21175 | | |
| *epy* | 26.2 | RBRH_RS19750 | | NC_014722.1 |
| *10C* | 16.6 | RBRH_RS08665 | | |
| *chm* | 35.5 | NZ_ATZZ00000000.1 | *C. koreensis DSM 17726* (3) | shotgun genome |

**Table S5.** The $^1$H (600 MHz) and $^{13}$C NMR (150 MHz) data of compound **4** in $CH_3DO$ and compound **5** and **6** in $CDCl_3$

| | no | 4 | | 5 | | 6 | |
|---|---|---|---|---|---|---|---|
| | | $\delta_C$ | $\delta_H$ ($J$ in Hz) | $\delta_C$ | $\delta_H$ ($J$ in Hz) | $\delta_C$ | $\delta_H$ ($J$ in Hz) |
| FA | 1 | 176.62, C | | 171.75, C | | 174.2, C | |
| | 2 | 36.83, $CH_2$ | 2.16, m | 22.38, $CH_3$ | 1.71, s | 36.2, $CH_2$ | 1.98, m |
| | 3 | 27.04, $CH_2$ | 1.55, m | | | 25.5, $CH_2$ | 1.50, m[a] |
| | 4 | 30.44, $CH_2$ | 1.26, m[a] | | | 29.2, $CH_2$ | 1.25, m[a] |
| | 5 | 30.24, $CH_2$ | 1.25, m[a] | | | 28.9, $CH_2$ | 1.25, m[a] |
| | 6 | 33.00, $CH_2$ | 1.22, m[a] | | | 31.7, $CH_2$ | 1.25, m[a] |
| | 7 | 23.78, $CH_2$ | 1.25, m[a] | | | 22.6, $CH_2$ | 1.25, m[a] |
| | 8 | 14.54, $CH_3$ | 0.85, t (7.05) | | | 14.0, $CH_3$ | 0.86, t (7.0) |
| Leu | 1 | 175.42, C | | 173.3, C | | | |
| | 2 | 53.84, CH | 4.38, dd (6.18, 9.18) | 52.1, CH | 4.51, m | 172.6, C | 4.42, m |
| | 3a | 41.73, $CH_2$ | 1.55, m[a] | 40.9, $CH_2$ | 1.46, m[a] | 51.9, CH | 1.50, m[a] |
| | 3b | | | | 1.57, m[a] | 40.8, $CH_2$ | 1.62, m[a] |
| | 4 | 26.14, CH | 1.64, m | 24.8, CH | 1.55, m[a] | 24.8, CH | 1.62, m[a] |
| | 5 | 22.26, $CH_3$ | 0.92, d (6.54) | 22.2, $CH_3$ | 0.89, d (5.88) | 22.1, $CH_3$ | 0.92, d (6.24) |
| | 6 | 23.35, $CH_3$ | 0.96, d (6.54) | 22.6, $CH_3$ | 0.91, d (6.00) | 22.9, $CH_3$ | 0.95, d (6.42) |
| | NH | | | | 6.65, brs | | 5.78, brs |
| Val | 1 | 173.34, C | | 172.0, C | | 170.6, C | |
| | 2 | 59.50, CH | 4.26, d (5.25) | 59.0, CH | 4.26, brs | 58.4, CH | 4.34, m |
| | 3 | 31.95, CH | 2.02, m | 29.7, CH | 2.11, m | 29.8, CH | 2.32, m |
| | 4 | 17.71, $CH_3$ | 0.69, d (6.84) | 17.8, $CH_3$ | 0.80, m[a] | 17.0, $CH_3$ | 0.82, d (6.9) |
| | 5 | 19.92, $CH_3$ | 0.76, d (6.84) | 19.4, $CH_3$ | 0.82, m[a] | 19.5, $CH_3$ | 0.89, d (6.84) |
| | NH | | | | 7.56, brs | | 6.68, brs |
| Trp | 1 | 175.36, C | | 174.5, C | | 172.2, C | |
| | 2 | 54.71, CH | 4.73, dd (4.8, 9.24) | 52.9, CH | 4.83, brs | 52.3, CH | 4.88, m |
| | 3a | 28.64, $CH_2$ | 3.19, dd (9.18, 14.94) | 26.9, $CH_2$ | 3.26, m | 27.3, $CH_2$ | 3.26 dd (5.34,14.94) |
| | 3b | | 3.37, dd (4.92, 14.82) | | | | 3.35 dd (6.18,14.94) |
| | 4 | 111.34, C | | 109.8, C | | 109.9, C | |
| | 5 | 124.63, CH | 7.05, m[a] | 123.4, CH | 6.98, brs | 123.2, CH | 6.95, s |
| | 6 | 128.83, C | | 127.4, C | | 127.4, C | |
| | 7 | 119.35, CH | 7.55, d (7.92) | 118.6, CH | 7.55 d (7.74) | 118.6, CH | 7.49, d (7.92) |
| | 8 | 119.90, CH | 6.99, m | 119.6, CH | 7.05, t (7.8) | 119.9, CH | 7.11, m |
| | 9 | 122.51, CH | 7.07, m[a] | 122.1, CH | 7.13, t (7.41) | 122.2, CH | 7.18, m |
| | 10 | 112.37, CH | 7.30, d (8.1) | 111.3, C | 7.31, d (7.98) | 111.3, CH | 7.36, d (11.88) |
| | 11 | 138.20, C | | 136.1, C | | 136.1, C | |
| | 2-NH | | | | 7.56, m[a] | | 6.78, d (6.72) |
| | 5-NH | | | | 8.64, brs | | 8.39, brs |
| OMe | 1 | | | | | 52.38, $CH_3$ | |

[a] overlapped

**Table S6.** The $^{1}H$ (600 MHz) and $^{13}C$ NMR (150 MHz) data of **7** and **10** in DMSO-$d_6$

| | no | **7** | | | no | **10** | |
|---|---|---|---|---|---|---|---|
| | | $\delta_C$ | $\delta_H$ ($J$ in Hz) | | | $\delta_C$ | $\delta_H$ ($J$ in Hz) |
| Octanoyl | 1 | 172.8, C | | Octanoyl | 1 | 172.4, C | |
| | 2 | 35.6, $CH_2$ | 2.11, m | | 2 | 35.2, $CH_2$ | 2.15, m[a] |
| | 3 | 25.8, $CH_2$ | 1.48, m[a] | | 3 | 25.4, $CH_2$ | 1.47, m[a] |
| | 4 | 28.9, $CH_2$ | 1.23, m[a] | | 4 | 28.4, $CH_2$ | 1.23, m[a] |
| | 5 | 28.9, $CH_2$ | 1.23, m[a] | | 5 | 28.5, $CH_2$ | 1.23, m[a] |
| | 6 | 31.7, $CH_2$ | 1.22, m[a] | | 6 | 31.2, $CH_2$ | 1.22, m[a] |
| | 7 | 22.5, $CH_2$ | 1.25, m[a] | | 7 | 22.0, $CH_2$ | 1.25, m[a] |
| | 8 | 14.4, $CH_3$ | 0.85, m[a] | | 8 | 13.9, $CH_3$ | 0.84, m[a] |
| Leu | 1 | 172.5, C | | Val | 1 | 171.1, C | |
| | 2 | 51.4, CH | 4.33, m | | 2 | 57.8, CH | 4.18, m[a] |
| | 3 | 40.7, $CH_2$ | 1.45, m[a] | | 3 | 30.0, CH | 1.97, m |
| | 4 | 24.7, CH | 1.58, m[a] | | 4 | 18.2, $CH_3$ | 0.82, m[a] |
| | 5 | 21.9, $CH_3$ | 0.83, m[a] | | 5 | 19.3, $CH_3$ | 0.84, m[a] |
| | 6 | 23.5, $CH_3$ | 0.86, m[a] | | NH | | 7.87, d (8.94) |
| | NH | | 8.03, m[a] | | | | |
| Val | 1 | 171.23, C | | Val | 1 | 170.8, C | |
| | 2 | 57.5, CH | 4.20, m | | 2 | 57.2, CH | 4.21, m[a] |
| | 3 | 31.4, CH | 1.98, m[a] | | 3 | 30.8, CH | 1.98, m[a] |
| | 4 | 18.3, $CH_3$ | 0.82, m[a] | | 4 | 17.9, $CH_3$ | 0.82, m[a] |
| | 5 | 19.6, $CH_3$ | 0.85, m[a] | | 5 | 19.1, $CH_3$ | 0.84, m[a] |
| | NH | | 7.57, d (8.7) | | NH | | 7.63, d (8.76) |
| Glu | 1 | 173.7, C | | Glu | 1 | 171.5, C | |
| | 2 | 52.2, CH | 4.13, m[a] | | 2 | 51.6, CH | 4.15, m[a] |
| | 3a | 27.5, $CH_2$ | 3.26, m | | 3a | 27.0, $CH_2$ | 1.78, m |
| | 3b | | 1.96, m[a] | | 3b | | 1.96, m[a] |
| | 4 | 31.7, $CH_2$ | 2.21, m | | 4 | 31.5, $CH_2$ | 2.21, m[a] |
| | 5 | 171.9, C | 8.14, d (7.1) | | 5 | 173.1, C | |
| | NH | | | | NH | | 8.14, d (7.44) |
| Lys | 1 | 174.2, C | | Lys | 1 | 173.6, C | |
| | 2 | 52.1, CH | 4.16, m[a] | | 2 | 51.5, CH | 4.16, m[a] |
| | 3a | 31.0, $CH_2$ | 1.55, m[a] | | 3a | 30.5, $CH_2$ | 1.55, m[a] |
| | 3b | | 1.70, m | | 3b | | 1.70, m |
| | 4 | 22.8, $CH_2$ | 1.34, m[a] | | 4 | 22.4, $CH_2$ | 1.33, m |
| | 5 | 27.0, $CH_2$ | 1.52, m[a] | | 5 | 26.6, $CH_2$ | 1.51, m[a] |
| | 6 | 39.1, $CH_2$ | 2.77, t (7.41) | | 6 | 38.6, $CH_2$ | 2.76, t (7.5) |
| | | | | | 2-NH | | 8.06, d (7.86) |

[a] overlapped

**Table S7.** The $^1$H (600 MHz) and $^{13}$C NMR (150 MHz) data of compound **13** in DMSO-$d_6$

| no | **13** | | no | **13** | |
|---|---|---|---|---|---|
| | $\delta_C$ | $\delta_H$ ($J$ in Hz) | | $\delta_C$ | $\delta_H$ ($J$ in Hz) |
| 1 | 172.1, C | | 24 | 169.8, C | |
| 2 | 51.0, CH | 4.63, m[a] | 25 | 56.5, CH | 4.21, q (5.5) |
| 3 | 30.0, $CH_2$ | 2.11, m | 26 | 61.3, $CH_2$ | 3.72, m |
| 4 | 29.9, $CH_2$ | 2.51, m[a] | 27 | 164.9, C | |
| 5 | 14.5, $CH_3$ | 2.04, s | 28 | 130.9, C | |
| 6 | 160.7, C | | 29 | 126.6, CH | 6.29, q (7.0) |
| 7 | 149.0, C | | 30 | 12.9, $CH_3$ | 1.67, d (7.1) |
| 8 | 124.1, CH | 8.19, s | 31 | 171.4, C | |
| 9 | 173.9, C | | 32 | 43.7, $CH_2$ | 2.35, m |
| 10 | 58.4, CH | 5.33, dd (2.4, 8.1) | 33 | 67.8, CH | 3.83, m |
| 11 | 31.5, $CH_2$ | 2.18, m[a] | 34 | 37.0, $CH_2$ | 1.38, m |
| | | 2.26, m | 35 | 25.0, $CH_2$ | 1.24, m[a] |
| 12 | 24.0, $CH_2$ | 1.98, m | | | 1.35, m[a] |
| | | 2.04, m[a] | 36 | 28.7, $CH_2$ | 1.23, m[a] |
| 13 | 46.8, $CH_2$ | 3.79, m | 37 | 29.0, $CH_2$ | 1.21, m[a] |
| 14 | 170.7, C | | 38 | 31.3, $CH_2$ | 1.22, m[a] |
| 15 | 50.4, CH | 4.59, m[a] | 39 | 22.1, $CH_2$ | 1.25, m[a] |
| 16 | 28.1, $CH_2$ | 1.59, m[a] | 40 | 14.0, $CH_3$ | 0.85, t (7.0) |
| | | 1.76, m | 1′ | 52.1, $CH_3$ | 3.65, s |
| 17 | 24.8, $CH_2$ | 1.52, m | 2-NH | | 8.63, d (8.0) |
| 18 | 40.4, $CH_2$ | 3.08, m | 15-NH | | 7.55, d (7.7) |
| 19 | 156.7, C | | 18-NH | | 7.58, brs |
| 20 | 163.5, C | | 21-NH | | 8.97, s |
| 21 | 129.9, C | | 25-NH | | 7.84, d (6.3) |
| 22 | 130.3, CH | 6.51, q (7.0) | 28-NH | | 9.33, s |
| 23 | 13.0, $CH_3$ | 1.62, d (7.1) | | | |

[a] overlapped

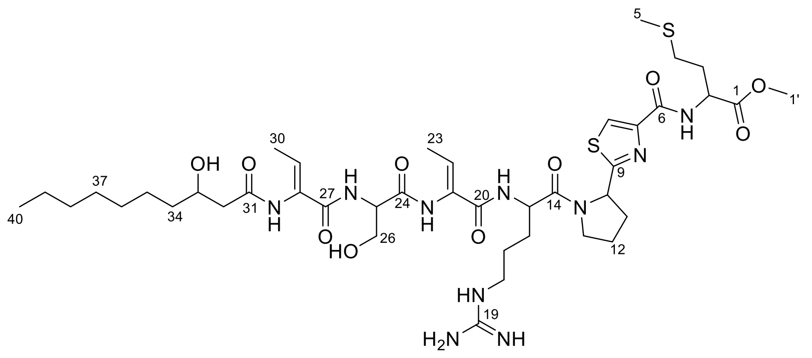

**Table S8.** $IC_{50}$ values of compounds **11-13** toward seven cancer cell lines (μM)

| Compounds | **HepG2** | **ACHN** | **H1299** | **HT-29** | **A549** | **AGS** | **AC-16** |
|---|---|---|---|---|---|---|---|
| **11** | > 80 | 52.43 | > 80 | > 80 | > 80 | > 80 | > 80 |
| **12** | 8.36 | > 80 | > 80 | > 80 | 12.02 | 32.72 | > 80 |
| **13** | > 80 | > 80 | > 80 | > 80 | 43.92 | > 80 | > 80 |
| chitinimide A | 39.11 | > 80 | 56.44 | > 80 | > 80 | > 80 | > 80 |
| Doxorubicin | 0.96 | 1.13 | 1.97 | 1.53 | 0.96 | 1.21 | 0.43 |

HepG2: Human liver cancer cell; ACHN: Human renal clear cancer cell; H1299: Human non-small cell lung cancer cell; HT-29: Human colon cancer cell; A549: Human non-small cell lung cancer cell; AGS: Human gastric adenocarcinoma cell; AC-16: Human cardiomyocyte Cell. The compound chitinimide A and Doxorubicin are the original product and drug of positive control, respectively.

### Supplementary Figures

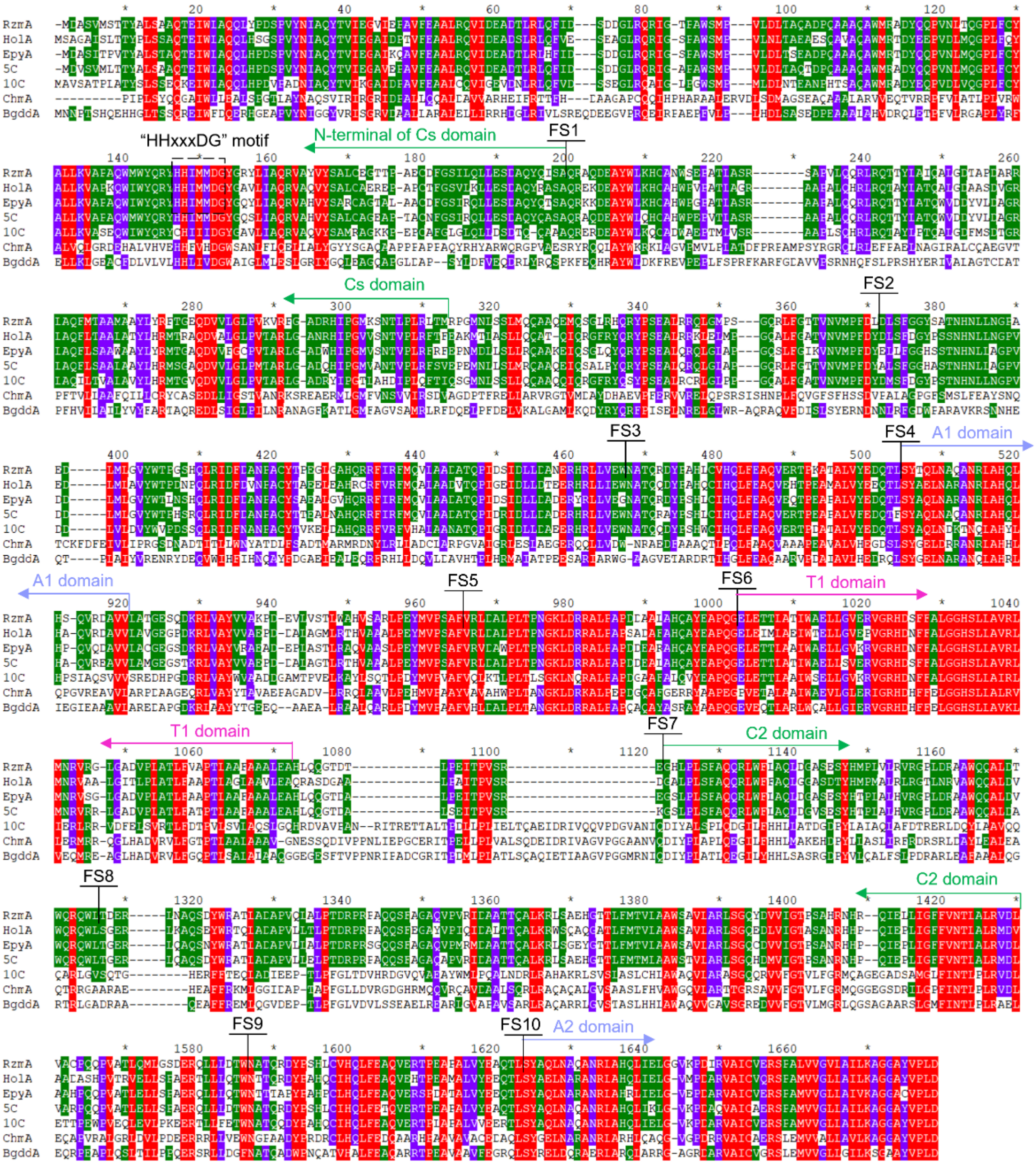

**Fig. S1.** The sequence alignment of the NRPS initiation module used in this study. Here, shows the location of ten sets of fusion sites (FS1-FS10). The sequences alignment of initiation regions including the Cs, $A_1$ domain and $T_1$ domain of the first module, the C2 domain and part of the $A_2$ domain of the second module (the *chm* BGC does not contain the Cs domain, and the sequences are the $C_2$, $A_2$, $T_2$ domains of the second module, the $C_3$ domain and part of the $A_3$ domain of the third module). This figure was obtained by sequence alignment of MAGE and modification with GeneDoc.

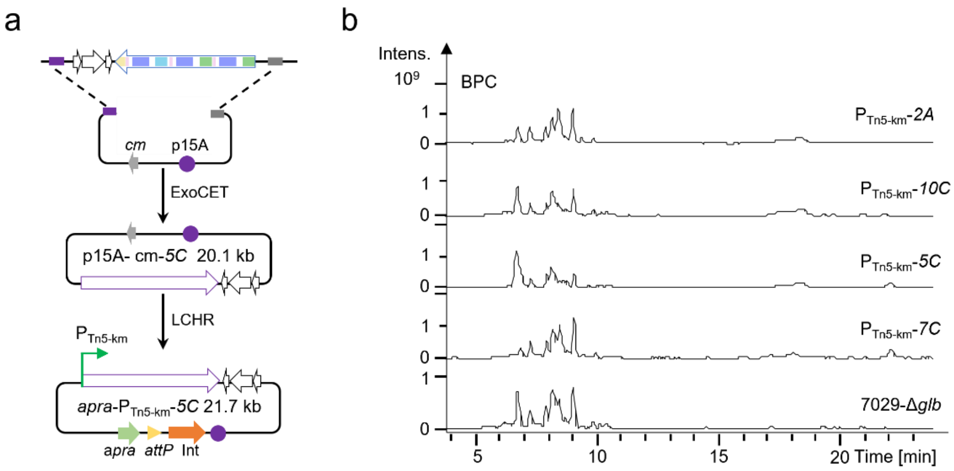

**Fig. S2**. Direct cloning and heterologous expression of silent NRPS from HKI 454. a. Schematic of direct cloning of gene clusters with promoter insertion (*5C* as an example). b. HPLC-MS analysis crude extracts of wild-type with heterologous expression of mutants.

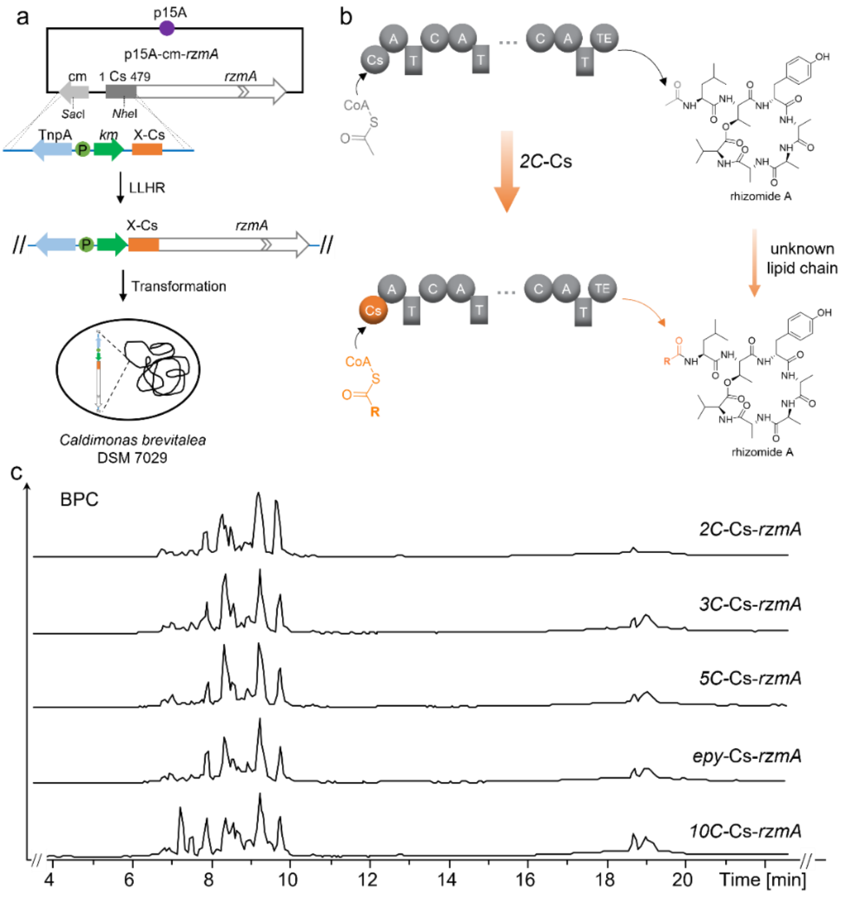

**Fig. S3.** Functional validation of the initiation module of the silent BGCs in HKI 454. a. Swapping the full-length Cs domain of RzmA with those Cs domain from silent NRPS, using recombineering in *E. coli*. The resulting constructs were transformed into DSM 7029 for heterologous expression. b. Schematic diagram of Cs domain exchange and synthesis of recombinant products. c. HPLC-MS analysis of fermentation crude extracts of constructs with different Cs domain exchanges.

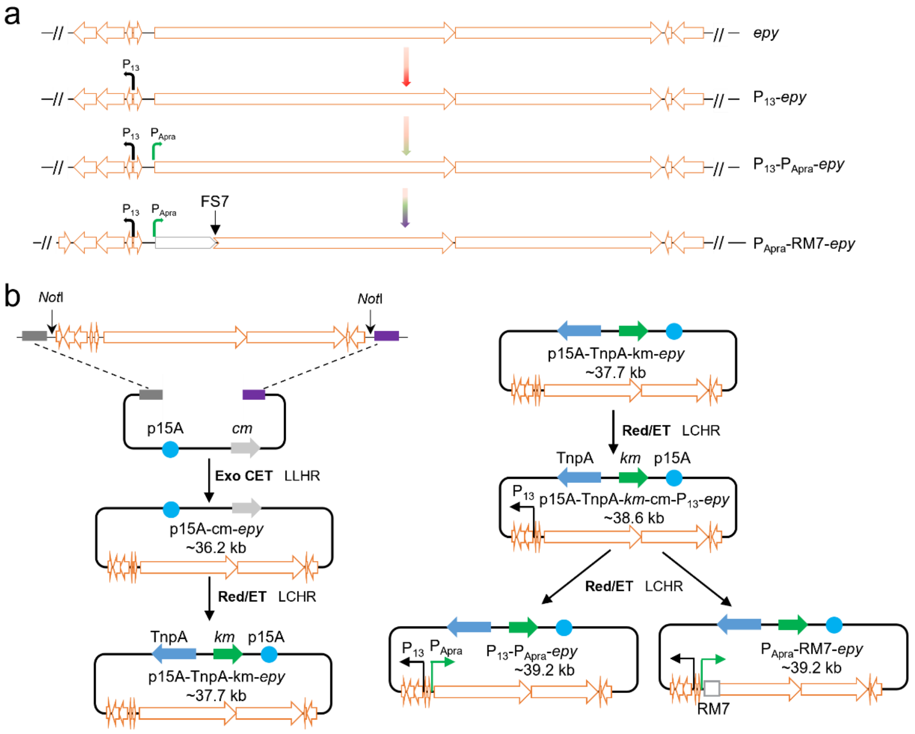

**Fig. S4**. Diagram of direct cloning and initiation module modification of the *epy* gene cluster. a. Promoter insertion and RM7 module substitution in *epy* BGC. b. Diagram of direct cloning and initiation module changing of *epy* BGC. *Not*I for genome digested, p15A-cm as vector, insertion of TnpA-km, insertion of the $P_{13}$ promoter before *epyC*, the original promoter of the core gene *epyD* was replaced with $P_{Apra}$, which was engineered with M7 from RzmA.

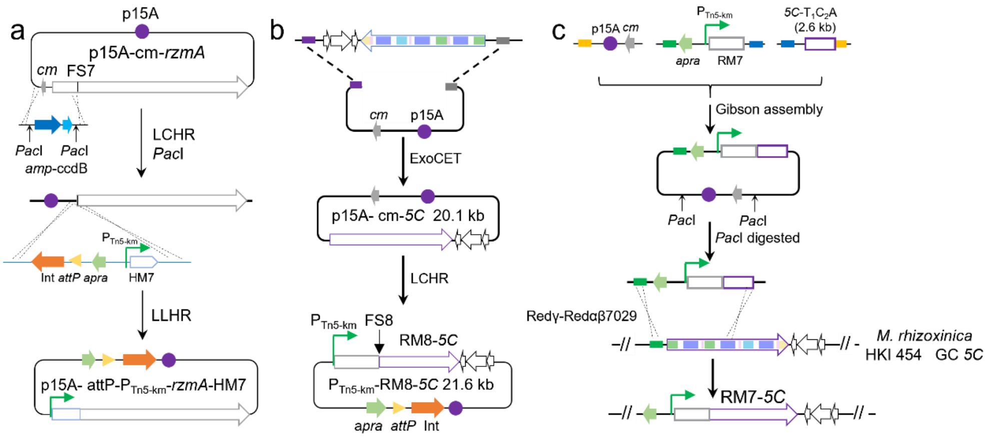

**Fig. S5**. Diagram of changing initiation module of cryptic NRPS gene clusters from HKI 454. a. Diagram for swapping of RzmA with HolA initiation modules in *E. coli*, taking HM7 changing based on LLHR as an example. b. Diagram for changing the initiation modules of silent NRPS BGCs in *E. coli*, direct cloning has been completed. Take the exchange of the 5C initiation region with RzmA-M8 as an example. c. Schematic showing initiation module substitution in HKI 454. Plasmids with long homology arms were constructed *in vitro*, enzyme digested to obtain linear recombinant DNA, and electroporated into HKI 454 for homologous recombineering. Take the exchange of the 5C initiation region with RzmA-M7 as an example.

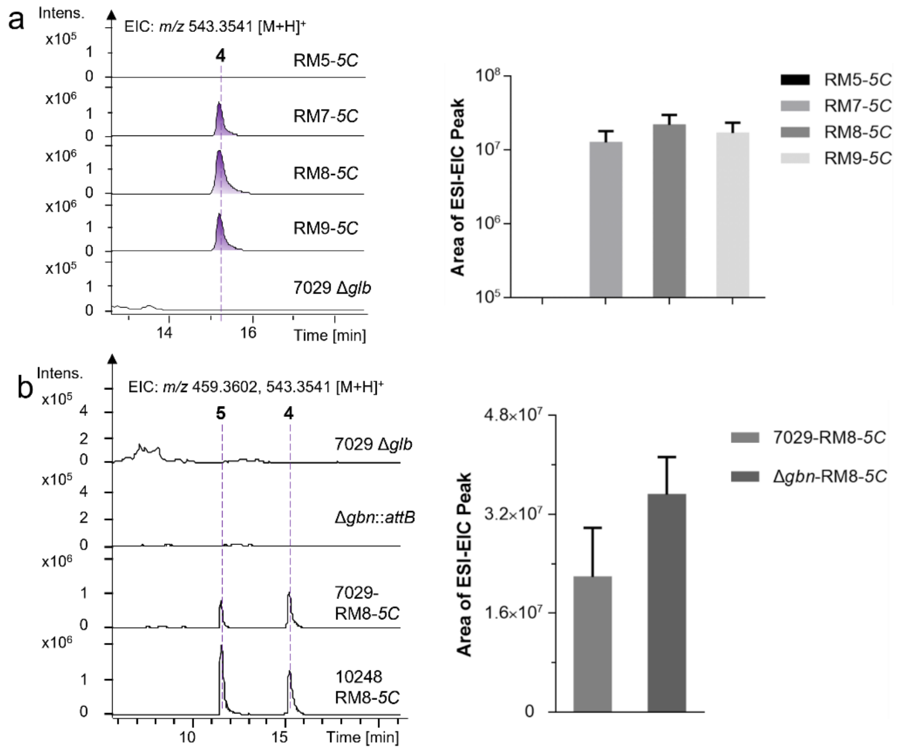

**Fig. S6.** Comparison the yields of compound **5** in different initiation modules for replacement or two heterologous hosts. a. HR-ESI-MS analysis and yield comparison of compound **5** with different initiation module changes from RzmA. a. HR-ESI-MS analysis and yield comparison of compound **5** in the chassis of *B. gladioli* Δ*gbn*::*attB* and DSM 7029Δ*glb*, EIC: *m/z* 459.3602 and 543.3541 [M+H] + .

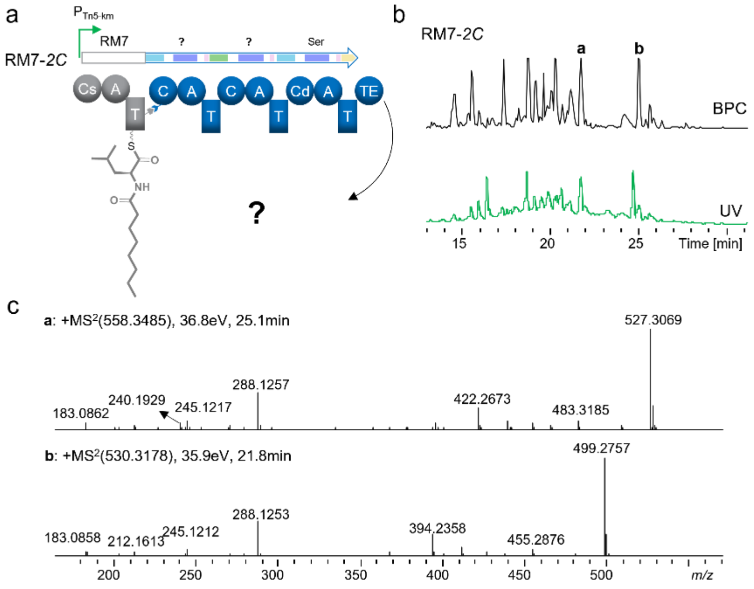

**Fig. S7.** Engineering the initiation unit to access the BGC of *2C* in *M. rhizoxinica* HKI 454. a. Schematic representation of product synthesis after modification of *2C* gene cluster by the RzmA initiation module RM7. b. HRESIMS analysis of crude extracts from the recombinant strain HKI 454/RM7-*2C*. (c) MS/MS fragmentation analysis of compound **a** and **b**.

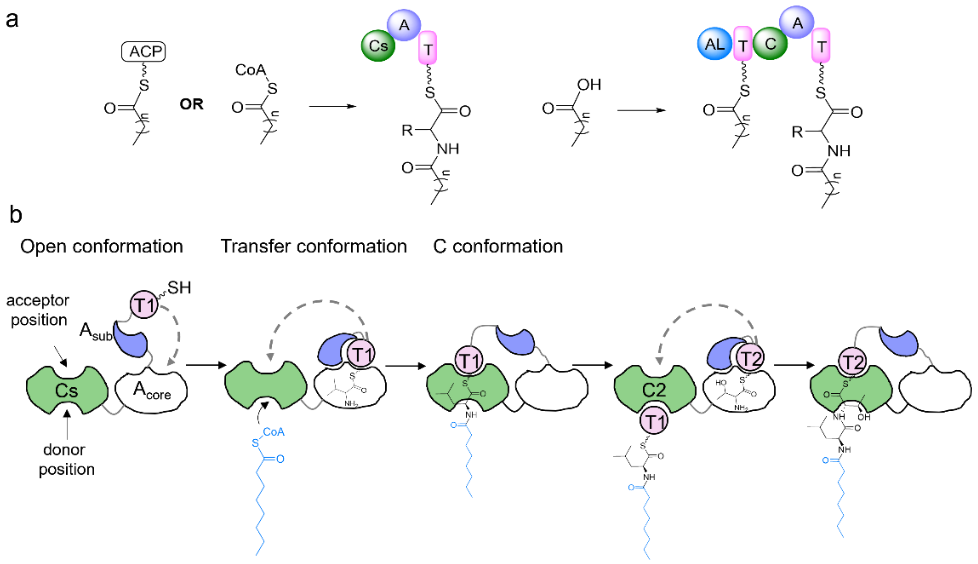

**Fig. S8.** Loading of lipid chains in nonribosomal lipopeptide and dynamic interaction between NRPS initiation regions. The figure is draw as an example of the replacement of RzmA-CsAT by HolA-CsAT (HM7) (20).

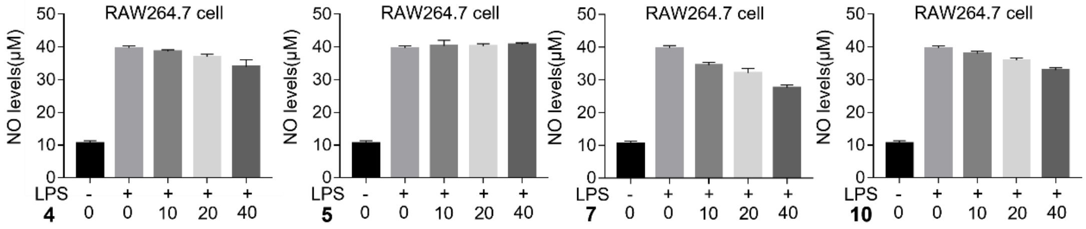

**Fig. S9.** Anti-inflammatory activity assay for compounds **4**, **5**, **7** and **10**. The *t* test unpaired (Mean with SD), LPS (10μg/mL), compounds (μM).

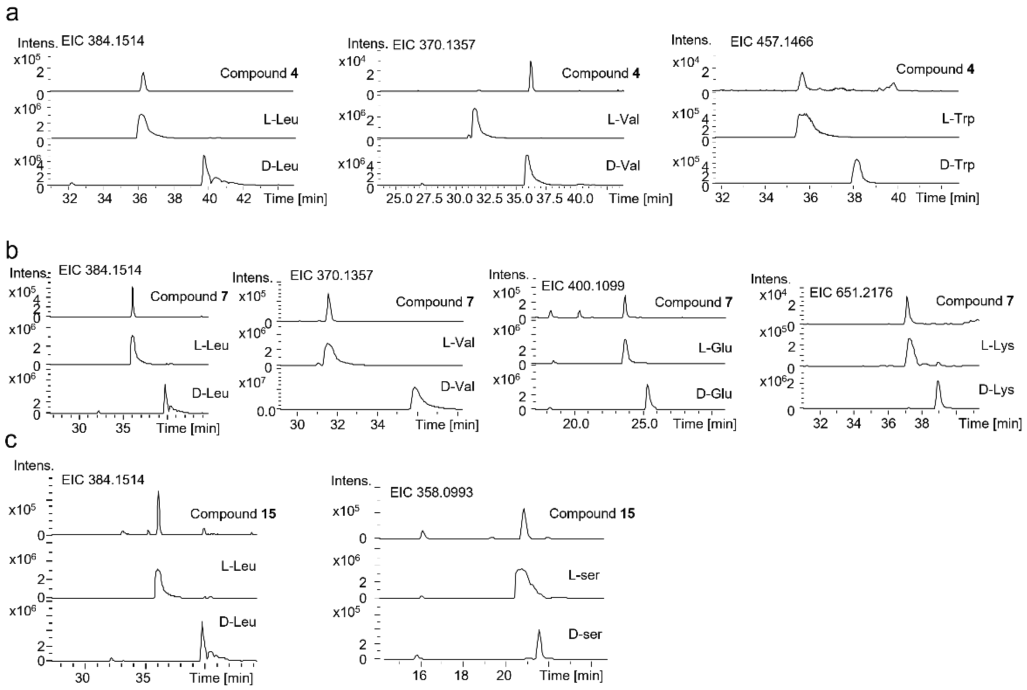

**Fig. S10.** Marfey's analysis of the amino acid constituents of compounds **4** and **7**.

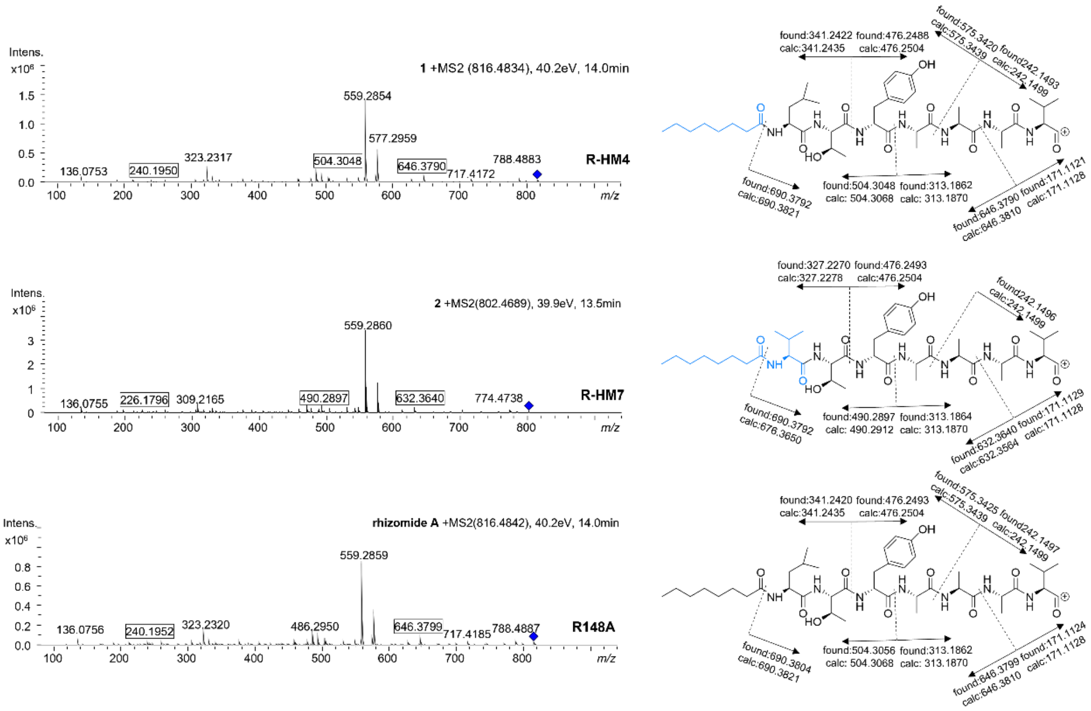

**Fig. S11.** HR-ESI-MS analysis of MS/MS fragments of compound **1** and **2** in GB05-MtaA.

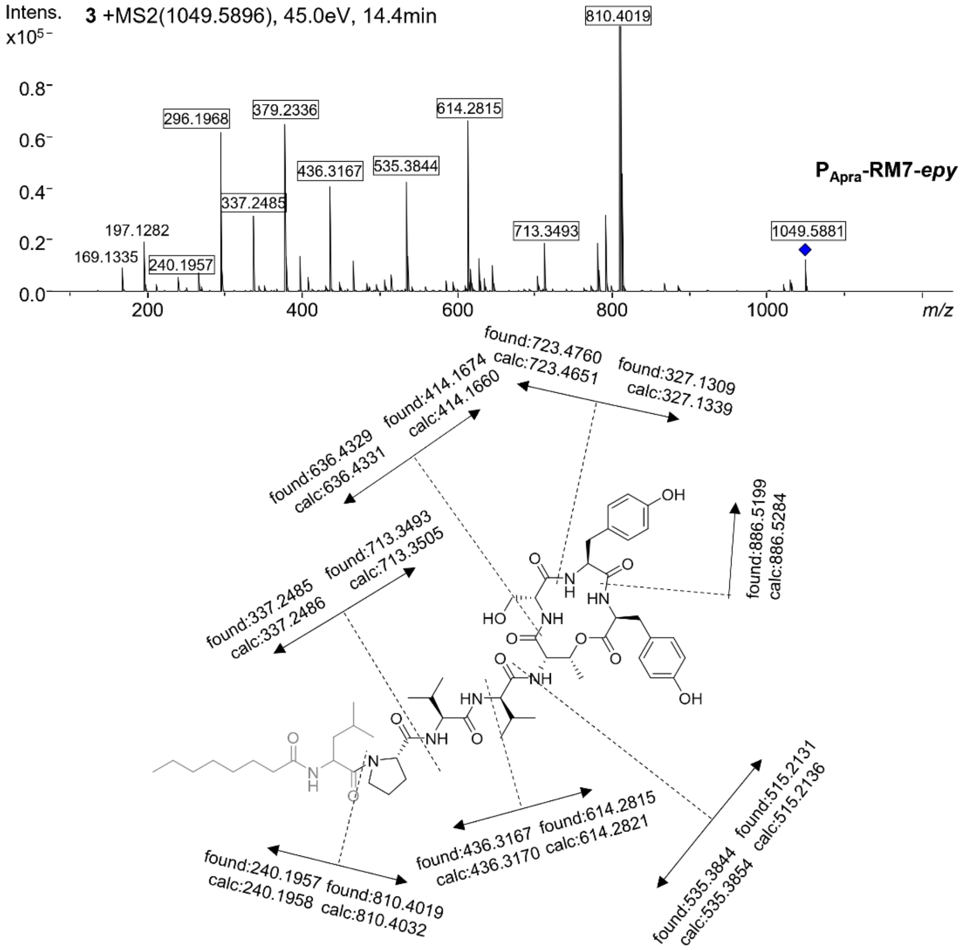

**Fig. S12.** HR-ESI-MS analysis of MS/MS fragments of compound **3** in 7029 Δ*glb*.

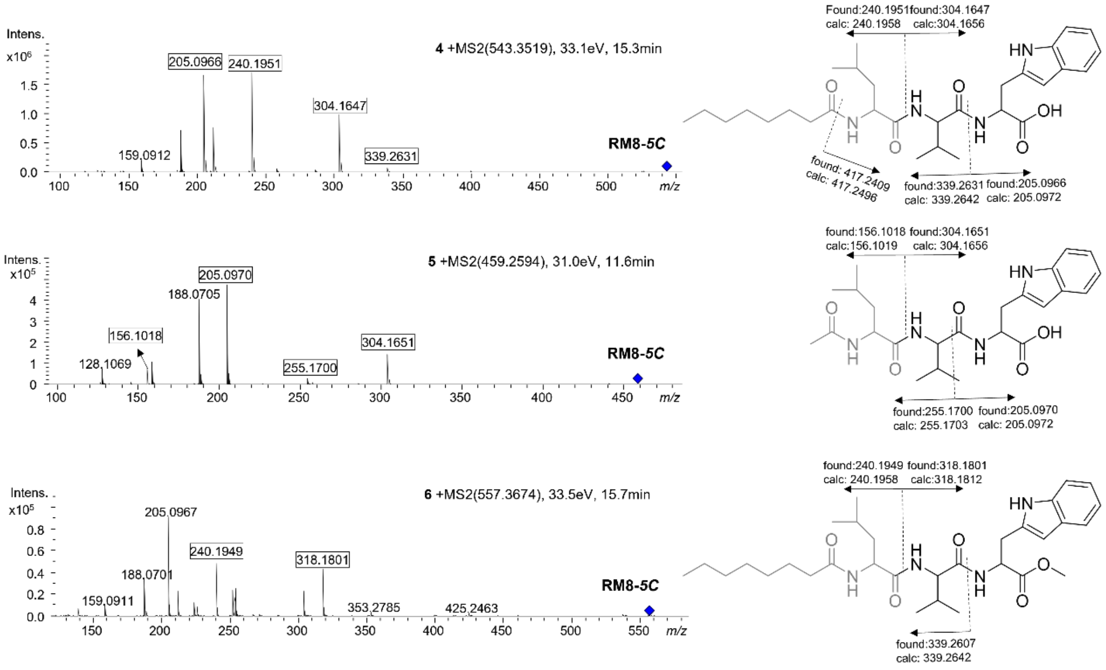

**Fig. S13**. HR-ESI-MS analysis of MS/MS fragments of compound **4**, **5** and **6** in 7029 Δ*glb*.

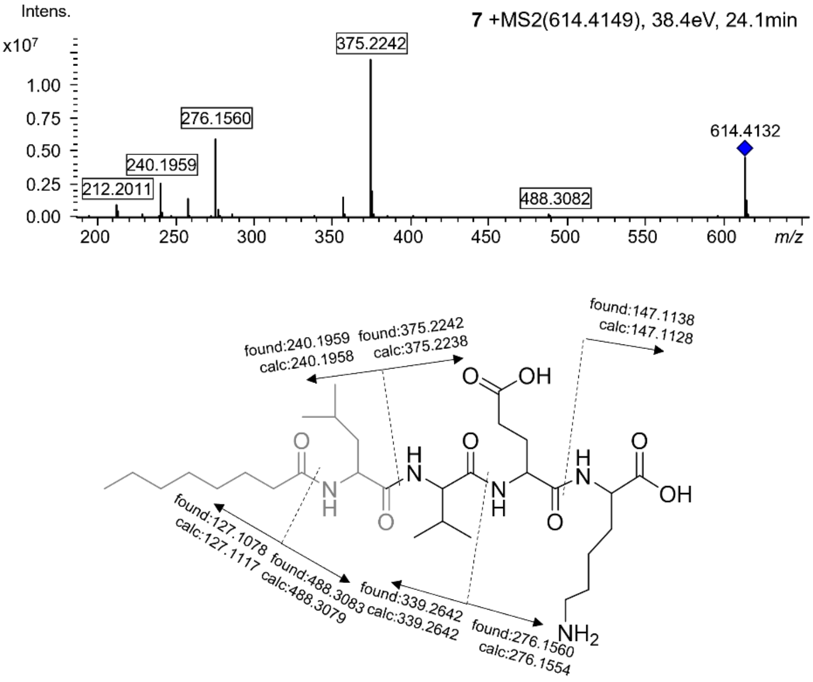

**Fig. S14.** HR-ESI-MS analysis of MS/MS fragments of compound **7** in HKI 454 Δ*rhi*.

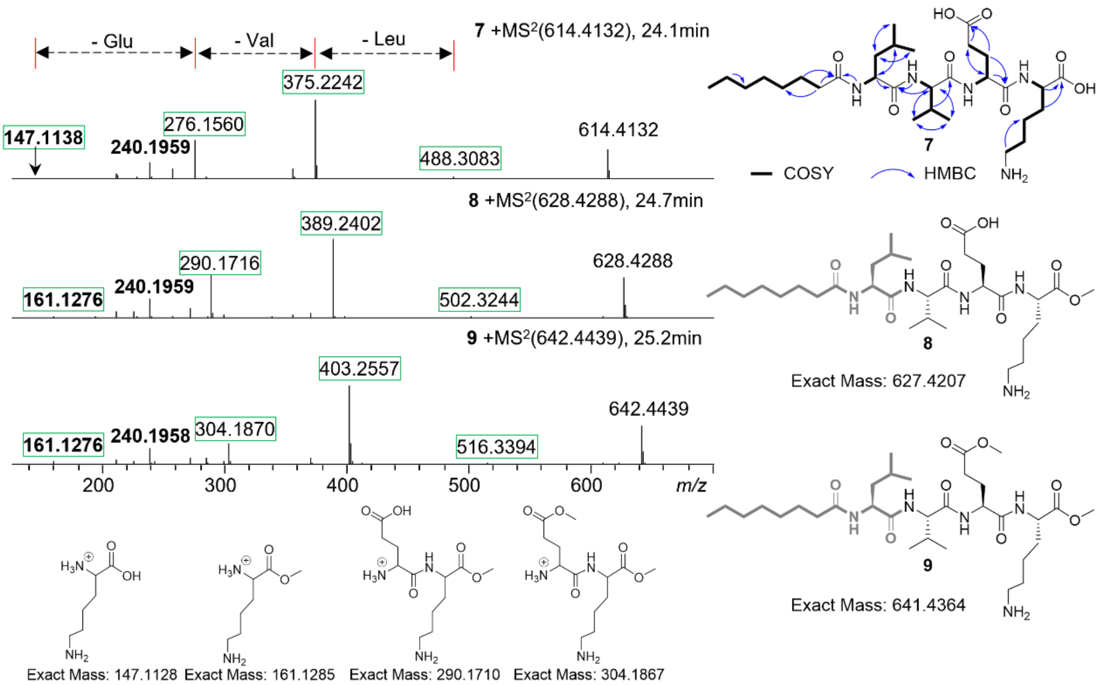

**Fig. S15.** Comparative analysis of HR-ESI-MS analysis of MS/MS fragments of compound **8**, **9** and **7** in HKI 454 Δ*rhi*.

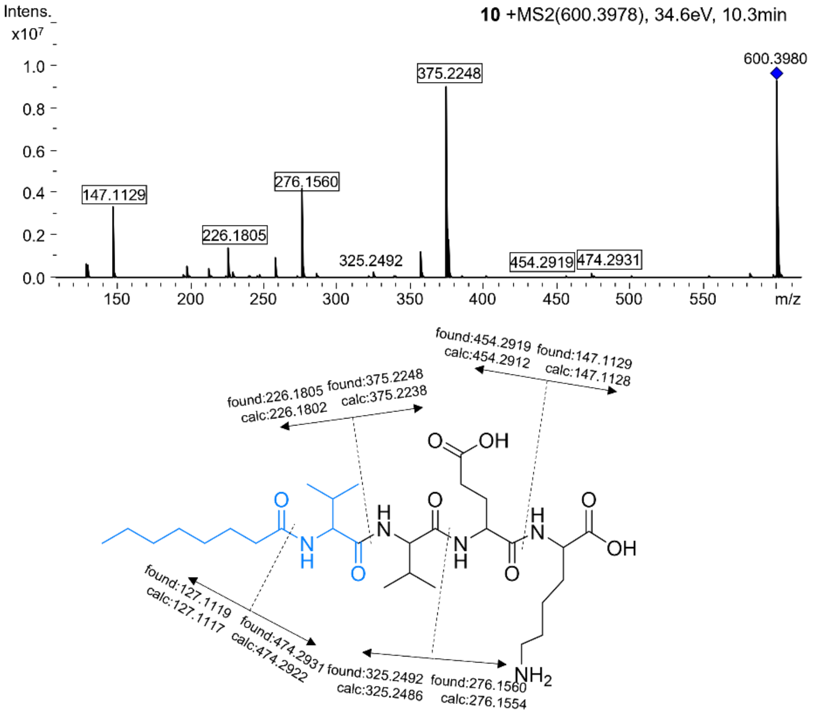

**Fig. S16.** HR-ESI-MS analysis of MS/MS fragments of compound **10** in 10248 Δ*gbn*::*attB*.

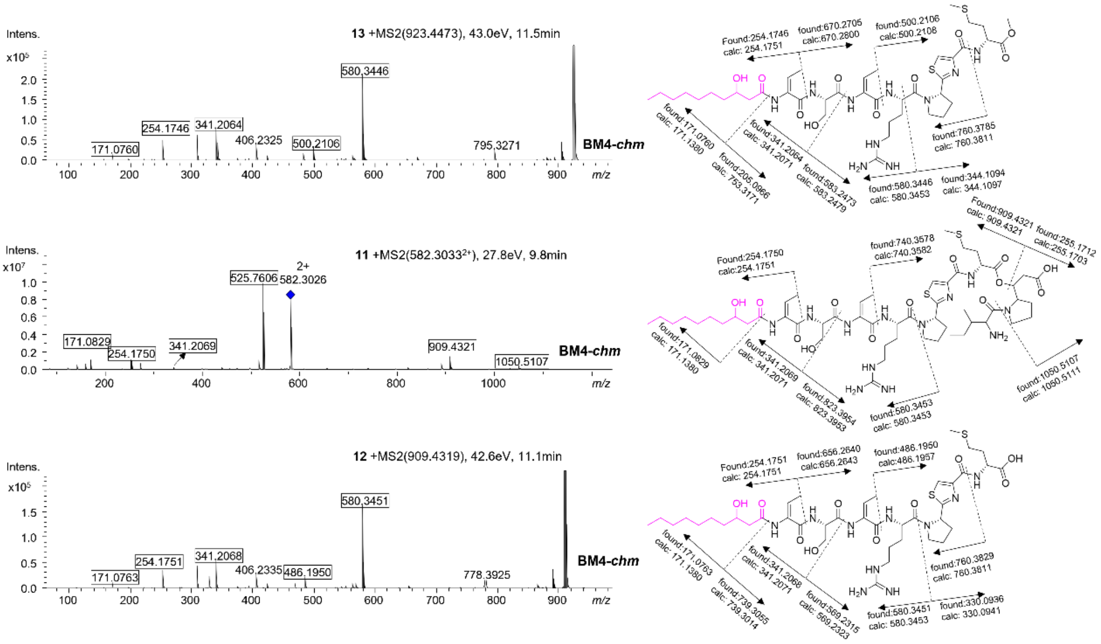

**Fig. S17.** HR-ESI-MS analysis of MS/MS fragments of compound **11**, **12** and **13**.

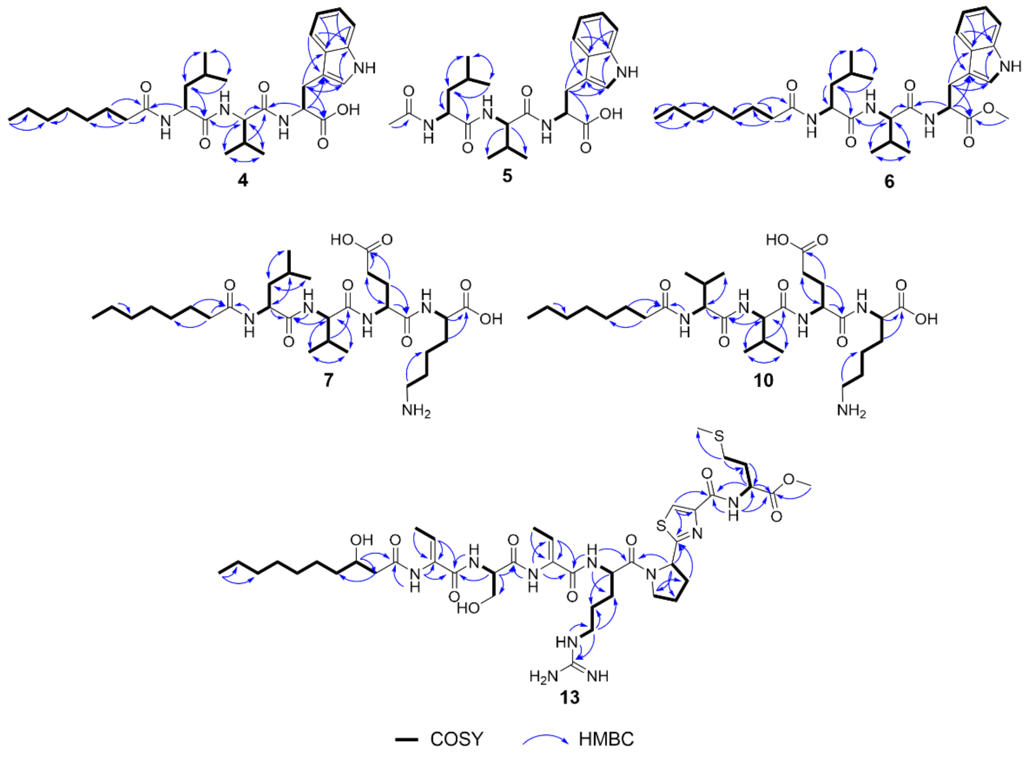

**Fig. S18.** Complete structures and Key COSY and HMBC correlations of compounds **4**, **5**, **6**, **7**,**10** and **13**.

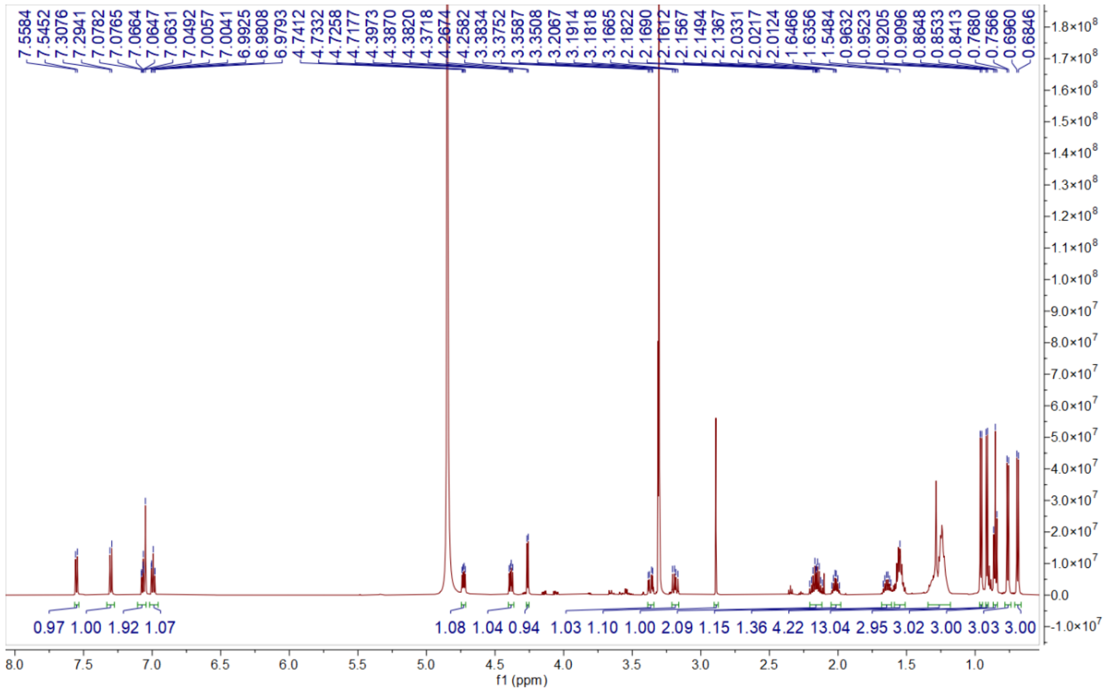

**Fig. S19.** $^{1}H$ NMR spectrum of compound **4** in $CH_3DO$.

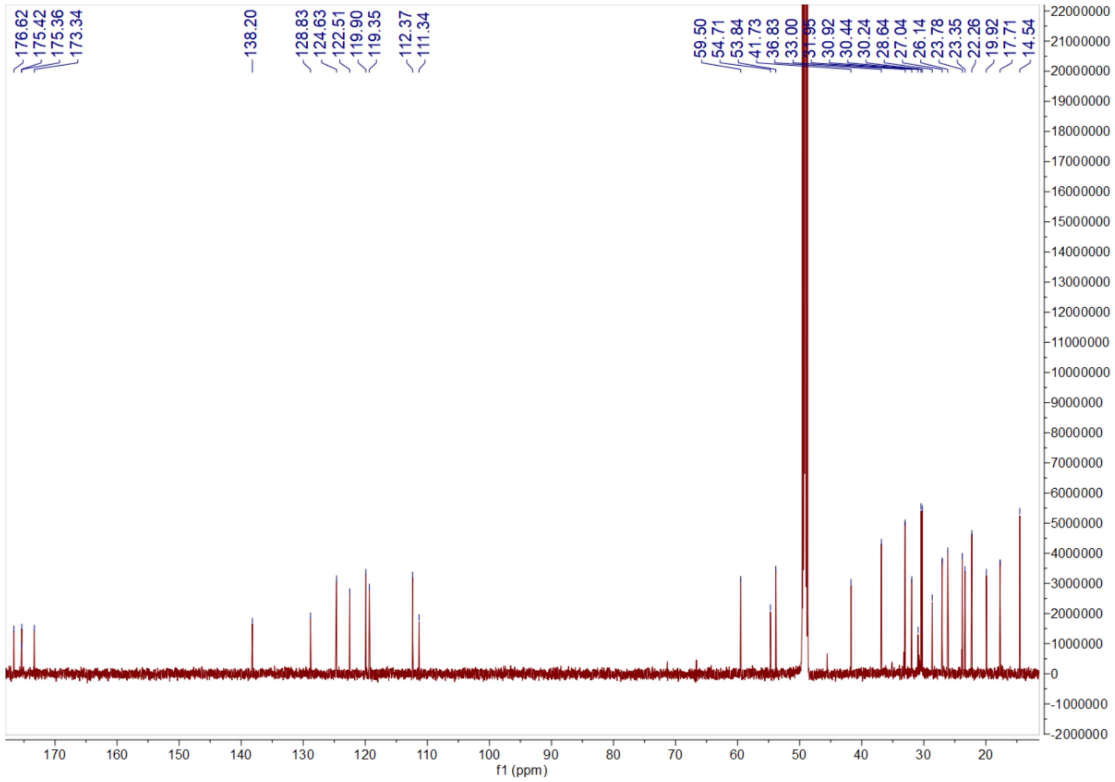

**Fig. S20.** $^{13}C$ NMR spectrum of compound **4** in $CH_3DO$.

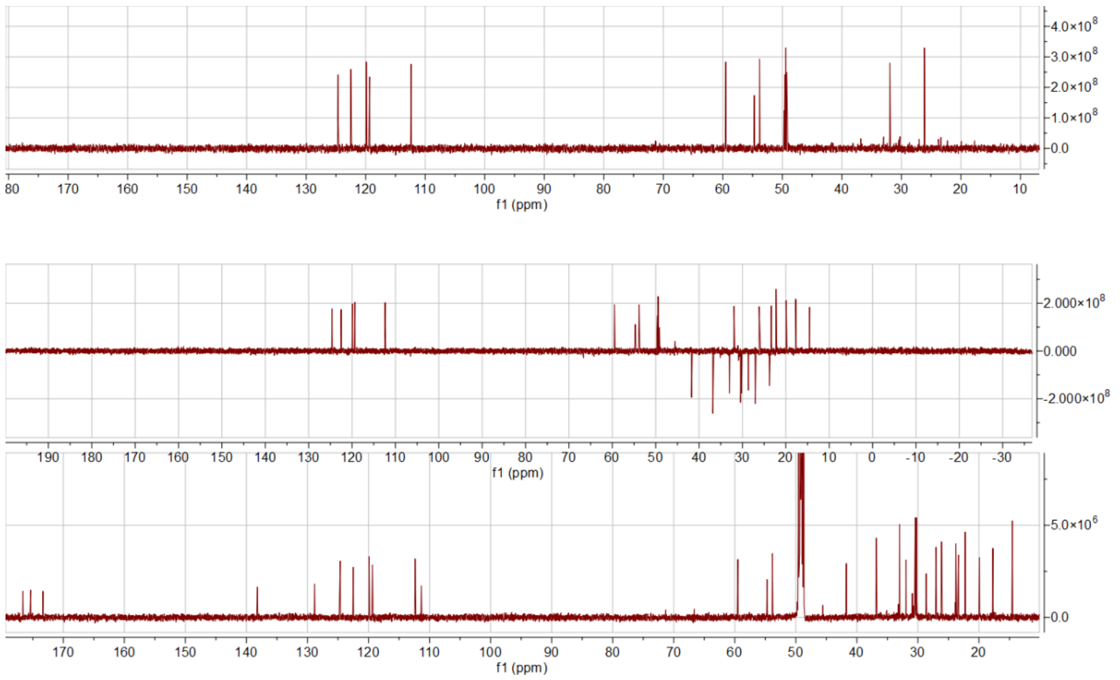

**Fig. S21.** DEPT spectrum of compound **4** in $CH_3DO$.

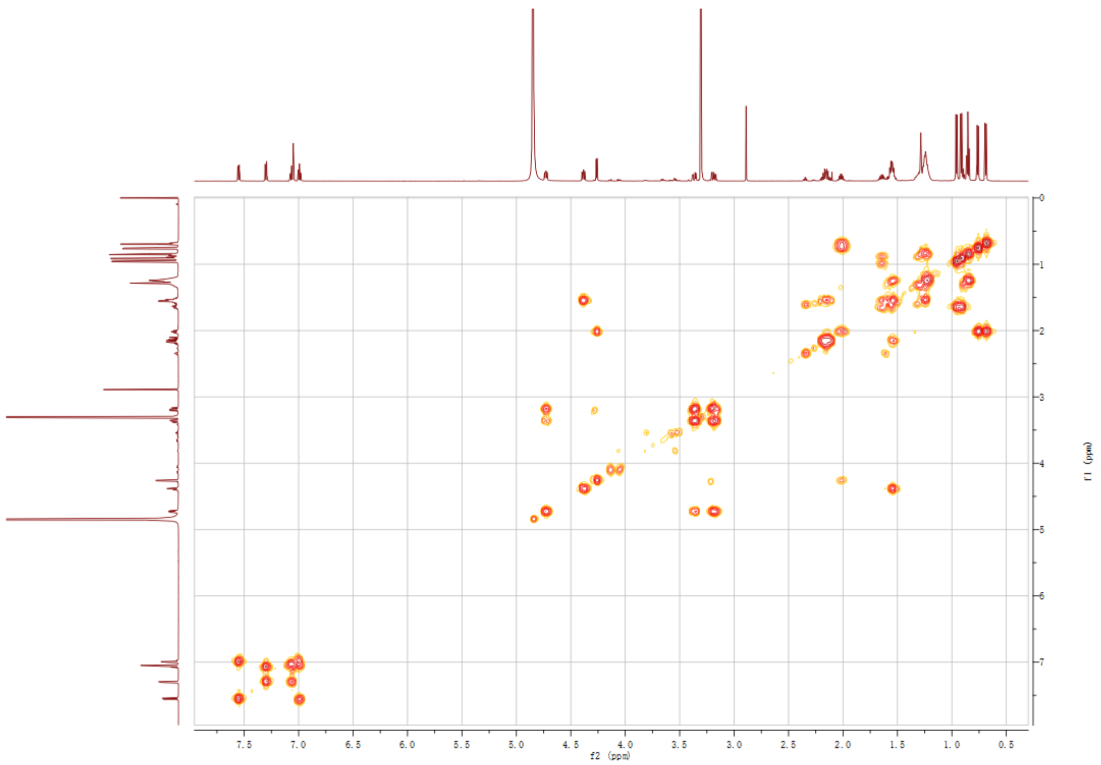

**Fig. S22.** $^1H$-$^1H$ COSY NMR spectrum of compound **4** in $CH_3DO$.

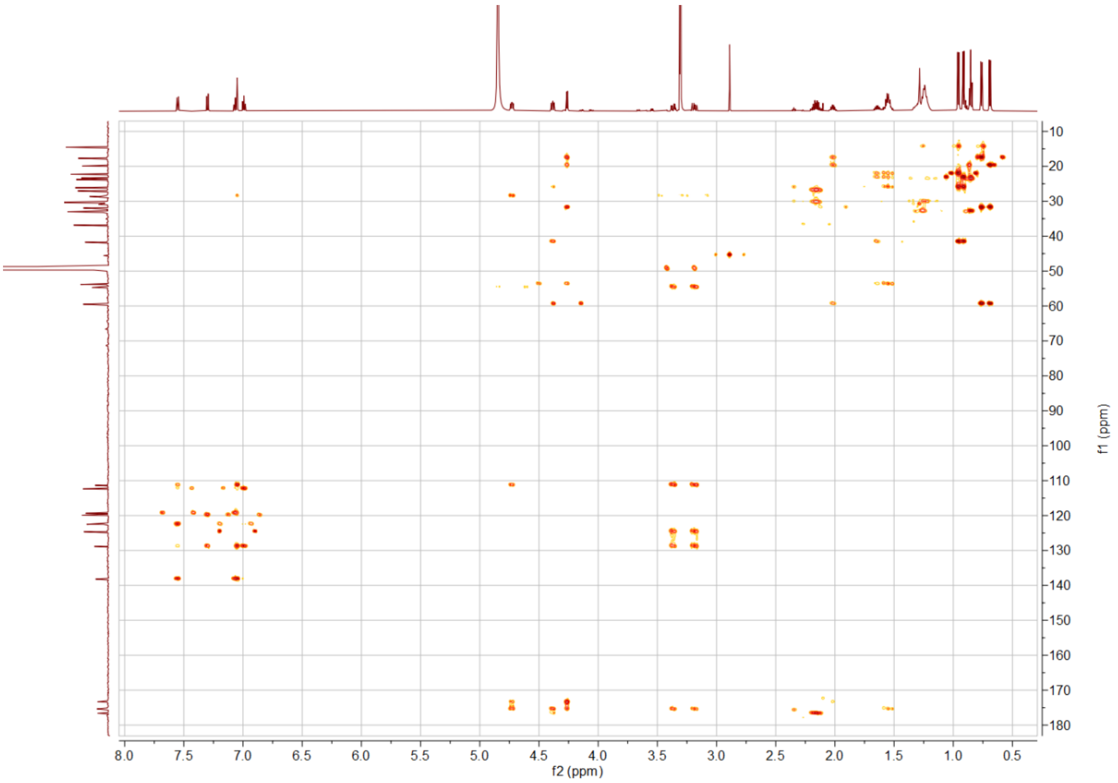

**Fig. S23.** HMBC spectrum of compound **4** in CH3DO.

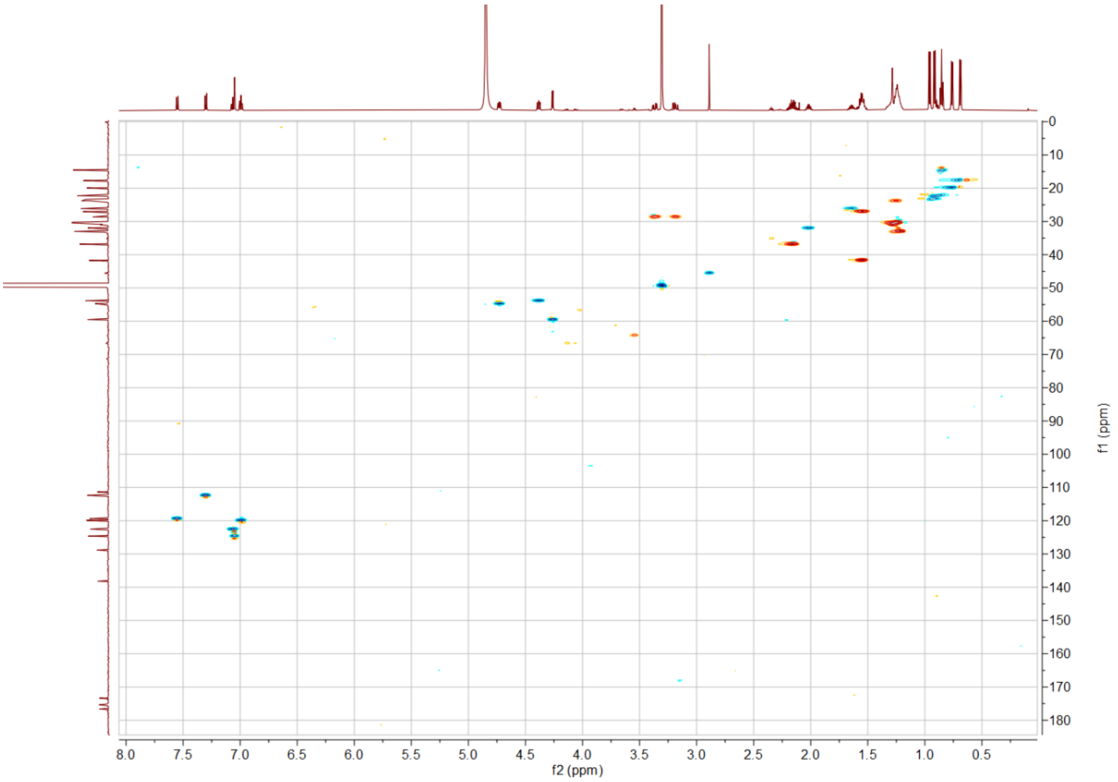

**Fig. S24.** HSQC spectrum of compound **4** in CH3DO.

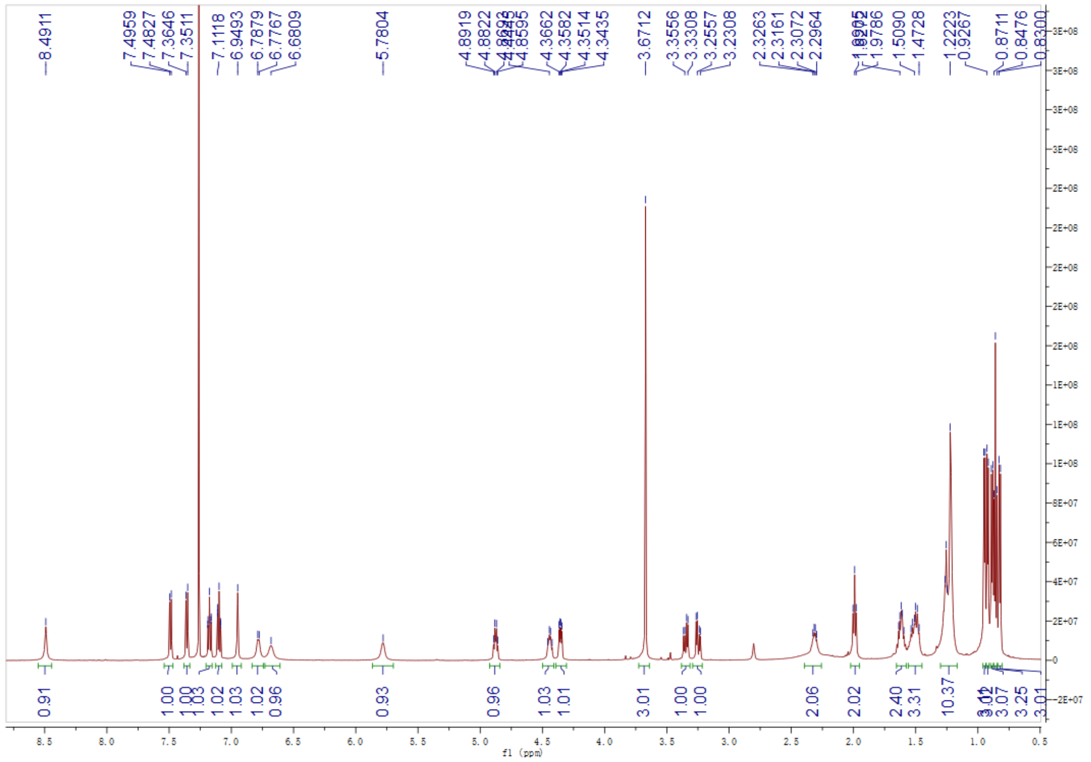

**Fig. S25.** $^{1}H$ NMR spectrum of compound **5** in $CDCl_3$.

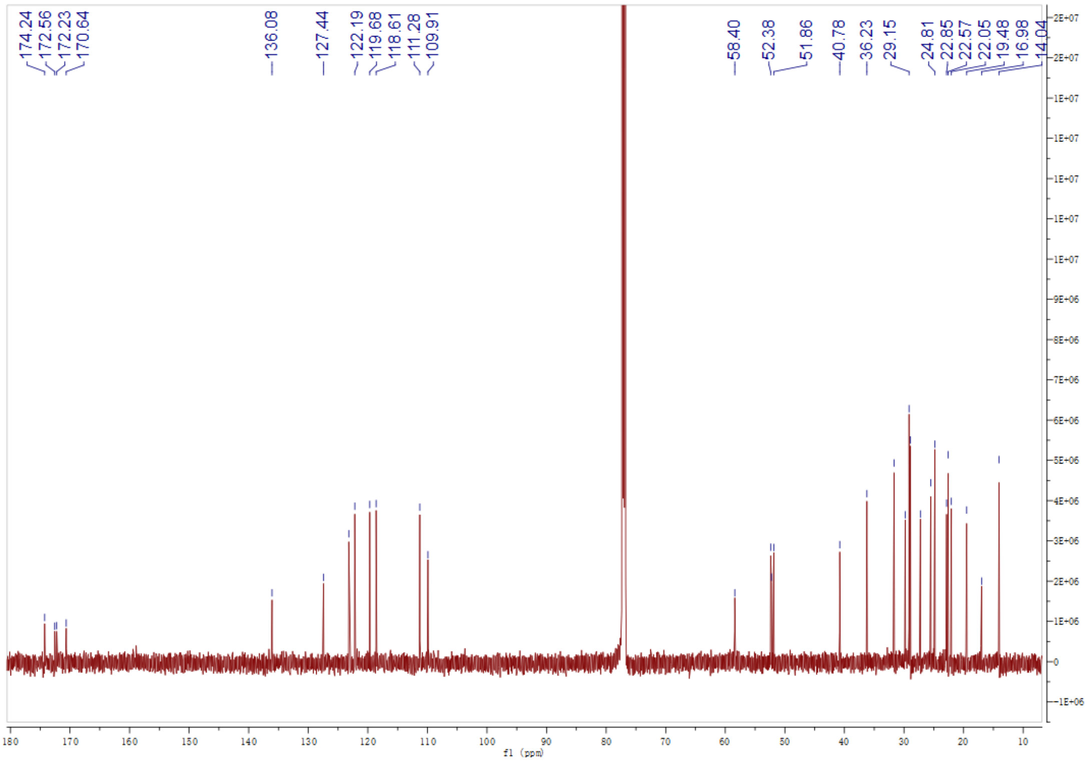

**Fig. S26.** $^{13}C$ NMR spectrum of compound **5** in $CDCl_3$.

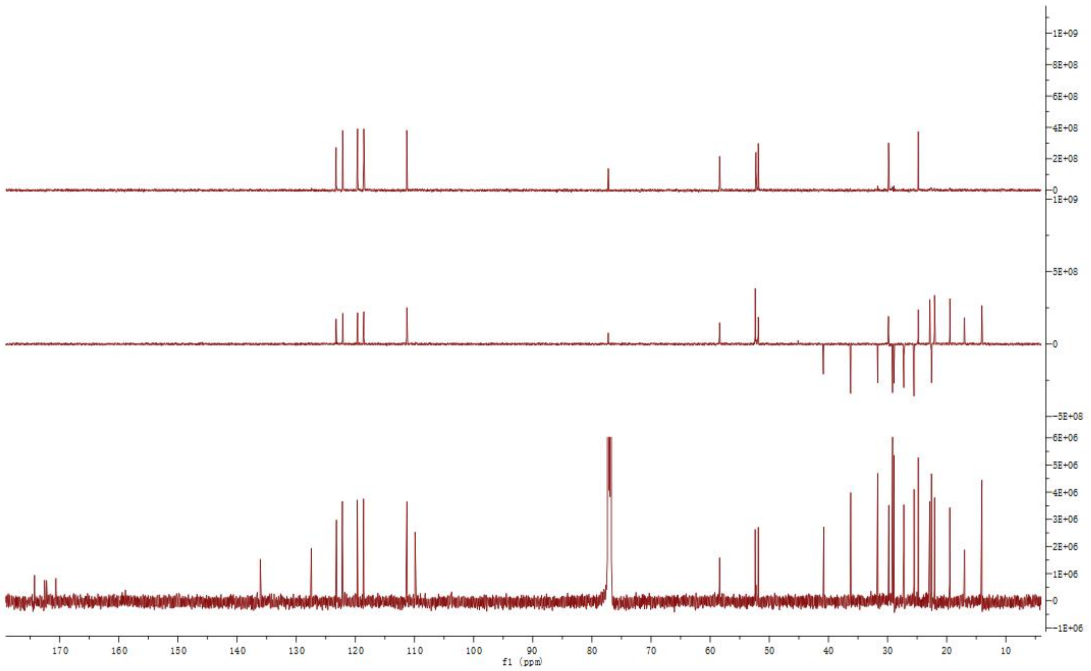

**Fig. S27.** DEPT spectrum of compound **5** in $CDCl_3$.

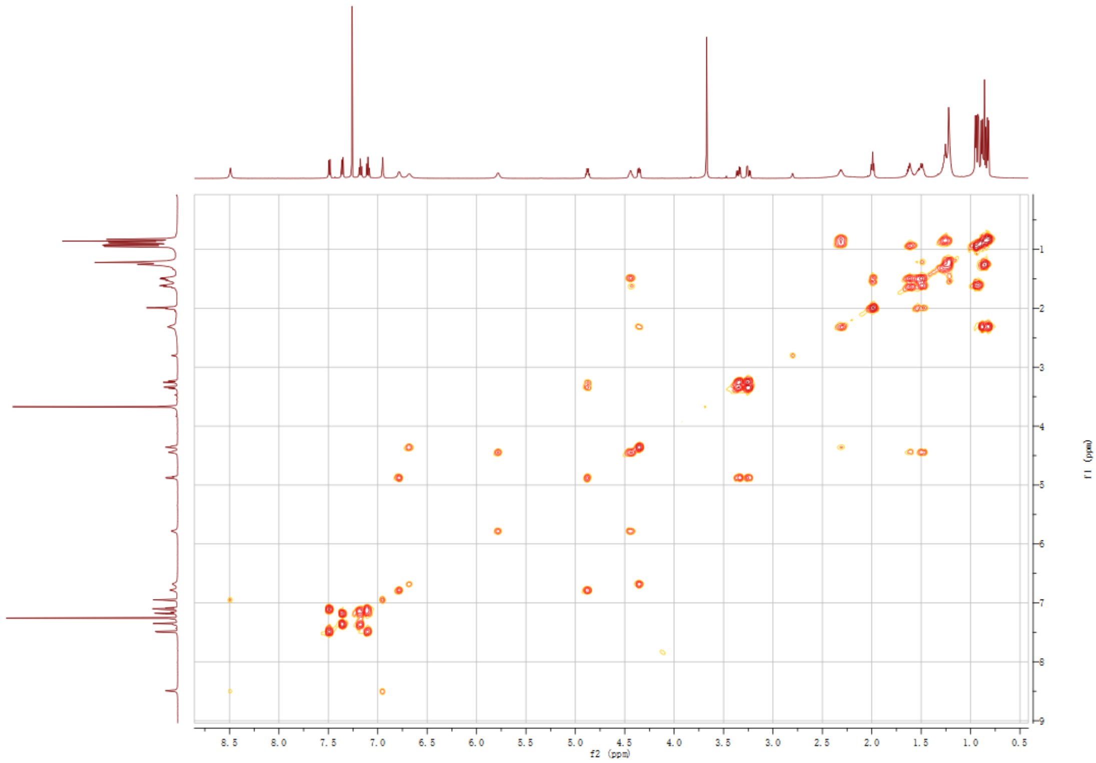

**Fig. S28.** $^1H$-$^1H$ COSY NMR spectrum of compound **5** in $CDCl_3$.

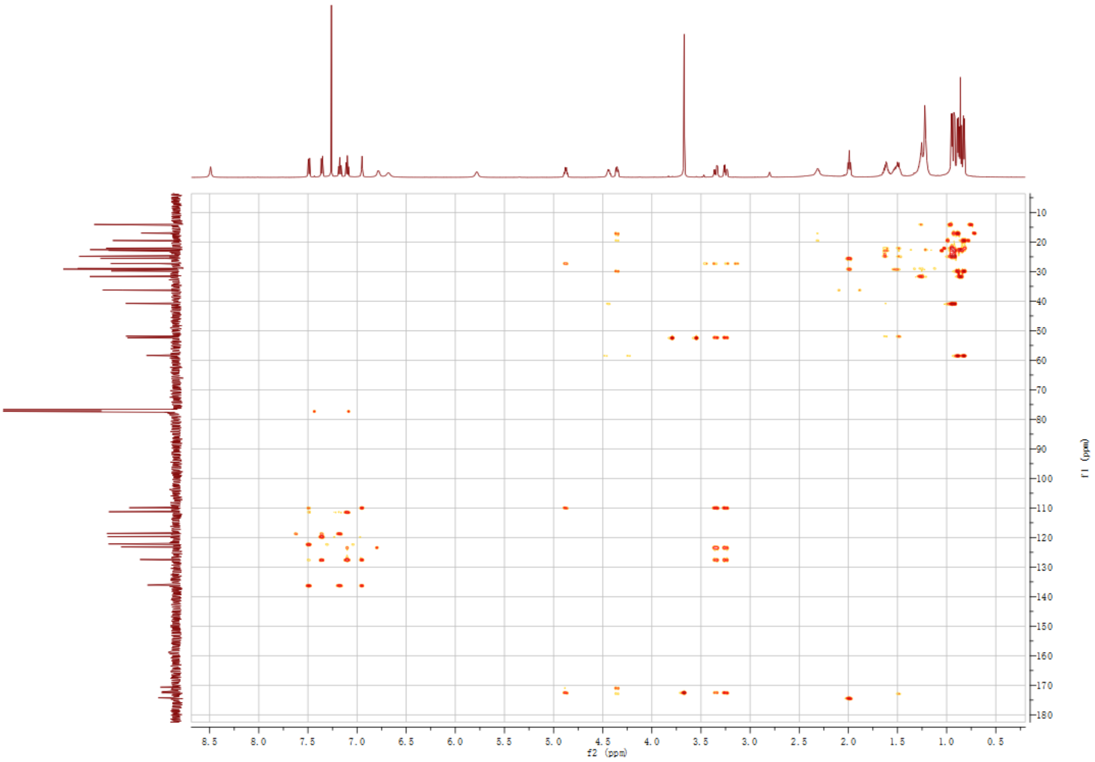

**Fig. S29.** HMBC spectrum of compound **5** in $CDCl_3$.

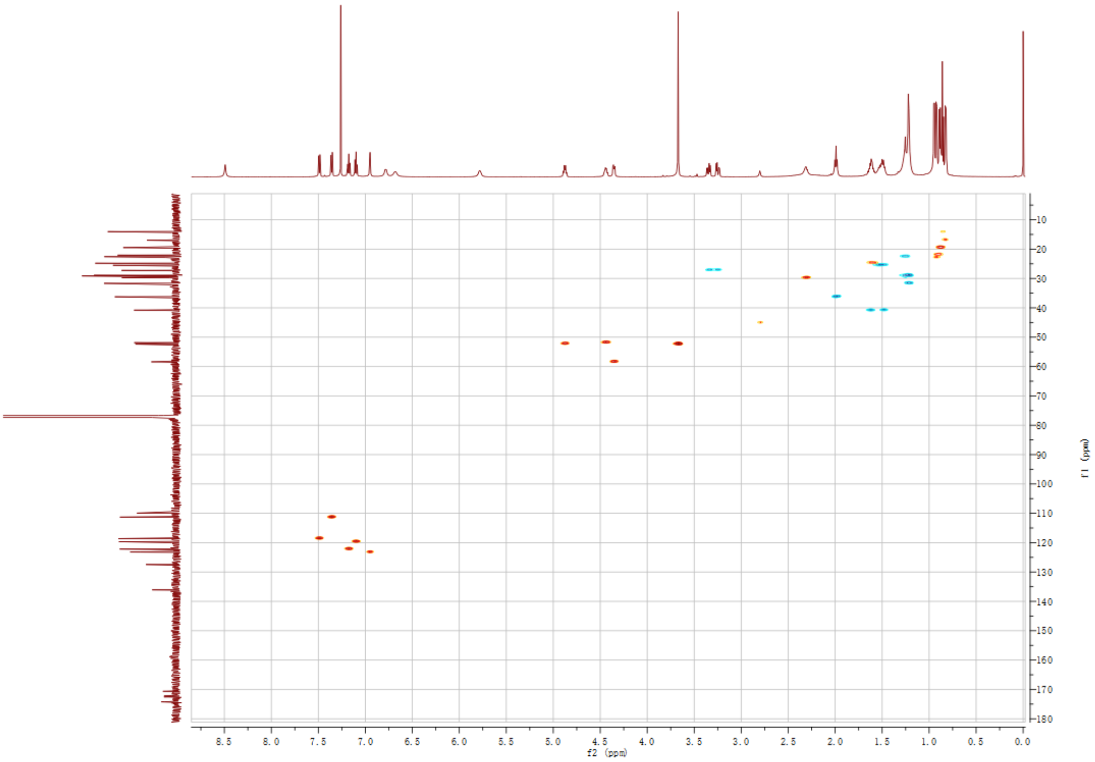

**Fig. S30.** HSQC spectrum of compound **5** in $CDCl_3$.

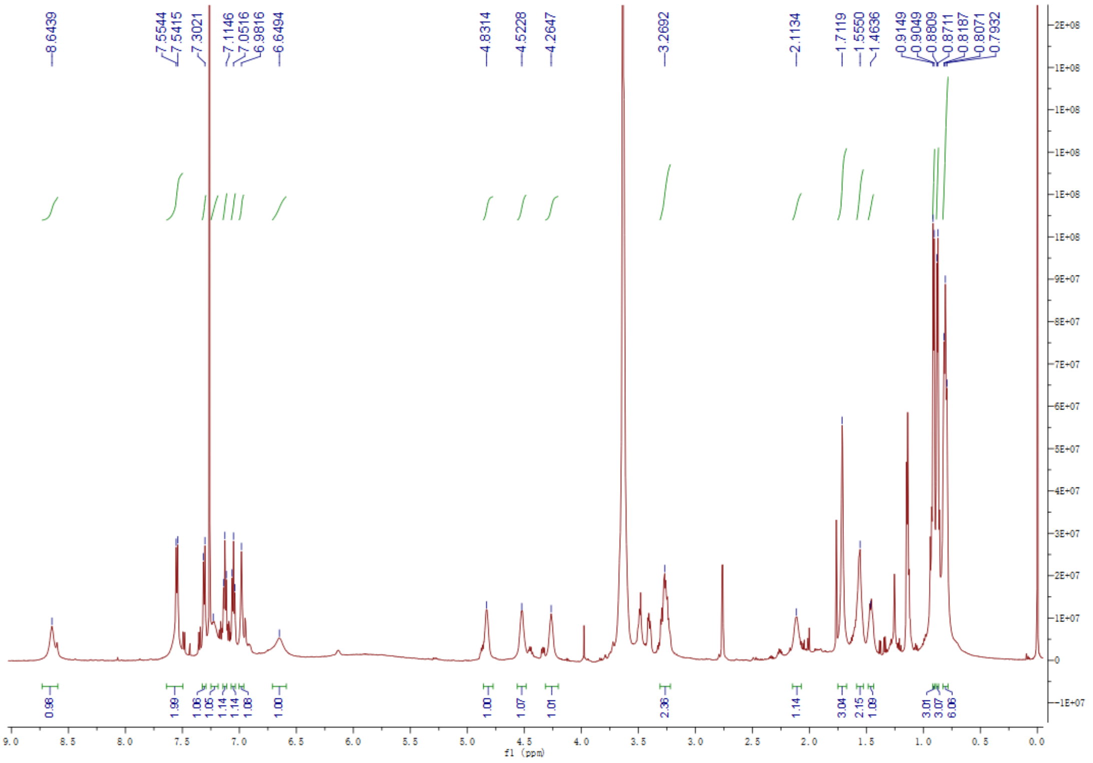

**Fig. S31.** [1]H NMR spectrum of compound **6** in $CDCl_3$.

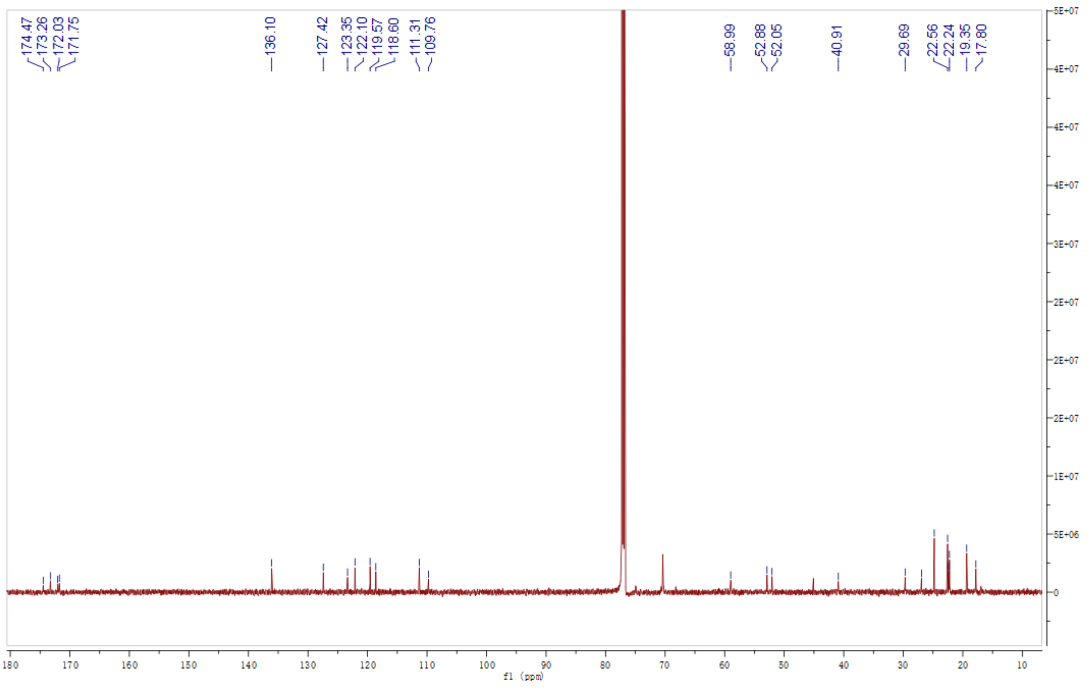

**Fig. S32.** [13]C NMR spectrum of compound **6** in $CDCl_3$.

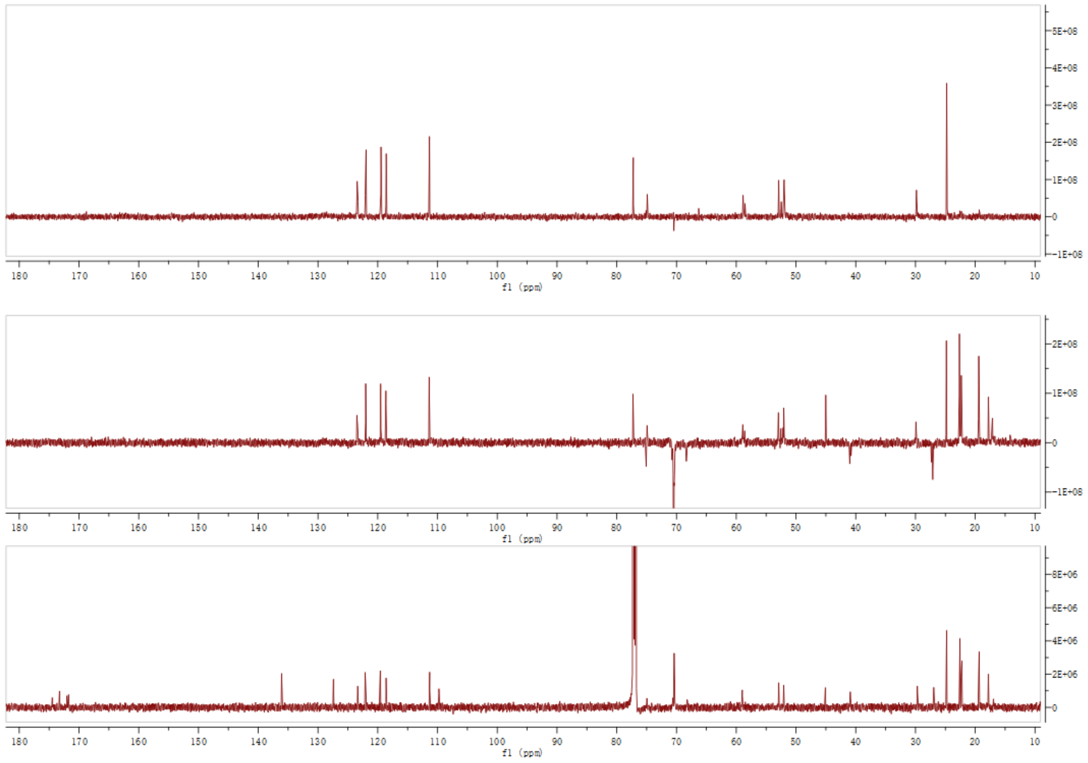

**Fig. S33.** DEPT spectrum of compound **6** in $CDCl_3$.

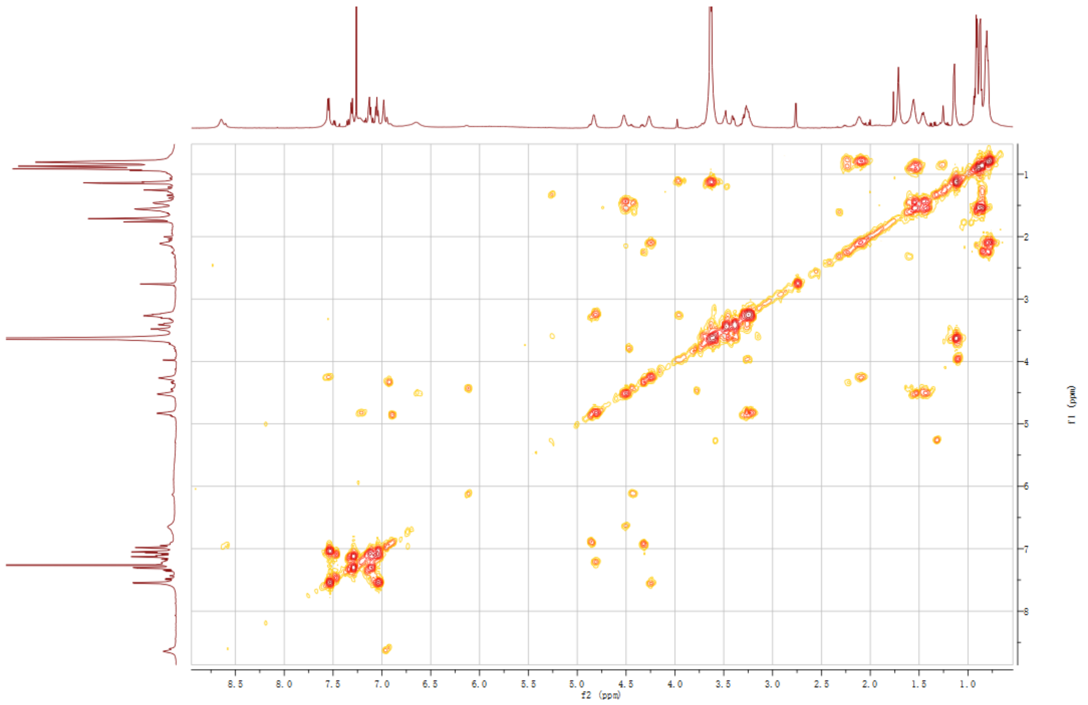

**Fig. S34.** $^{1}H$-$^{1}H$ COSY NMR spectrum of compound **6** in $CDCl_3$.

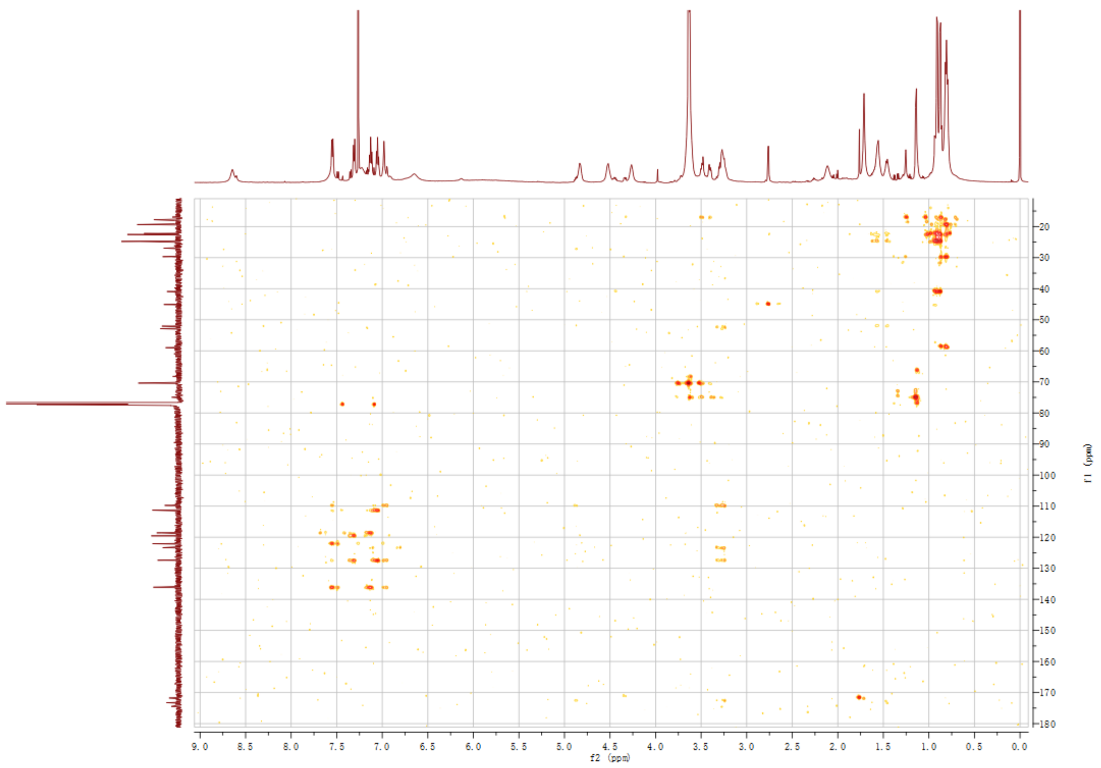

**Fig. S35.** HMBC spectrum of compound **6** in $CDCl_3$.

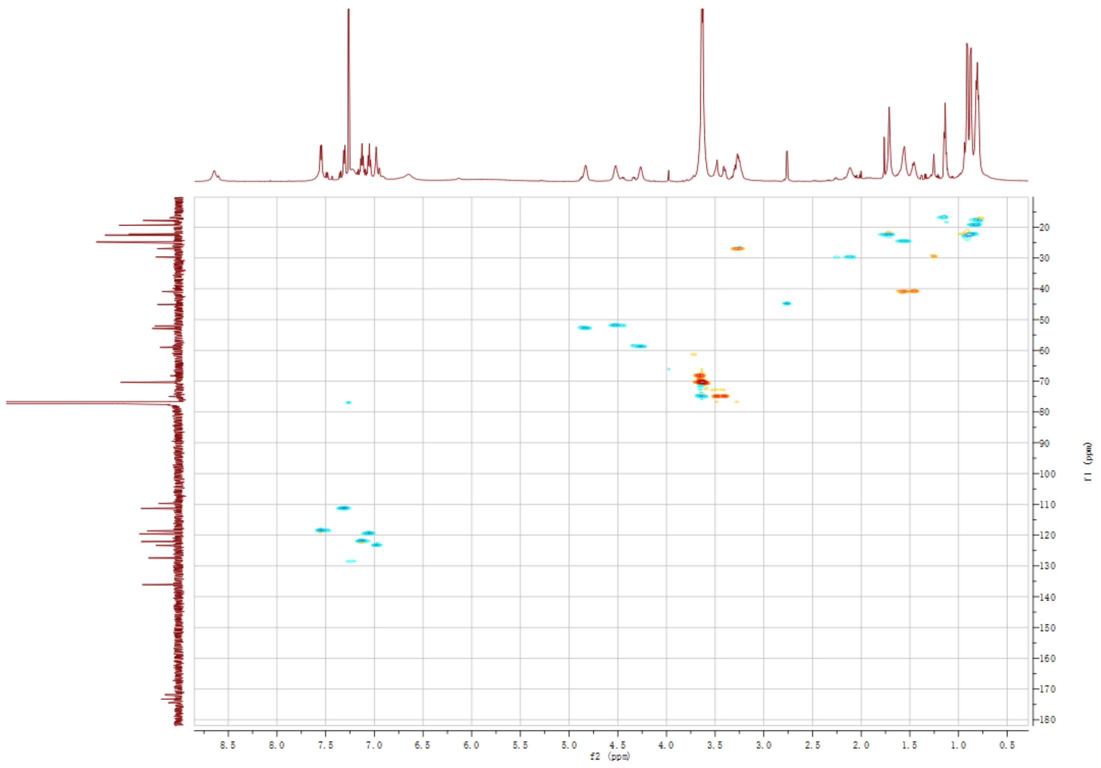

**Fig. S36.** HSQC spectrum of compound **6** in $CDCl_3$.

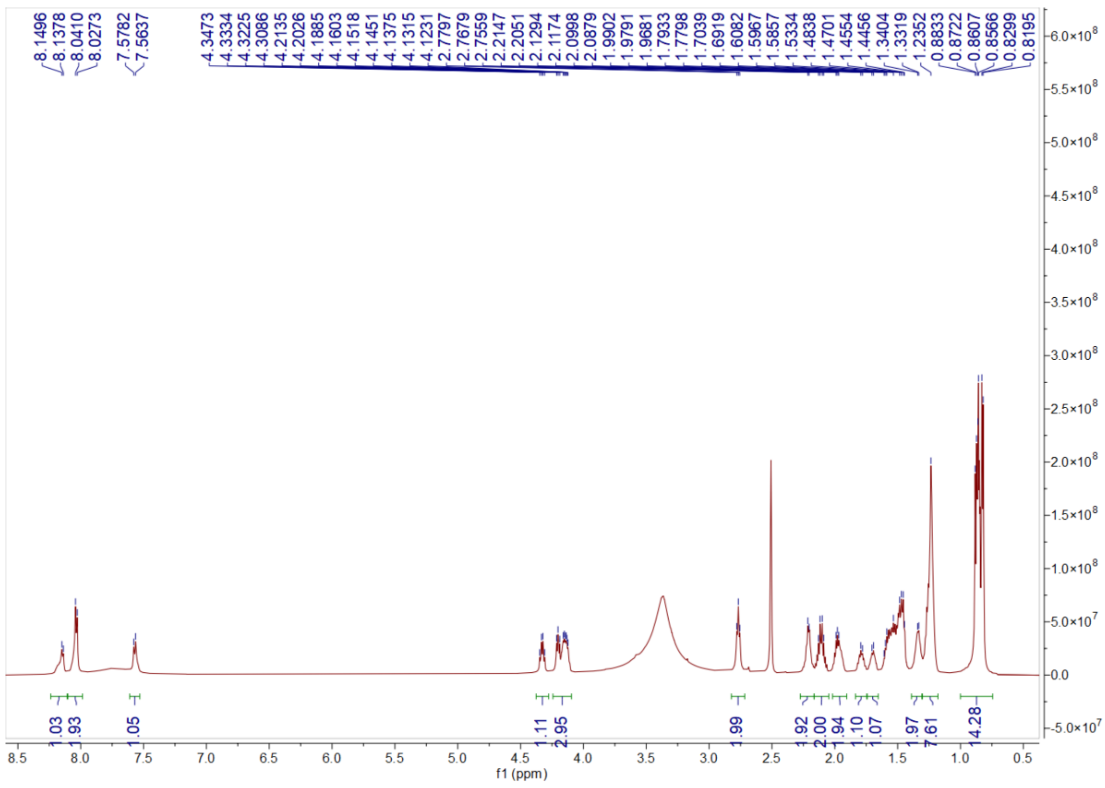

**Fig. S37.** $^{1}H$ NMR spectrum of compound **7** in DMSO-$d_6$.

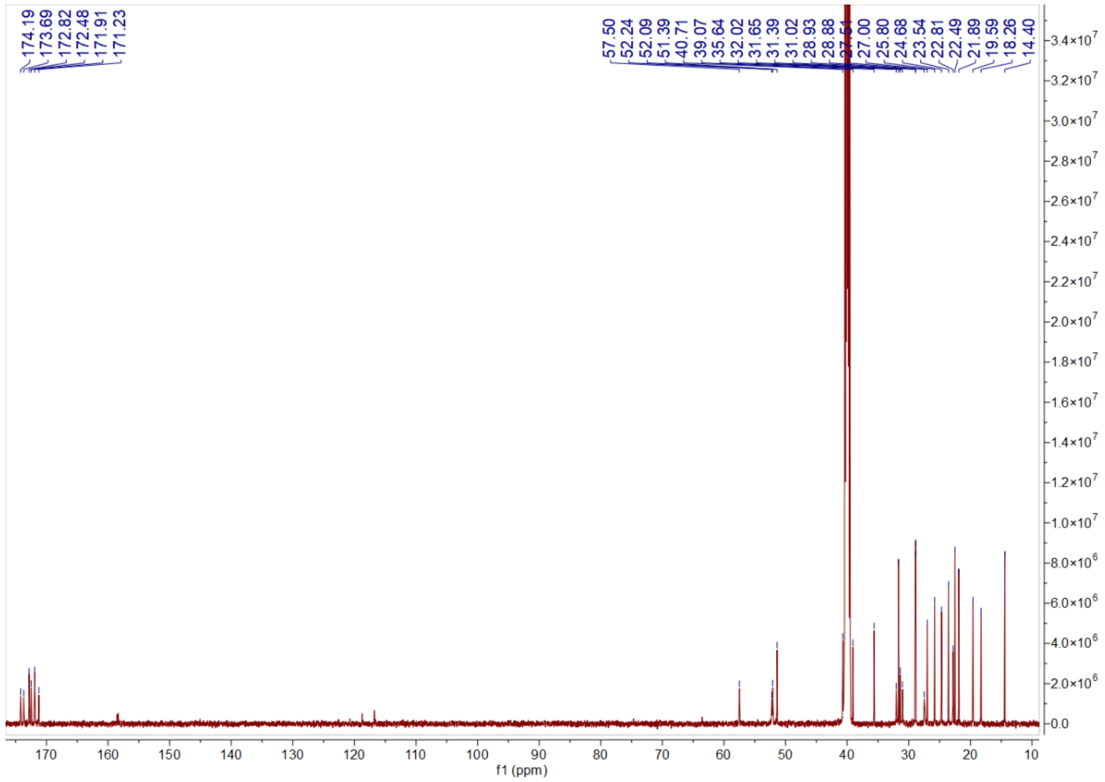

**Fig. S38.** $^{13}C$ NMR spectrum of compound **7** in DMSO-$d_6$.

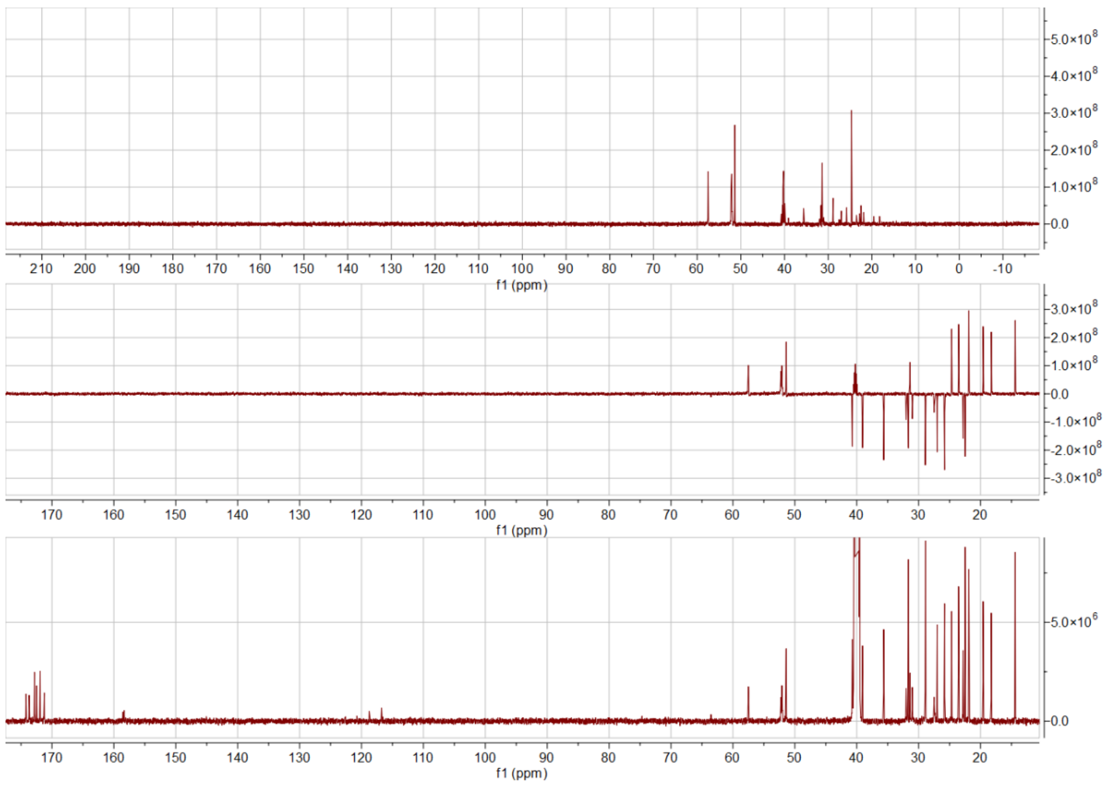

**Fig. S39.** DEPT spectrum of compound **7** in DMSO-$d_6$.

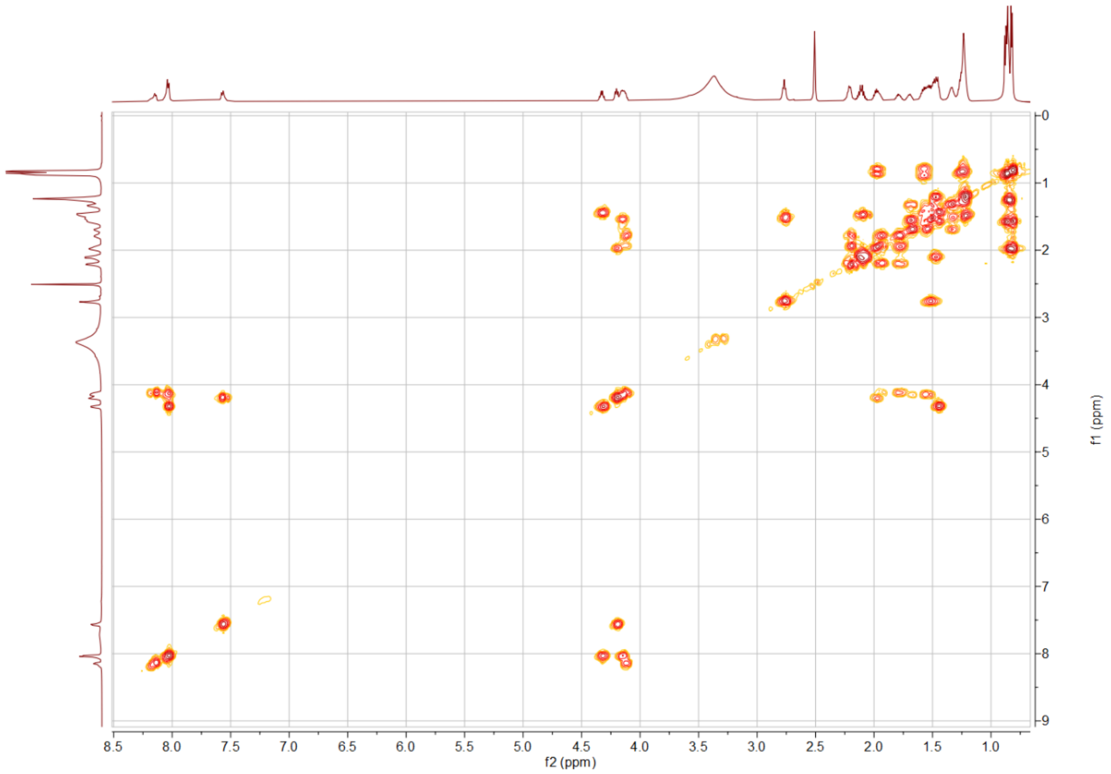

**Fig. S40.** $^1$H-$^1$H COSY NMR spectrum of compound **7** in DMSO-$d_6$.

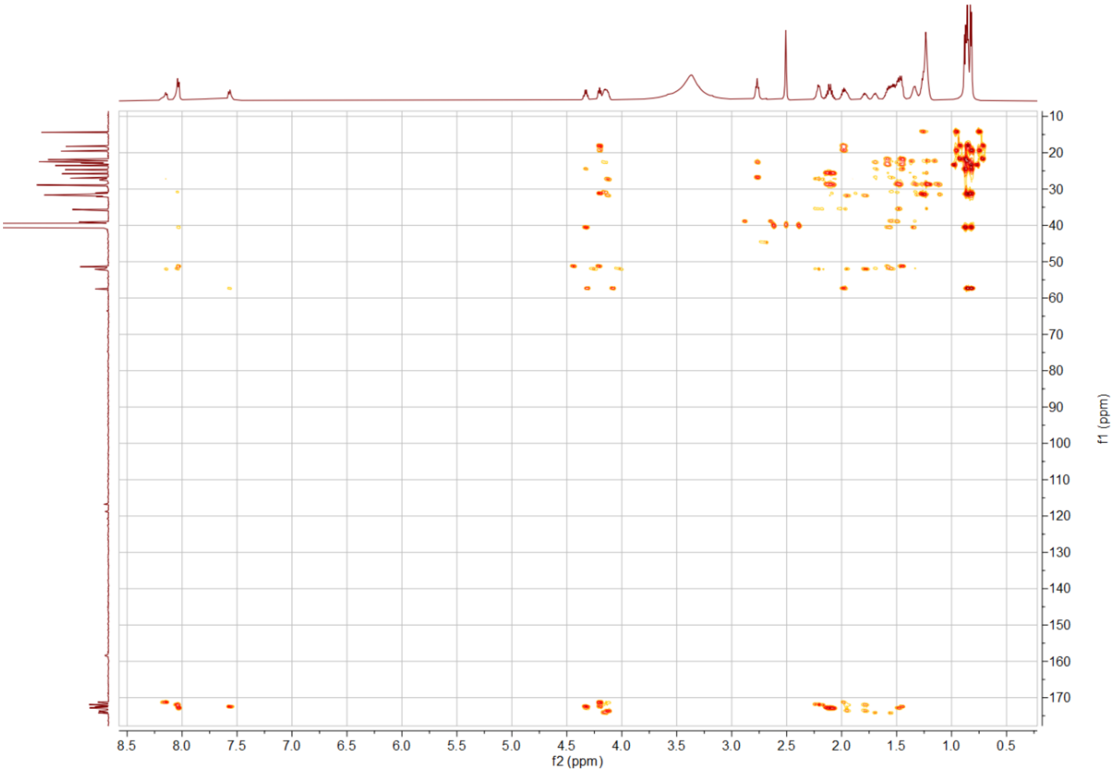

**Fig. S41.** HMBC spectrum of compound **7** in DMSO-$d_6$.

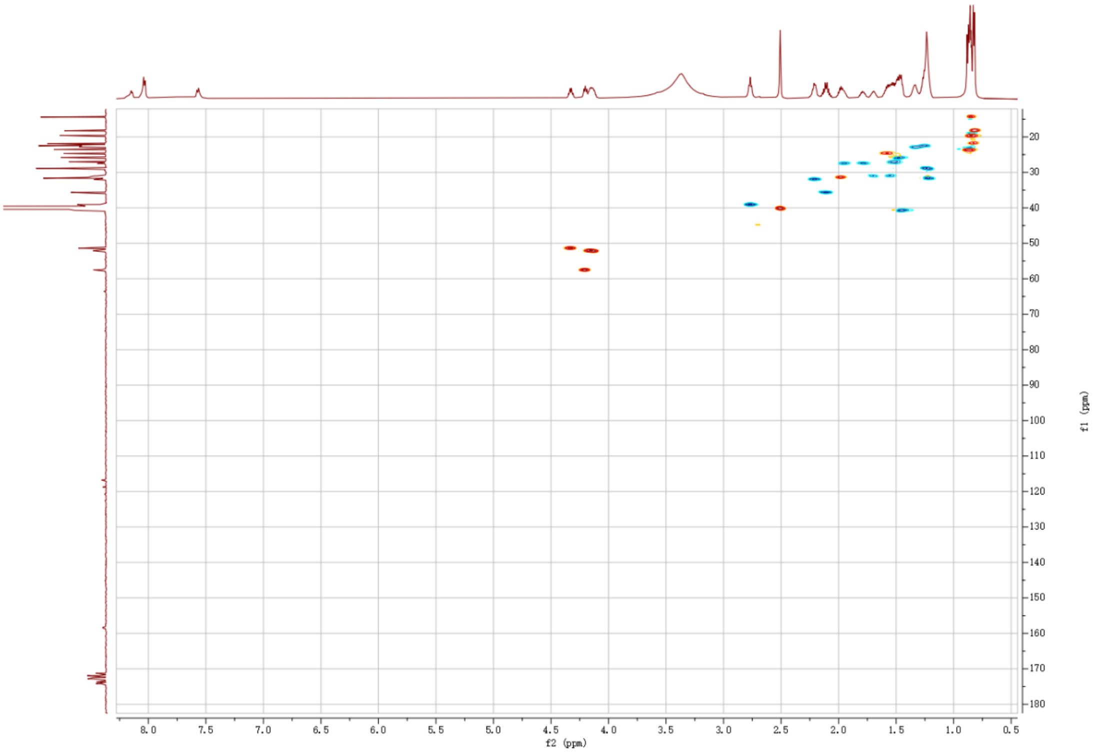

**Fig. S42.** HSQC spectrum of compound **7** in DMSO-$d_6$.

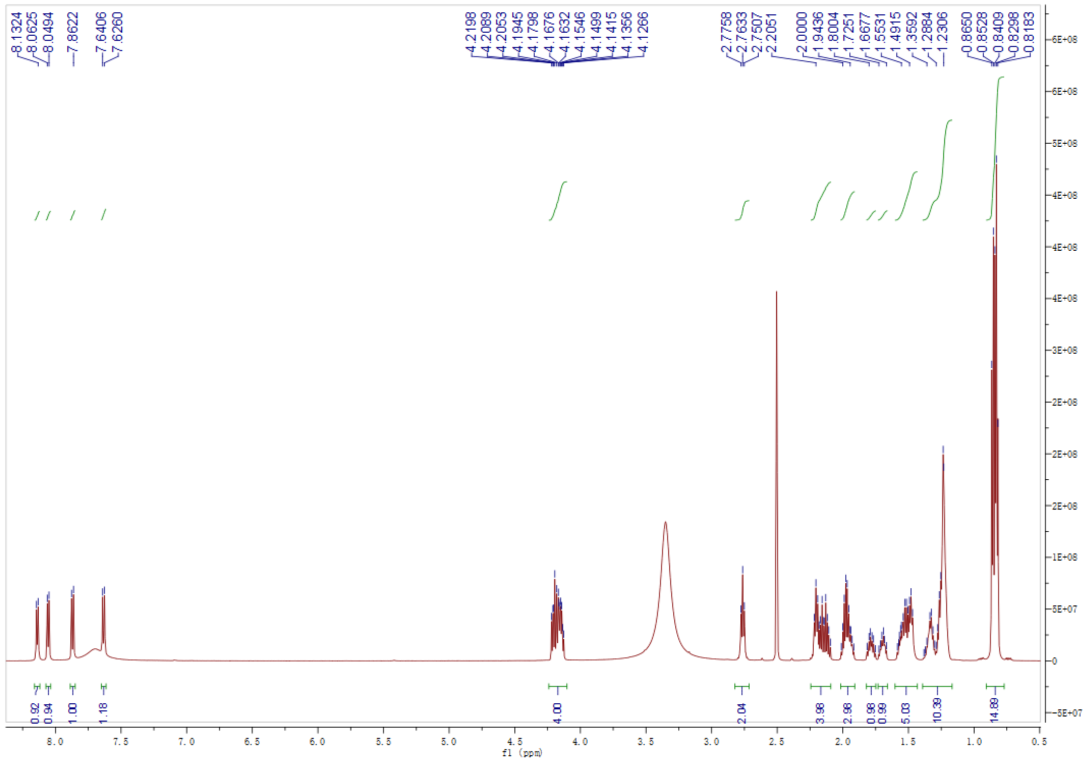

**Fig. S43.** $^{1}H$ NMR spectrum of compound **10** in DMSO-$d_6$.

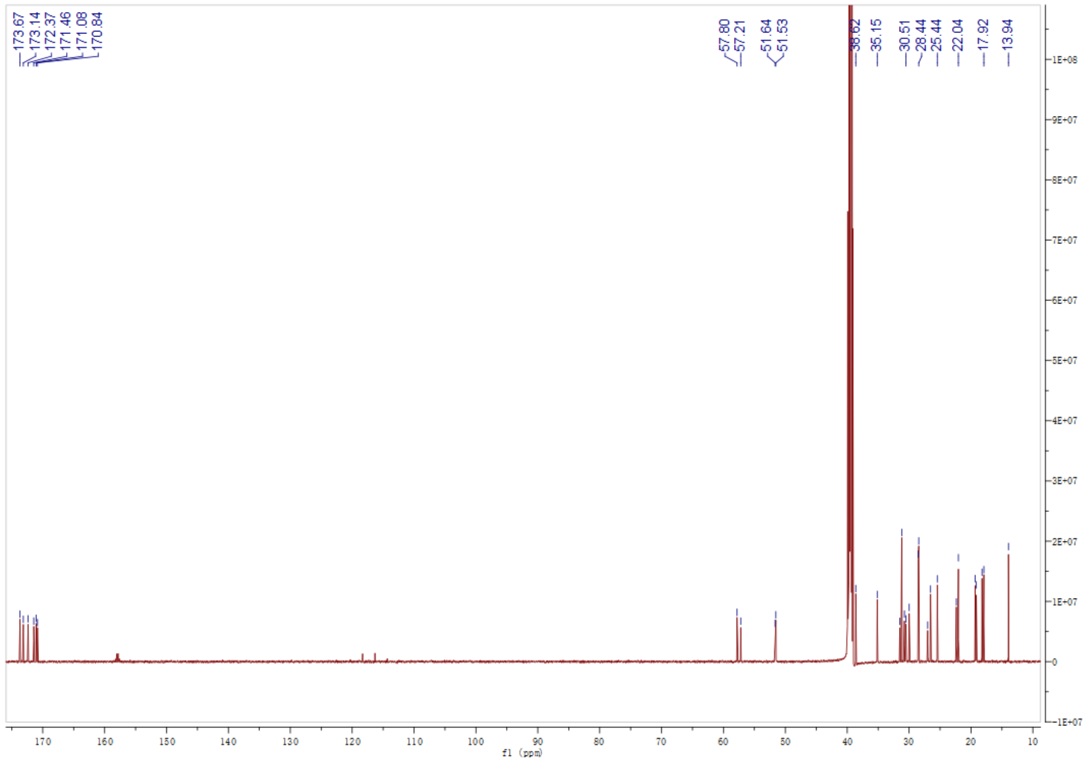

**Fig. S44.** $^{13}C$ NMR spectrum of compound **10** in DMSO-$d_6$.

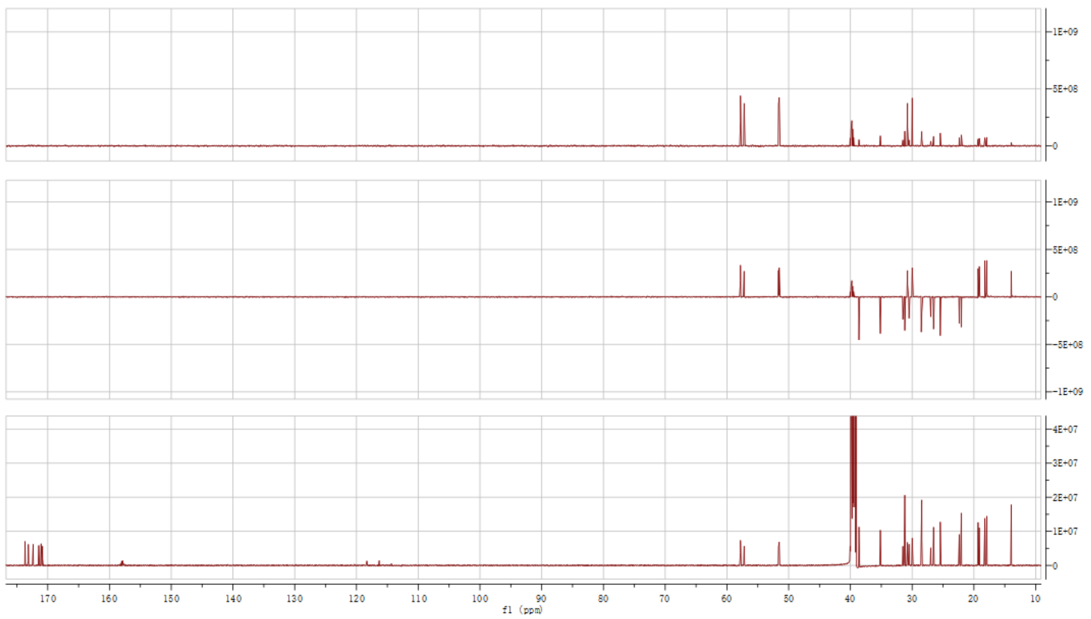

**Fig. S45.** DEPT spectrum of compound **10** in DMSO-$d_6$.

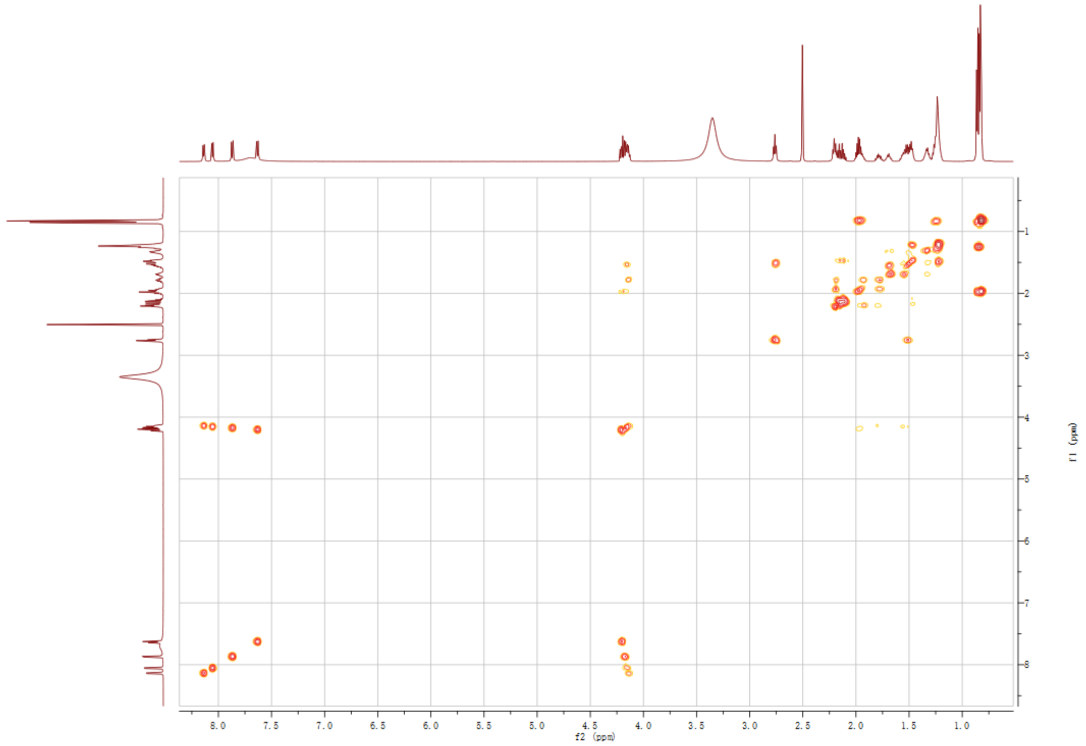

**Fig. S46.** $^1$H-$^1$H COSY NMR spectrum of compound **10** in DMSO-$d_6$.

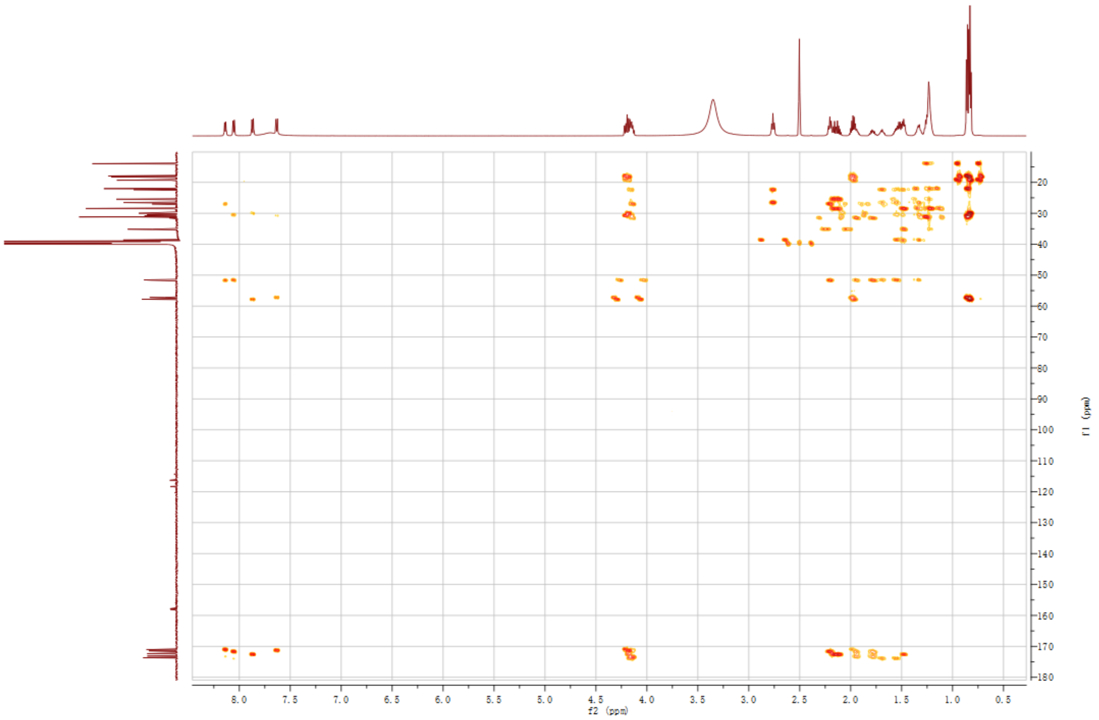

**Fig. S47.** HMBC spectrum of compound **10** in DMSO-$d_6$.

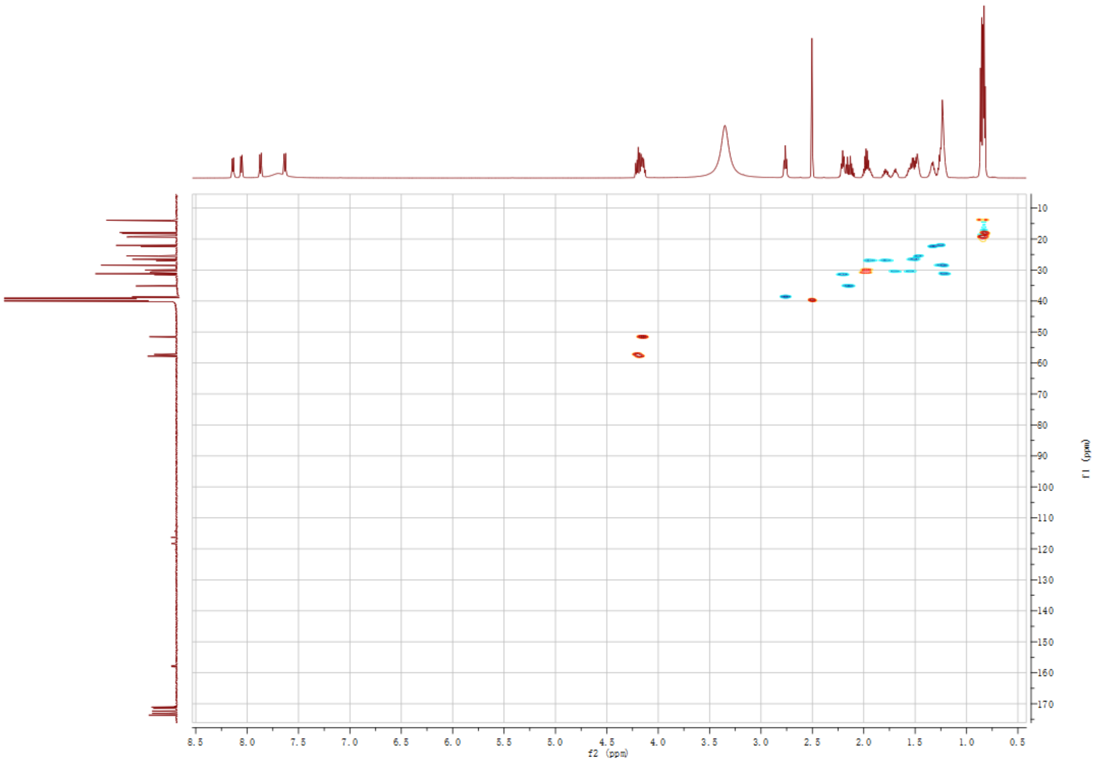

**Fig. S48.** HSQC spectrum of compound **10** in DMSO-$d_6$.

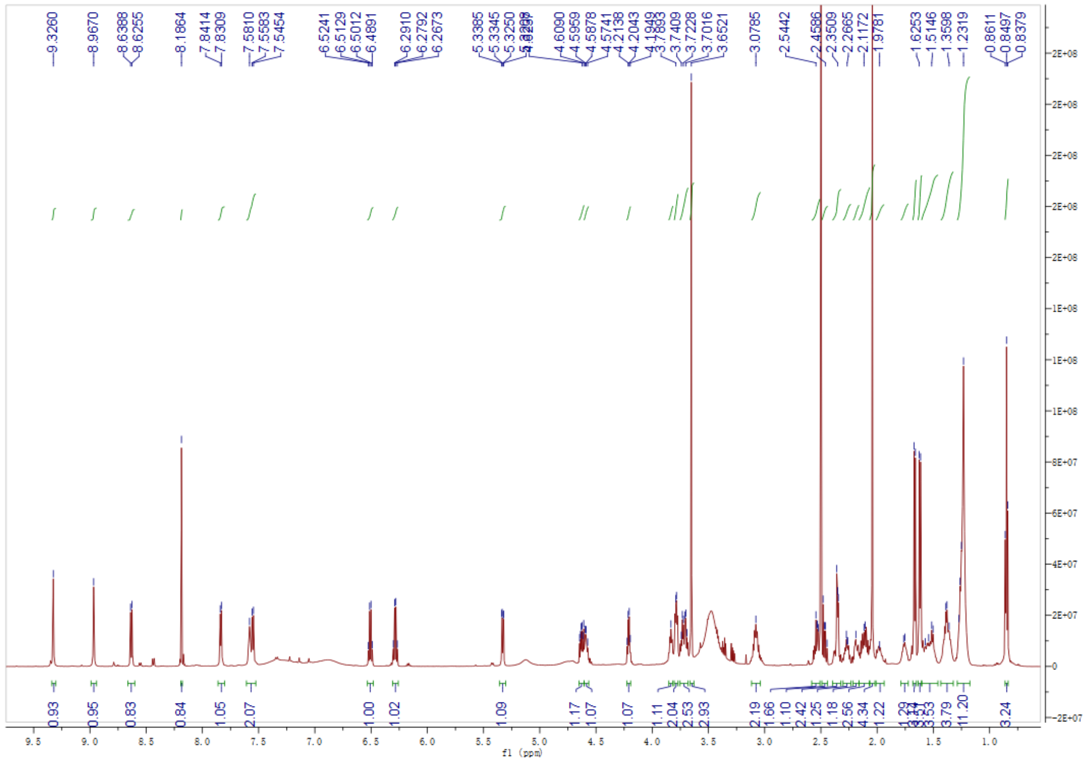

**Fig. S49.** $^{1}H$ NMR spectrum of compound **13** in DMSO-$d_6$.

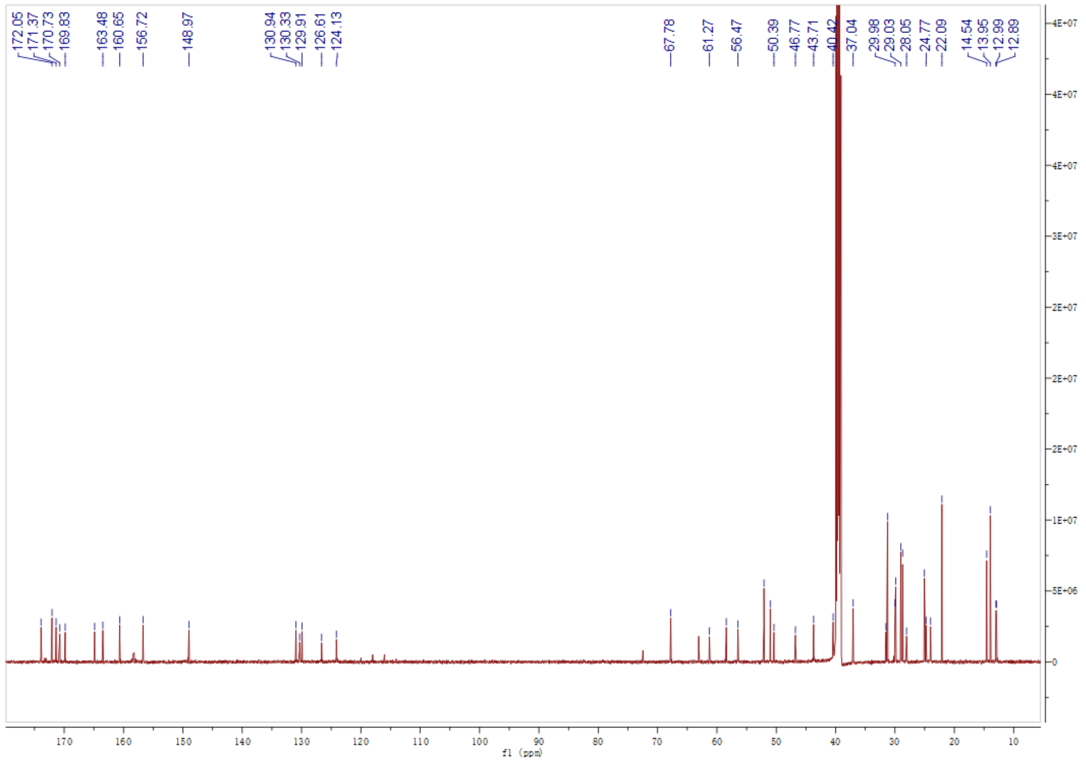

**Fig. S50.** $^{13}C$ NMR spectrum of compound **13** in DMSO-$d_6$.

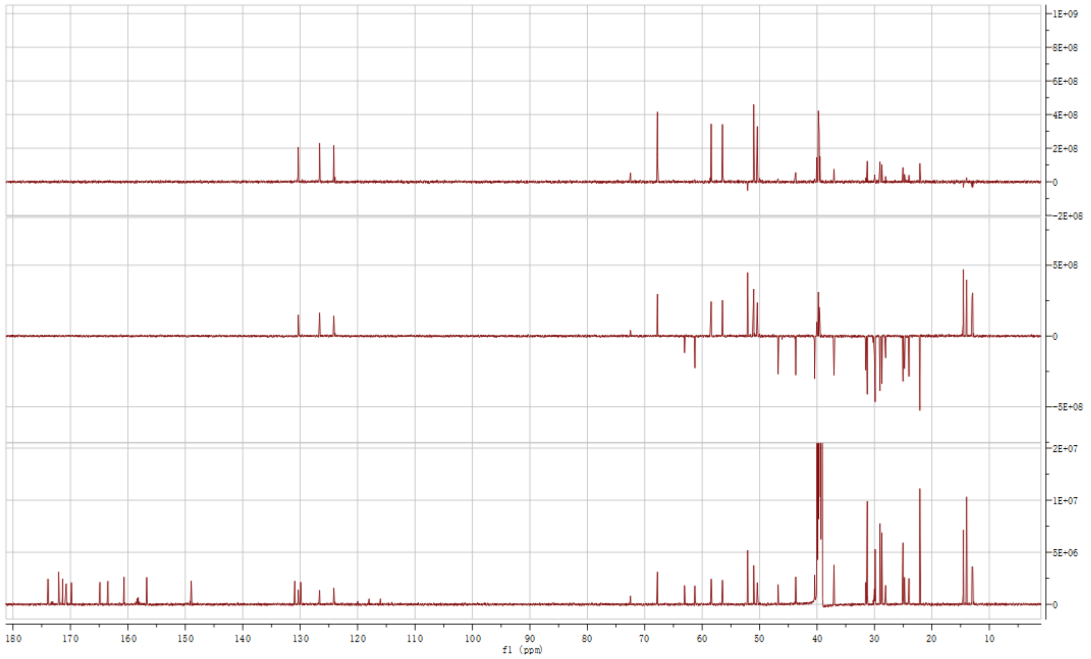

**Fig. S51.** DEPT spectrum of compound **13** in DMSO-$d_6$.

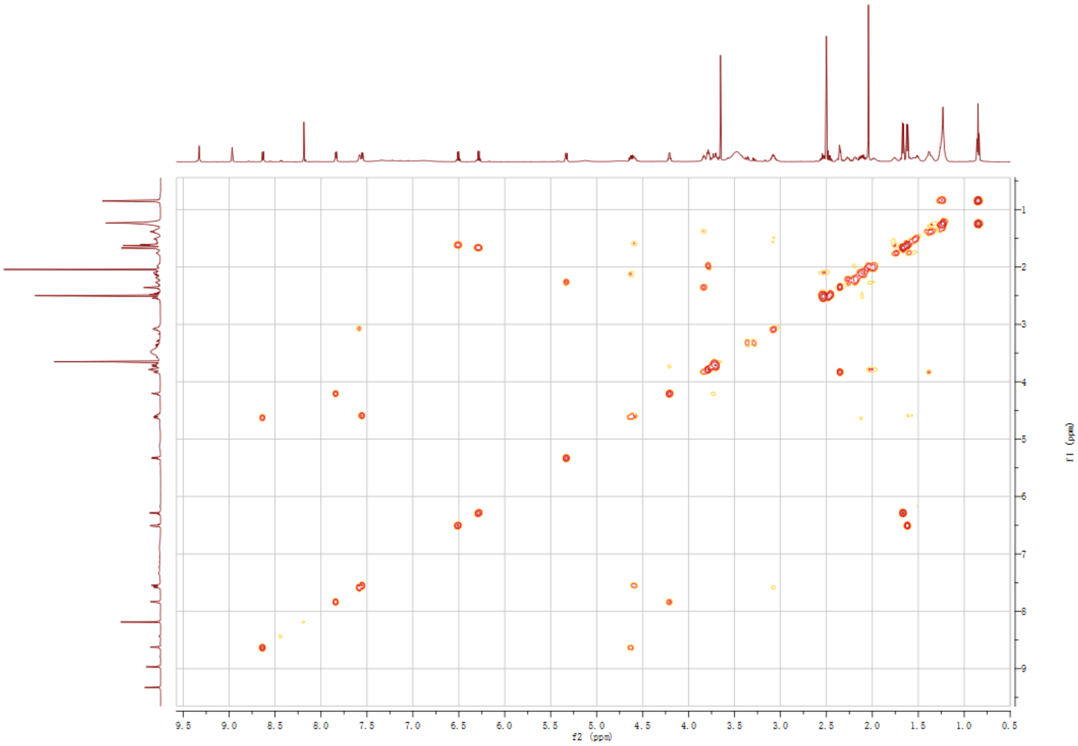

**Fig. S52.** $^1$H-$^1$H COSY NMR spectrum of compound **13** in DMSO-$d_6$.

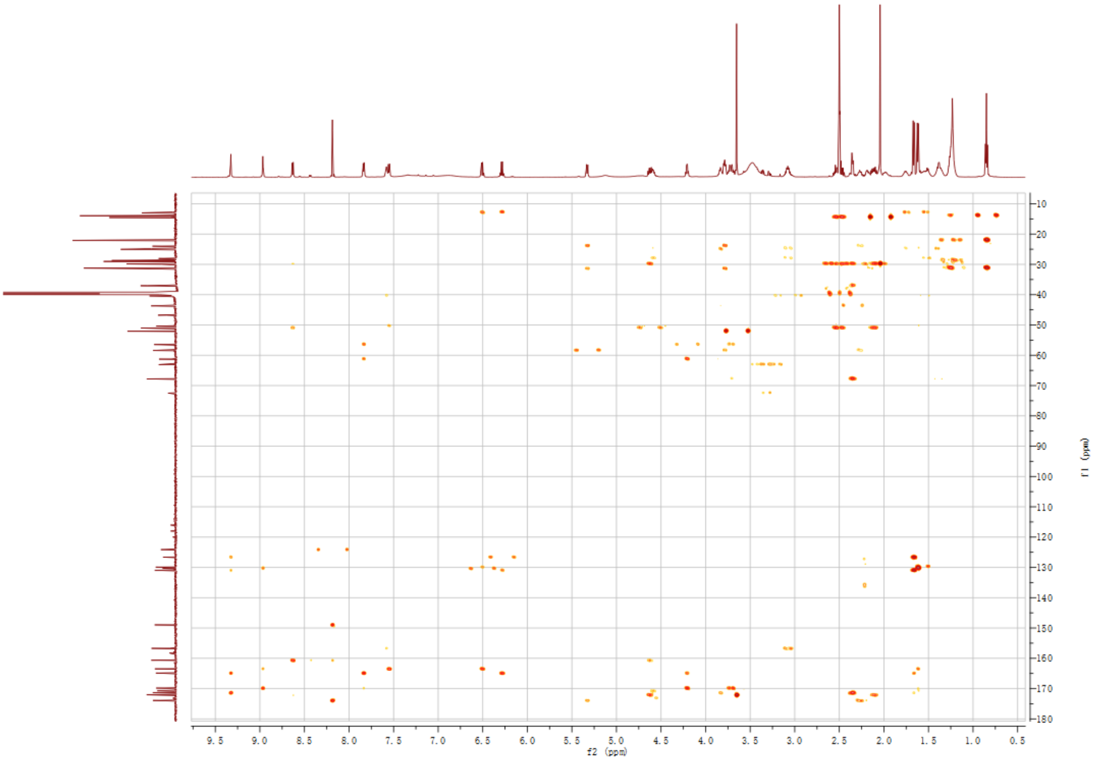

**Fig. S53.** HMBC spectrum of compound **13** in DMSO-$d_6$.

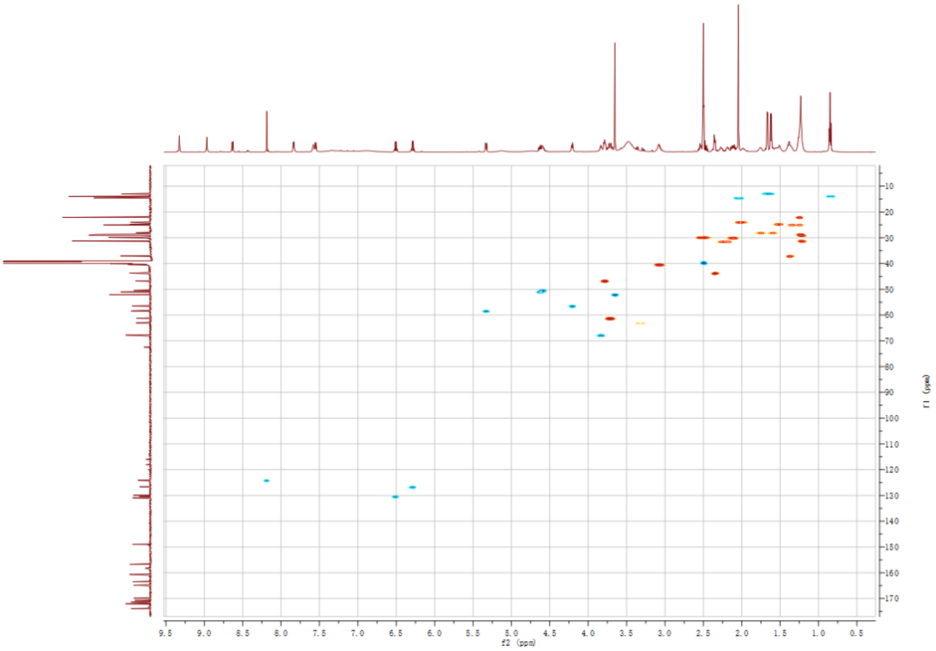

**Fig. S54.** HSQC spectrum of compound **13** in DMSO-$d_6$.

**Reference**

1. J. Fu, *et al.*, Full-length RecE enhances linear-linear homologous recombination and facilitates direct cloning for bioprospecting. *Nat Biotechnol* **30**, 440–446 (2012).

2. H. Chen, *et al.*, Identification of Holrhizins E–Q Reveals the Diversity of Nonribosomal Lipopeptides in *Paraburkholderia rhizoxinica*. *J. Nat. Prod.* **83**, 537–541 (2020).

3. J. Liu, *et al.*, Rational construction of genome-reduced Burkholderiales chassis facilitates efficient heterologous production of natural products from proteobacteria. *Nat Commun* **12**, 4347 (2021).

4. X. Bai, *et al.*, Heterologous Biosynthesis of Complex Bacterial Natural Products in *Burkholderia gladioli*. *ACS Synth. Biol.* **12**, 3072–3081 (2023).

5. X. Bian, *et al.*, Direct Cloning, Genetic Engineering, and Heterologous Expression of the Syringolin Biosynthetic Gene Cluster in *E. coli* through Red/ET Recombineering. *ChemBioChem* **13**, 1946–1952 (2012).

6. X. Wang, *et al.*, Discovery of recombinases enables genome mining of cryptic biosynthetic gene clusters in Burkholderiales species. *Proc. Natl. Acad. Sci. U.S.A.* **115**, E4255-E4263 (2018).

7. K. Blin, *et al.*, antiSMASH 7.0: new and improved predictions for detection, regulation, chemical structures and visualisation. *Nucleic Acids Research* **51**, W46–W50 (2023).

8. R. He, *et al.*, Knowledge-guided data mining on the standardized architecture of NRPS: Subtypes, novel motifs, and sequence entanglements. *PLoS Comput Biol* **19**, e1011100 (2023).

9. K. Tamura, G. Stecher, S. Kumar, MEGA11: Molecular Evolutionary Genetics Analysis Version 11. *Mol Biol Evol* **38**, 3022–3027 (2021).

10. L. Zhong, *et al.*, Engineering and elucidation of the lipoinitiation process in nonribosomal peptide biosynthesis. *Nat Commun* **12**, 296 (2021).

11. H. Wang, *et al.*, ExoCET: exonuclease in vitro assembly combined with RecET recombination for highly efficient direct DNA cloning from complex genomes. *Nucleic Acids Res* **46**, e28–e28 (2018).

12. H. Wang, *et al.*, RecET direct cloning and Redαβ recombineering of biosynthetic gene clusters, large operons or single genes for heterologous expression. *Nat Protoc* **11**, 1175–1190 (2016).

13. Q. Ouyang, *et al.*, Promoter Screening Facilitates Heterologous Production of Complex Secondary Metabolites in Burkholderiales Strains. *ACS Synth. Biol.* **9**, 457–460 (2020).

14. H. Wang, *et al.*, Improved seamless mutagenesis by recombineering using ccdB for

counterselection. *Nucleic Acids Res* **42**, e37–e37 (2014).

15. K. Fujii, *et al.*, A Nonempirical Method Using LC/MS for Determination of the Absolute Configuration of Constituent Amino Acids in a Peptide: Elucidation of Limitations of Marfey's Method and of Its Separation Mechanism. *Anal. Chem.* **69**, 3346–3352 (1997).

16. R. Li, *et al.*, Development and application of an efficient recombineering system for *Burkholderia glumae* and *Burkholderia plantarii*. *Microbial Biotechnology* **14**, 1809–1826 (2021).

17. J. Fu, M. Teucher, K. Anastassiadis, W. Skarnes, A. F. Stewart, A Recombineering Pipeline to Make Conditional Targeting Constructs. *Methods Enzymol* **477**, 125-144 (2010).

18. C. Dehio, M. Meyer, Maintenance of broad-host-range incompatibility group P and group Q plasmids and transposition of Tn5 in Bartonella henselae following conjugal plasmid transfer from Escherichia coli. *J Bacteriol* **179**, 538–540 (1997).

19. H. Chen, *et al.*, Genomics-Driven Activation of Silent Biosynthetic Gene Clusters in Burkholderia gladioli by Screening Recombineering System. *Molecules* **26**, 700 (2021).

20. J. Rüschenbaum, W. Steinchen, F. Mayerthaler, A. Feldberg, H. D. Mootz, FRET Monitoring of a Nonribosomal Peptide Synthetase Elongation Module Reveals Carrier Protein Shuttling between Catalytic Domains. *Angew Chem Int Ed* **61**, e202212994 (2022).
